# Computational prediction of protein subdomain stability in *MYBPC3* enables clinical risk stratification in hypertrophic cardiomyopathy and enhances variant interpretation

**DOI:** 10.1101/2020.11.29.402974

**Authors:** Andrea D. Thompson, Adam S. Helms, Anamika Kannan, Jaime Yob, Neal K. Lakdawala, Samuel G. Wittekind, Alexandre C. Pereira, Daniel L. Jacoby, Steven D. Colan, Euan A. Ashley, Sara Saberi, James S. Ware, Jodie Ingles, Christopher Semsarian, Michelle Michels, Francesco Mazzarotto, Iacopo Olivotto, Carolyn Y. Ho, Sharlene M. Day, SHaRe investigators

## Abstract

**Purpose:** Variants in *MYBPC3* causing loss-of-function are the most common cause of HCM. However, a substantial number of patients carry missense variants of uncertain significance (VUS) in *MYBPC3.* We hypothesize that a structural-based algorithm, STRUM, which estimates the effect of missense variants on protein folding, will improve clinical risk stratification of patients with HCM and a *MYBPC3* VUS.

**Methods:** Among 7,963 patients in the multi-center Sarcomeric Human Cardiomyopathy Registry, 120 unique missense VUSs in *MYBPC3* were identified. Variants were evaluated for their effect on subdomain folding and a stratified time-to-event analysis for an overall composite endpoint (first occurrence of ventricular arrhythmia, heart failure, all-cause mortality, atrial fibrillation, and stroke) was performed for patients with HCM and a *MYBPC3* missense VUS.

**Results:** We demonstrated that patients carrying a *MYBPC3* VUS predicted to cause subdomain misfolding (STRUM +, ΔΔG ≤-1.2 kcal/mol) exhibited a higher rate of adverse events compared to those with a STRUM-VUS (Hazard Ratio=2.29, P=0.0282). *In silico* saturation mutagenesis of *MYBPC3* identified 4,943/23,427 (21%) missense variants that were predicted to cause subdomain misfolding.

**Conclusions:** STRUM enables clinical risk stratification of patients with HCM and a *MYBPC3* VUS and has the capacity to improve prognostic predictions and clinical decision making.

## Introduction

Genetic variant interpretation is an ongoing challenge in clinical medicine, particularly when the gene of interest lacks robust functional assays^1, 2^. A variety of computational algorithms have been developed to predict variant pathogenicity, but their sensitivity and specificity are often poor, particularly when applied broadly across different diseases and different genes^1, 3^. Loss-of-function pathogenic variants are common^1, 4, 5^, resulting from either frameshift or nonsense variants creating a premature stop codon, splice errors, disruption of enzymatic activity, alteration of protein-protein interactions, or protein misfolding^1, 6, 7^. Recognizing a common mechanism by which variants in a particular gene lead to loss-of-function can inform the development of gene-specific computational algorithms to more accurately predict pathogenicity among variants that cannot be confidently classified based on clinical and family data alone^6, 7^.

Herein we focus on *MYBPC3* (encoding the protein, cardiac myosin binding protein C, or MyBP-C). Pathogenic variants in *MYBPC3* account for ∼50% of patients with sarcomeric HCM^8, 9^, and are inherited in an autosomal dominant fashion (OMIM 115197). Hypertrophic cardiomyopathy (HCM) is characterized by left ventricular hypertrophy and fibrosis. Patients with HCM can experience a variety of adverse clinical outcomes, including outflow tract obstruction, arrhythmias, heart failure, and sudden cardiac death^8^. Genetic variants in *MYBPC3* consist of both truncating and non-truncating types. Rarely found in healthy populations, truncating *MYBPC3* variants result in a premature stop codon and cause HCM through complete loss-of-function and haploinsufficiency at the transcript and protein level^10–13^. Thus, interpretation of these truncating variants as pathogenic is straightforward^14^.

However, the interpretation of missense variants within *MYBPC3* presents a major challenge. Single amino acid substitutions (missense variants), are found commonly in healthy populations. Further, since missense variants do not disrupt the reading frame, protein function may be tolerant to these minor sequence changes. Thus, many missense variants lack sufficient evidence to be classified as either pathogenic or benign and are classified as variants of uncertain significance (VUS)^14, 15^. While identifying pathogenic variants allows for predictive genetic testing in at-risk relatives^16^, a VUS is not clinically actionable and may lead to misinterpretation by clinicians and patients^17^.

Identification of a pathogenic sarcomere genetic variant for HCM also has important prognostic implications. Patients with HCM and a pathogenic sarcomere variant (sarcomeric HCM) have a higher risk of adverse clinical outcomes compared to those without a sarcomere gene variant (non-sarcomeric HCM)^8^. Patients carrying a sarcomere gene VUS, on average, exhibit an intermediate risk of adverse events^8^, most likely because VUSs represent a mixed pool of pathogenic and benign variants that cannot be parsed on the basis of clinical and genetic data alone.

Because loss-of-function is an established mechanism for pathogenic variants in *MYBPC3*, we hypothesized that applying a computational approach, called STRUM^18–20^, that incorporates both sequence-based and structure-based algorithms to missense *MYBPC3* VUSs will identify those variants that result in protein subdomain misfolding (STRUM+), thereby supporting pathogenicity and improving variant interpretation. We further predict that this approach will improve clinical risk stratification by identifying a subpopulation of patients with HCM and a STRUM+ *MYBPC3* missense VUS who are at risk for adverse clinical outcomes, at a frequency similar to patients with HCM carrying known pathogenic variants

## Methods and Materials

### SHaRe Registry Data Extraction and *MYBPC3* Variant Classification

The generation of the centralized SHaRe database has been previously described^8^. Data for this study were exported from quarter 1 of 2019. Inclusion criteria included a site-designated diagnosis of HCM using standard diagnostic criteria^8^. SHaRe non-truncating *MYBPC3* missense variants (Tables S1,S2) were classified as previously reported^14^ in accordance with American College of Medical Genetics and Genomics (ACMGG) guidelines, leveraging available clinical and experimental data^9, 14, 21–23^. Since variants in *MYBPC3* present in gnomAD with allele frequencies of > 4E-05 and absent in SHaRe are unlikely to be independently pathogenic for HCM, these variants were included in our list of benign *MYBPC3* variants^14^. More details regarding variant interpretation is provided within supplemental materials.

It has previously been shown that patients carrying pathogenic non-truncating exhibit similar clinical outcomes to those carrying truncating *MYBPC3* variants^14^. Thus, a reference population including previously adjudicated truncating and non-truncating *MYBPC3* pathogenic/likely pathogenic (pathogenic) variants [*MYBPC3*-path-all] was used. A second reference population included patients with HCM who underwent genetic testing and were negative for sarcomere variants Sarc-^8^.

### Computational Structural and Protein Folding Stability Predictive Modeling

MyBP-C is made up of immunoglobulin and fibronectin subdomains (C0-C10) [NM_000256.3, NP_000247.2]. For *MYBPC3* missense variants we utilized STRUM to calculate the effect of the missense variant on the Gibbs free energy of local subdomain folding (referred to as the ΔΔG)^18^ (Table S3). The STRUM algorithm utilizes I-TASSER subdomain modeling while calculating ΔΔG. I-TASSER (Iterative Threading ASSEmbly Refinement) is a hierarchical approach to protein structure and function prediction^25–27^. A negative ΔΔG value indicates the degree of reduced folding energy (kcal/mol) relative to the wild-type subdomain, or folding destabilization^18^. Previous experimental validation of this algorithm compared STRUM predictions to 3,421 experimentally tested variants from 150 proteins and demonstrated a Pearson’s correlation coefficient of 0.79 and root mean square error of prediction of 1.2 kcal/mol^18^. Thus, a value of ΔΔG ≤-1.2 kcal/mol was defined as the cut-off for destabilizing (deleterious) variants. Further details regarding STRUM analysis and structural models are provided within the Supplemental Materials (Figure S1-S3, Table S3).

### Computational Sequence Based Variant Analysis (PolyPhen-2, SIFT, CardioBoost)

We compared the STRUM prediction for *MYBPC3* missense variants with a sequence-based algorithm embedded in STRUM (SIFT)^28, 29^. We also analyzed these variants with PolyPhen-2 (HumVar database), a sequence based algorithm which incorporates basic structural information such as b-factor into its algorithm^30^. Finally, we compared our result to those obtained using CardioBoost which is a recently published disease specific machine learning classifier to predict pathogenicity of rare (gnomAD Allele Frequency < 0.1%) missense variants in genes associated with cardiomyopathies and arrhythmias^6^.

### Clinical Outcomes Analysis

Only patients with HCM carrying a single *MYBPC3* missense VUS were included in clinical outcomes analyses to avoid confounding from cases with multiple gene variants^31^. Comparisons using time-to-event analysis were made between variants predicted to be deleterious and those predicted to be non-deleterious by computational algorithms. The primary outcome was an overall composite of HCM patients carrying a single MYBPC3 missense VUS predicted to cause subdomain misfolding by STRUM compared to HCM patients carrying a single MYBPC3 missense VUS not predicted to cause subdomain misfolding by STRUM. The overall composite was previously defined as the first occurrence of any component of the ventricular arrhythmia composite, heart failure composite (without inclusions of LV ejection fraction), all-cause mortality, atrial fibrillation (AF), or stroke^8^. Results were compared to reference populations *MYBPC3*-path-all and Sarc-. A secondary analysis of a heart failure composite, ventricular arrhythmia composite, and atrial fibrillation was also performed. Finally, a secondary analysis using alternative computational algorithms (SIFT, Polyphen-2, CardioBoost) was also performed. Composite outcomes are defined in more detail in the supplemental materials.

### Statistical Analysis

Data presented as mean ± standard deviation were analyzed by t-test for two groups or ANOVA for >2 groups with Tukey’s post hoc test for multiple comparisons. Data presented as frequency were analyzed by a chi-square test. Odds ratio (with 95% Confidence Interval), specificity, and sensitivity were calculated to evaluate the association between computational prediction algorithms and known pathogenic/likely pathogenic (pathogenic) or benign/likely benign (benign) variants (further details provided in supplemental materials). Primary and secondary clinical outcomes were analyzed by the Kaplan-Meier method from time of birth. Analysis from time of birth is appropriate given that the genetic variant is present from birth and variability in time to, and reason for, clinical presentation could confound the results if time from diagnosis were used. Patients who did not have the outcome of interest were censored at the time of their last recorded follow-up in SHaRe. Comparison between curves was performed using Log-rank Mantel-Cox test with p-values of < 0.05 considered statistically significant. Median event free survival and hazard ratio (mantel-Haenszel) are also reported. Statistical analyses were performed using GraphPad Prism software (San Diego, CA).

## Results

### Patients with HCM and a *MYBPC3* missense VUS predicted to disrupt subdomain folding (STRUM+) exhibit a higher incidence of adverse clinical outcomes

We began by evaluating all *MYBPC3* missense VUS within SHaRe using STRUM. *MYBPC3* VUSs exhibited a mean ΔΔG of -0.73 +/- 1.06 kcal/mol (Figure S4). Of 120 unique *MYBPC3* missense VUSs, 34 (28%) were predicted to cause domain misfolding with ΔΔG values ≤1.2 kcal/mol (deleterious) (Table S2). Next, we evaluated clinical characteristics and outcomes in patients with HCM and a single missense *MYBPC3* VUS predicted to disrupt subdomain folding (STRUM+) compared to patients carrying a VUS not predicted to disrupt subdomain folding (STRUM-). For this analysis, we included only patients who carried a single VUS within *MYBPC3*, and excluded patients who carried a second pathogenic variant or variant of uncertain significance. Of the 181 patients carrying a *MYBPC3* missense VUS denoted in Table S2, 105 patients carried a single *MYBPC3* missense VUS while the remaining 76 patients carried an additional VUS or pathogenic variant. Patients with a STRUM+ vs STRUM-*MYBPC3* VUS exhibited similar clinical characteristics including BMI, gender, ancestry, age at diagnosis, wall thickness, ejection fraction, left ventricular outflow tract obstruction (Table 1). We observed that patients carrying a STRUM+ VUS experienced higher rates of composite adverse events compared to patients carrying a STRUM-VUS (Figure 1, hazard ratio 2.3, p = 0.03). Furthermore, patients carrying a STRUM+ VUS exhibited a similar rate of adverse clinical events compared to patients carrying a pathogenic variant (*MYBPC3*-Path-all). Conversely, patients carrying STRUM-VUSs exhibited a lower frequency of outcomes, similar to Sarc-patients (Figure 2). There were no statistically significant differences between groups for the individual component outcomes, including ventricular arrhythmias, heart failure, or atrial fibrillation (Figure S5).

**Figure 1:**
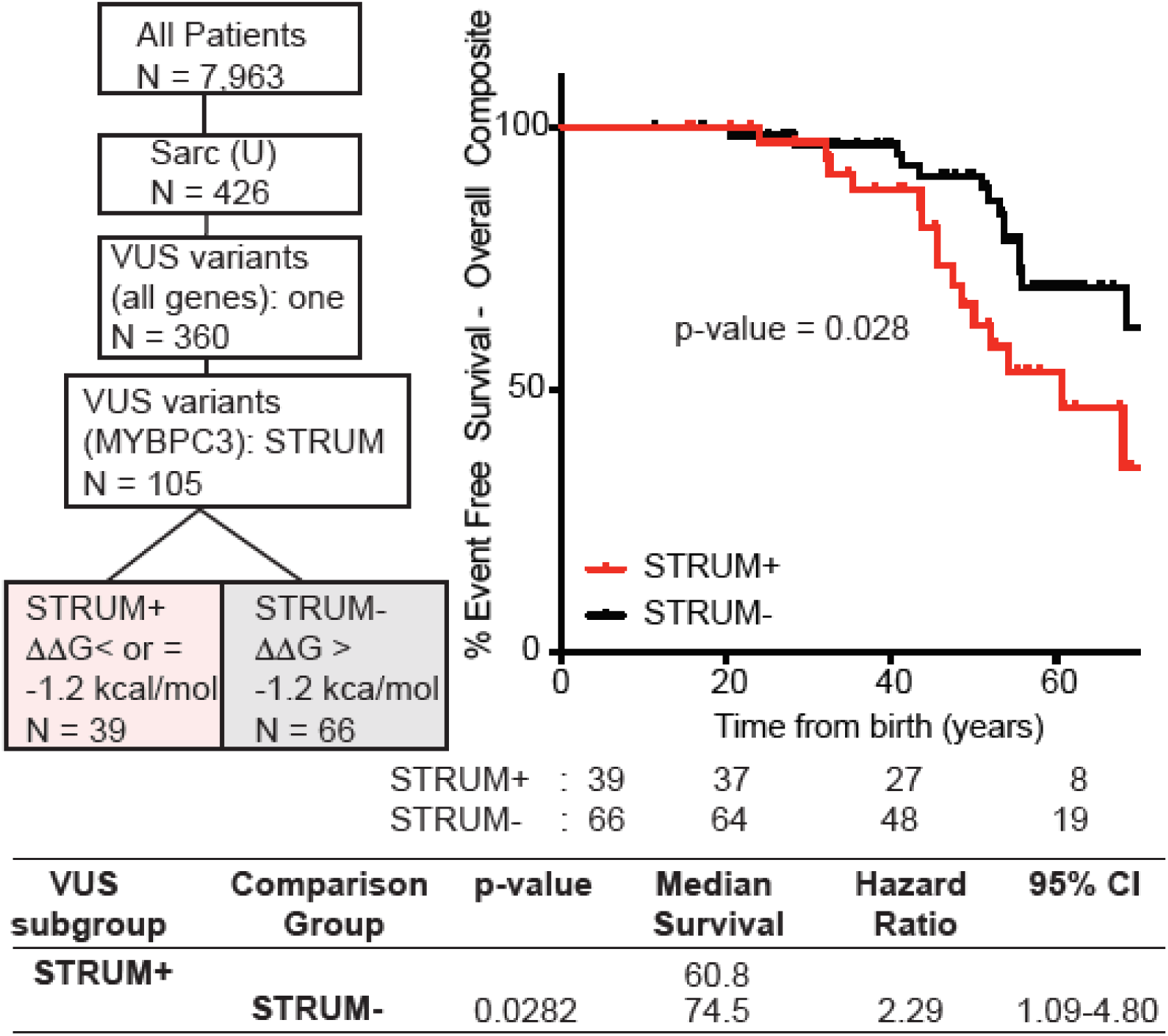
Patients with a *MYBPC3* VUS identified as deleterious by STRUM (STRUM +) are associated with an increased risk for adverse HCM-related outcomes. Selection within ShaRe of patients with HCM carrying a single MYBPC3 missense VUS is shown on the left. 105 patients carry a single MYBPC3 *missense* VUS, covering 77 distinct MYBPC3 VUSs. Kaplan Meier curves, median event free survival (years), and hazard ratio with corresponding 95% CI reveal that patients carrying a STRUM+ *MYBPC3* VUS (red) exhibited a higher rate of adverse HCM-related outcomes (Overall Composite) compared to patients carrying a STRUM-variant (black).

**Figure 2:**
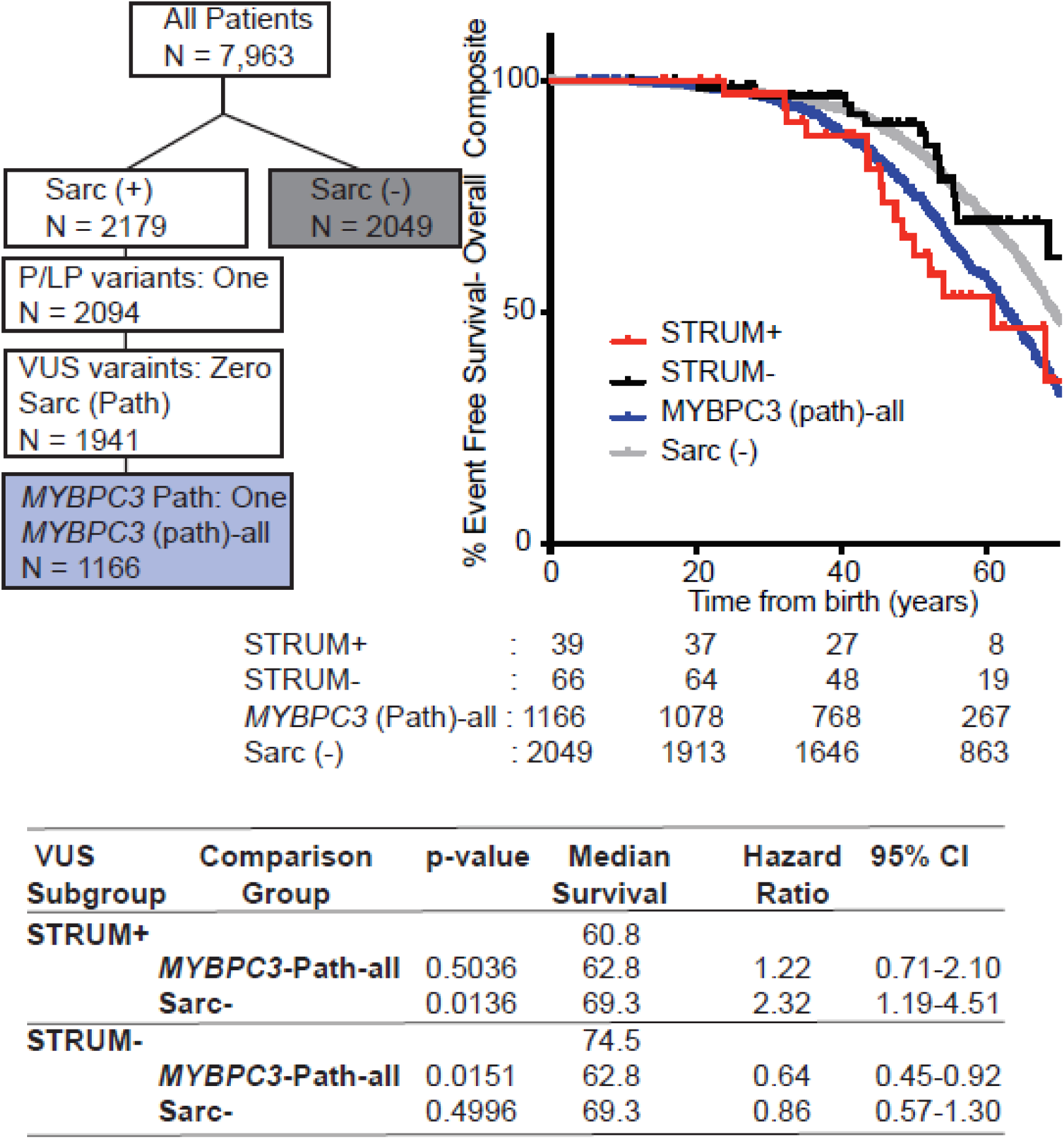
Patients with a *MYBPC3* VUS identified as deleterious by STRUM (STRUM +) exhibit clinical outcomes similar to patients with a *MYBPC3* pathogenic variants. Selection within SHaRe of patients with HCM and a single *MYBPC3* pathogenic variant (*MYBPC3* -Path-all) and patients with HCM without a sarcomere gene variant after clinical genotype analysis (Sarc -) is shown on the left. Kaplan Meier curves, median event free survival (median survival), and hazard ratio with corresponding 95% confidence interval (CI) reveal patients carrying a STRUM+ *MYBPC3* VUS (red) exhibited overall composite outcomes similar to *MYBPC3*-Path-all patients (blue, p-value 0.5036). Whereas, patients carrying a STRUM-variant (black) exhibited a lower rate of adverse HCM-related outcomes (overall Composite) similar to Sarc-patients (grey).

**Table 1:** Demographics of patients with HCM and single *MYBPC3* VUS

|  | STRUM (+)<br>n = 39 (37%) | STRUM (-)<br>n = 66 (63%) | p-value<br>STRUM (+)<br>vs. STRUM (-) |
| --- | --- | --- | --- |
| **Baseline Characteristics** |  |  |  |
| Female, n (%) | 10 (26%) | 14 (21%) | 0.6308 |
| Age at diagnosis, mean (STD), y | 40.31 (17.13) | 41.34 (21.17) | 0.9930 |
| Follow-up time, mean (STD), y | 9.82 (10.00) | 12.18 (11.83) | 0.7575 |
| Maximum BMI, mean (STD) kg/m2 | 28.87 ( 4.10) | 27.96 ( 6.03) | 0.9390 |
| Race, n % |  |  | 0.5107 |
| White | 35 (90%) | 60 (91%) |  |
| Black | 2 ( 5%) | 1 ( 2%) |  |
| Other/not reported | 2 ( 5%) | 5 ( 8%) |  |
| Proband, n (%) | 35 (90%) | 64 (97%) | 0.1232 |
| Family History HCM, n (%) | 13 (33%) | 19 (29%) | 0.6249 |
| Family History SCD n, (%) | 8 (21%) | 9 (14%) | 0.3553 |
| **Echocardiogram Data** |  |  |  |
| Maximal LVWT, mean (STD), mm | 22.73 ( 6.97) | 20.83 ( 7.29) | 0.4287 |
| Minimum LVEF, mean (STD), % | 59.88 ( 8.32) | 60.12 (11.34) | 0.9996 |
| LVOT peak gradient > or = 30 mmHg; n (%) | 9 (23%) | 21 (32%) | 0.3380 |

BMI = body mass index, LVWT = left ventricular wall thickness, LVEF = left ventricular ejection fraction, LVOT = left ventricular outflow tract

### STRUM exhibits improved specificity over established sequence-based prediction algorithms and improved sensitivity when combined with CardioBoost

To determine the sensitivity and specificity of STRUM to differentiate pathogenic from benign variants within *MYBPC3* we performed STRUM analysis on all known pathogenic missense variants within ShaRe (n = 19) and known missense benign variants within SHaRe and gnomAD (n =110, Table S1, Figure 3A). These variants were present in 412 patients with HCM within the SHaRe registry. *MYBPC3* benign variants exhibited a mean ΔΔG of -0.31 +/- 0.60 kcal/mol which was significantly higher than *MYBPC3* VUS (ΔΔG of -0.73 +/- 1.06 kcal/mol, p = 0.005) (Figure S4) and *MYBPC3* pathogenic variants (mean ΔΔG of -1.00 +/- 1.08 kcal/mol, p = 0.016) (Figure 3A). We found that variants predicted to be deleterious by STRUM were more likely to be pathogenic variants (OR 5.9, 95% CI 1.8-19.6) (Figure 3C). Algorithms that were purely sequence-based achieved greater sensitivity but performed inferiorly to STRUM in regard to specificity. STRUM exhibited a 93% specificity for benign variants and PolyPhen-2 and SIFT exhibited a specificity of 62% (OR 4.5, 95% 1.5-13.5) and 54% (OR 1.3, 95% CI 0.5-3.4) respectively (Figure 3C, Figure S6). Additionally, variant interpretation by SIFT or PolyPhen-2 did not stratify patients carrying a *MYBPC3* VUS for clinical adverse outcomes (Figure S6).

**Figure 3.**
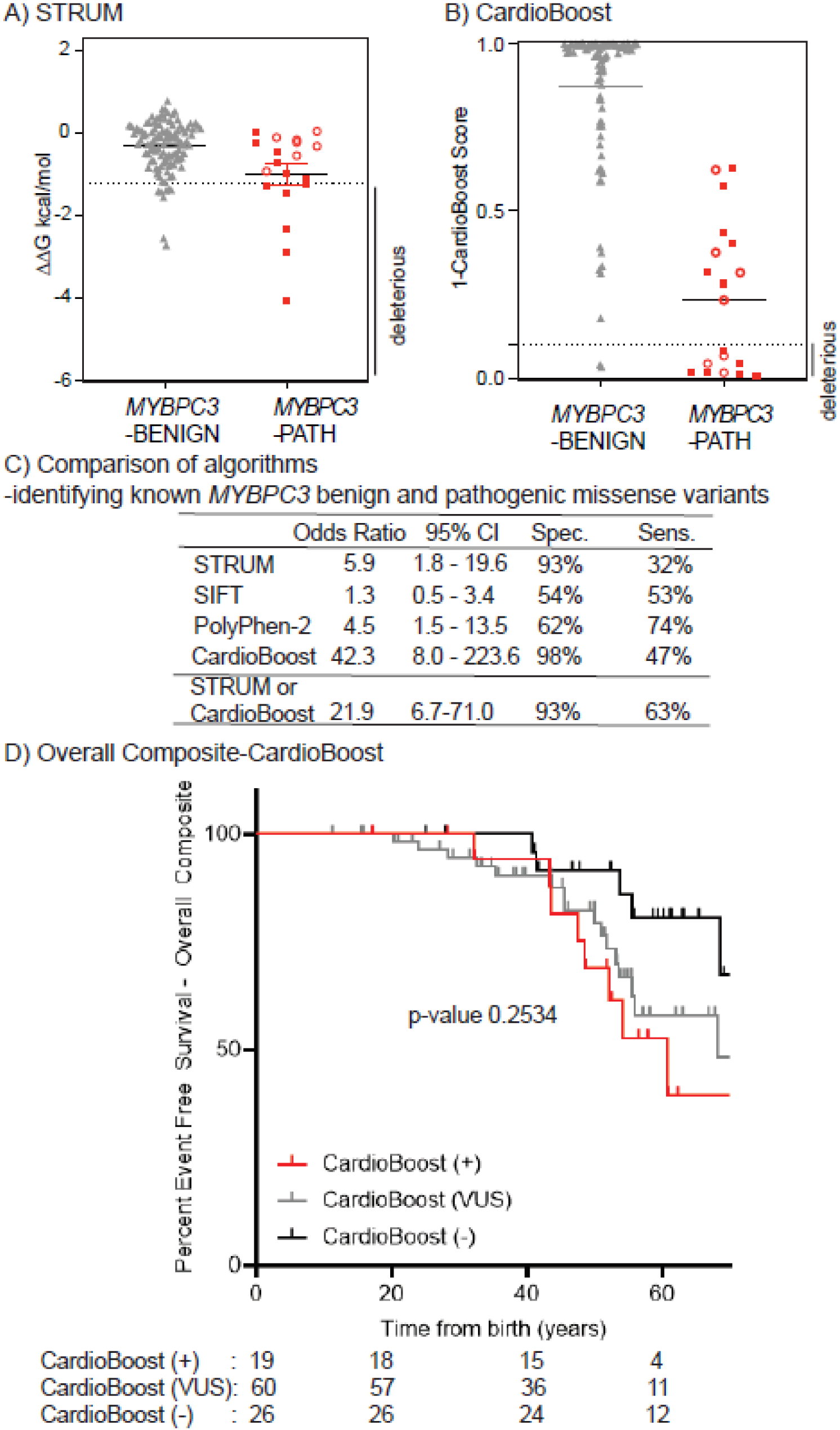
STRUM is complementary to CardioBoost. (A) STRUM: Results of computational analysis for each unique *MYBPC3*-Benign (grey triangles, n= 110) and *MYBPC3*-Path (red circles, n =19) variant are shown. Mean and SEM for each group depicted. The cut-off for deleterious variants for STRUM was ΔΔG ≤-1.2 kcal/mol, (B) CardioBoost: Results of computational analysis for each unique MYBPC3- Benign (grey triangles, n = 96, 14 not amenable to CardioBoost analysis) and *MYBPC3*-Path (red circles, n = 19) variant are shown. The cut-off for deleterious variants for CardioBoost (CardioBoost+) was a probability score > 0.90 this is graphed as 1-CardioBoostScore <0.10. C3 pathogenic variants are depicted in open circles in panel A and B. (C) Statistical Analysis of computational method utilized herein STRUM (Figure 3), CardioBoost (Figure 3), SIFT (Figure S6), and Polyphen-2 (Figure S6) is shown including Odd’s ratio, 95% confidence interval (CI), Sensitivity, and Specificity. (D) Using the same patient selection criteria in SHaRe detailed in Figure 1, patients with HCM and a MYBPC3 missense VUS were analyzed by CardioBoost. CardioBoost (+) was a probability score > 0.90, CardioBoost VUS ≥0.10 and ≤ 0.90, and CardioBoost (-) < 0.10. Of the 105 patients analyzed by STRUM 19 were CardioBoost(+). (F) Kaplan Meier curves reveal that patients carrying a CardioBoost (+) *MYBPC3* VUS (red) exhibited higher rates of adverse HCM-related outcomes (Overall Composite), than patients carrying a CardioBoost (-) MYBPC3 VUS (black), however the null hypothesis could not be excluded, p-value 0.0945. This remains true when comparing patients carrying a MYBPC3 VUS that is CardioBoost (+) red), CardioBoost (VUS) (grey), and CardioBoost (-) (black) (p-value 0.2534).

In comparison, the machine-learning based algorithm, CardioBoost (32), demonstrated a specificity of 98% for correctly classifying benign *MYBPC3* variants (OR 42.3, CI 8.0-223.6) (Figure 3, Table S1). For pathogenic variants, CardioBoost demonstrated a sensitivity of 47%. Interestingly, there was little overlap among known pathogenic variants predicted to be deleterious by STRUM and those predicted to be deleterious by CardioBoost, making the 2 algorithms complementary (Table S1). Combining these algorithms to classify any variant predicted to be deleterious by CardioBoost or STRUM as pathogenic, maintained a high specificity of 93% and improved sensitivity to 63% (Figure 3C).

When examining patients with HCM and a *MYBPC3* missense VUS, STRUM and CardioBoost exhibited good agreement with STRUM identifying a larger number of *MYBPC3* VUS as deleterious. Sixteen of the thirty-nine (41%) patients with a STRUM+ *MYBPC3* VUS were also identified as CardioBoost+. Further, only 3 additional patients with HCM and a *MYBPC3* VUS were uniquely identified as CardioBoost+ (Table S2). While there is a trend towards clinical risk stratification in patients with HCM and a *MYBPC3* VUS using CardioBoost, the null hypothesis could not be excluded (Figure 3D).

### STRUM predictions within pathogenic variants are consistent with experimental modeling

Prior experimental characterization of *MYBPC3* pathogenic missense variants within the C10 domain, Leu1238Pro and Asn1257Lys, demonstrated that these variants failed to localize to the sarcomere and were rapidly degraded within primary cardiomyocytes^14^. Consistent with these experimental findings, pathogenic C10 domain variants are uniformly predicted to destabilize protein folding (ΔΔG of -2.89 and -1.45 kcal/mol respectively) (Figure 4).

**Figure 4.**
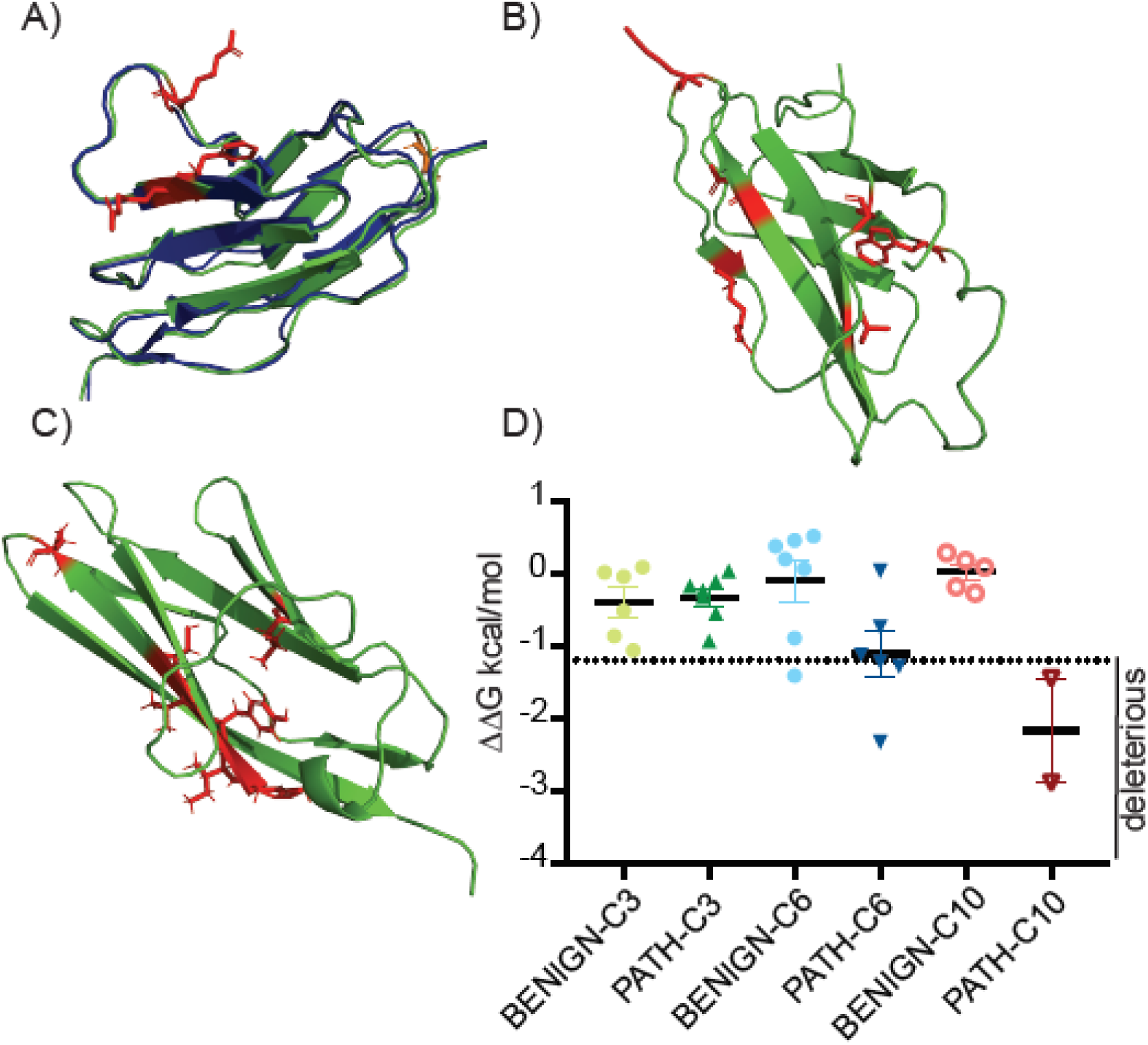
Structural Analysis of Pathogenic Missense *MYBPC3* variants. MyBP-C (the protein encoded by *MYBPC3*) domains C3, C6, and C10 were structurally modeled using I-TASSER (25, 26) (PyMOL,cartoon, green) Wild-type residues that are affected by missense pathogenic variants are depicted in red (PyMOL, sticks). (A) For the C3 domain, the I-TASSER model is aligned with an available NMR structure [2mq0.pdb(32), blue, PyMOL cartoon)]. Pathogenic variants within C3 largely cluster in a surface exposed region. (B) C6 domain and (C) C10 domain pathogenic variants do not cluster within a specific region of the domain. (D) Results of STRUM(18) analysis for *MYBPC3* pathogenic and benign variants within C3, C6, and C10 are shown, with mean and SEM for each group depicted. Graph is labeled to indicate variants predicted to be deleterious.

Conversely, of the pathogenic *MYBPC*3 variants not predicted to be deleterious by STRUM (Figure 3), a large number were localized within the C3 domain (Figure 3A, open circles; 7/13) and exhibited a mean ΔΔG -0.32 kcal/mol, (range -0.93 to 0.04). A large number of known pathogenic variants cluster within the C3 domain (15). This result is consistent with prior experimental and structural characterization data of these C3 pathogenic variants. Arg495Gln, Arg502Trp, and Phe503Leu incorporate normally into the sarcomere and have protein ½ lives that are indistinguishable from wild-type MyBP-C within primary cardiomyocytes^14^. Further, the NMR structure of the *MYBPC3* Arg502Trp C3 domain reveals preserved subdomain folding^32^. Pathogenic C3 domain variants cluster near a surface-exposed flexible linker region (Figure 4)^15^. Thus, these variants would be predicted to alter electrostatic protein-protein interactions but would not be expected to disrupt subdomain folding.

While C3 and C10 pathogenic variants have a narrow range of ΔΔG values, ΔΔG predictions for C6 pathogenic variants vary from -2.33 to 0.04 (mean ΔΔG -1.11). We previously examined two C6 domain variants, Arg810His and Trp792Arg, and found that they incorporate normally into the sarcomere and exhibit normal protein ½ lives in primary cardiomyocytes^14^. However, both of these variants were predicted to destabilize protein folding by STRUM, exhibiting values near the cutoff: Arg810His (ΔΔG - 1.22 kcal/mol), Trp792Arg (ΔΔG -1.28 kcal/mol). They are also predicted to be pathogenic by CardioBoost Table S2). These observations suggest that a subset of pathogenic variants mildly disrupt subdomain folding without causing complete destabilization of MyBP-C. Subdomain destabilization in these cases could interfere with protein-protein interactions or MyBP-C conformational dynamics.

### *In silico* saturation mutagenesis of *MYBPC3* identified 4,943 missense variants predicted to cause subdomain misfolding

Only a subset of amino acid substitutions has been observed in patients with HCM and are cataloged in publicly available databases, such as Clinvar. However, previously unreported variants frequently arise in probands with HCM who undergo clinical genetic testing^33^. Given the ability of STRUM to identify a subset of patients with a *MYBPC3* missense VUS which exhibit clinical outcomes similar to patients with a pathogenic *MYBPC3* variant, we also performed testing on all possible *MYBPC3* single amino acid substitutions (*in silico* mutagenesis) to develop a compendium of STRUM+ variants that may be useful for the research and clinical community. We found that 4,943 of 24,665 (20%) amino acid substitutions were predicted to disrupt subdomain folding (Figure S6, Table S5). The range of STRUM values were similar across most fibronectin and immunoglobulin domains with mean ΔΔG values ranging from -0.37 to -0.79 kcal/mol (Figure S6, Table S4). A notable exception was the C2 domain (C-score 0.42) which exhibited the lowest mean ΔΔG -1.11 +/- 0.99 kcal/mol value and the highest frequency of variants predicted to disrupt subdomain folding (P<0.0001). This analysis included synonymous variants and known pathogenic and benign variants. When these variants were excluded 4,929 of 23,298 (21%) amino acid substitutions were predicted to disrupt subdomain folding.

## Discussion

We have implemented a computational algorithm called STRUM, that incorporates sequence and structural data, and improves variant interpretation and prognostication for patients with HCM carrying a *MYBPC3* VUS. We have identified a subpopulation of patients with a *MYBPC3* missense VUS that are predicted to disrupt subdomain protein folding (STRUM+) who exhibit clinical outcomes indistinguishable from patients with a pathogenic *MYBPC3* variant. Conversely, patients carrying a *MYBPC3* VUS not predicted to affect subdomain folding (STRUM-), exhibit a lower prevalence of adverse clinical outcomes similar to patients with non-sarcomeric HCM. We have demonstrated that STRUM is capable of clinically risk stratifying HCM patients with a *MYBPC3* VUS and providing supporting evidence for improved variant adjudication. While computational prediction should not be exclusively relied on, these results suggest that results for STRUM could be incorporated into Bayesian models to improve prediction and prioritize variants that warrant further investigation in experimental studies or clinical co-segregation analyses.

Clinical risk stratification has been a cornerstone of clinical HCM management. It is well-established that patients with sarcomeric HCM have a higher rate of adverse clinical outcomes compared to non-sarcomeric HCM, enabling the incorporation of genetic data into clinical risk stratification in HCM^8^. Yet, refinement of clinical risk for patients with a VUS remains an ongoing challenge for clinicians^1, 5^. An important observation in this study is that STRUM enabled clinical risk stratification of patients carrying a missense VUS in *MYBPC3*. Although the methodology of parsing these variants is different for *MYBPC3* because of differing underlying mechanisms, these findings are analogous to a recent study in *MYH7* in which patients with HCM carrying VUSs that were located within the interacting-heads motif had a higher rate of adverse clinical outcomes compared to patients carrying VUSs that were outside of this motif^24^. These studies together suggest that VUSs in sarcomere genes are primarily an admixture of pathogenic and benign variants. So, while patients with HCM carrying sarcomere gene VUSs as a whole exhibit a prevalence of clinical outcomes that are intermediate between patients with or without pathogenic sarcomere variants^8^, a computational approach specifically leveraging the pathogenic mechanism of *MYBPC3* has enabled the identification of higher risk subpopulation that exhibit clinical outcomes similar to sarcomeric HCM and a lower risk subpopulation that exhibit clinical outcomes similar to non-sarcomeric HCM.

In addition to enabling clinical risk stratification, these results support the use of STRUM to provide supporting evidence for pathogenicity in an additive manner with other methods for variant adjudication. Given that previously unseen *MYBPC3* variants are frequently identified by genetic testing of probands with HCM^33^, we completed an *in silico* “saturation mutagenesis” of *MYBPC3* compiling a complete list of STRUM+ variants. STRUM+ *MYBPC3* variants identified in patients with HCM should be prioritized for additional clinical and experimental investigation. Specifically, functional experimental studies to evaluate the direct effects of *MYBPC*3 VUSs on protein stability, folding, and localization, as we have done previously for a subset of pathogenic variants^14^, will be important for improved variant adjudication. Further, familial co-segregation analysis on patients carrying a *MYBPC3* STRUM + VUS could further aid variant classification.

When known benign missense variants were evaluated by STRUM, 102 out of 110 variants were correctly predicted, with an overall specificity of 93%. However, for known pathogenic variants, only 7 of 19 were predicted to alter subdomain folding by STRUM, yielding a sensitivity of 32%. This, lower sensitivity was in large part explained by a known cluster of pathogenic variants within C3^15^. None of the 7 known pathogenic variants in C3 had a ΔΔG value below the threshold of -1.2 kcal/mol. This is consistent with experimental data that demonstrates C3 variants localize normally to the sarcomere and exhibit protein ½ lives similar to wild-type MyBP-C. Additionally, an NMR structure of Arg502Trp demonstrates that this variant does not disrupt subdomain folding but rather is more likely to alter protein-protein interactions^14, 32^. In contrast, MyBP-C pathogenic variants in C10, predicted by STRUM to cause subdomain misfolding, fail to localize to the sarcomere and are rapidly degraded^14^. Together these experimental results support the accuracy of STRUM predictions for subdomain misfolding. Further, they highlight the fact that STRUM is only predictive of pathogenicity for variants that significantly alter protein folding as their primary loss-of-function mechanism. Thus, a ΔΔG value of > -1.2 kcal/mol does not exclude pathogenicity for variants that may cause loss or gain-of-function through an alternate mechanism such as alternative splicing or altered protein-protein interactions. STRUM is best applied to VUSs after other methods for variant adjudication which identify pathogenic variants that may function via alterative mechanisms have been implemented. For example, *MYBPC3* pathogenic variants which lead to loss-of-function by mechanisms other than subdomain misfolding have previously been well characterized and defined as pathogenic, including splice variants^14, 22, 23^ and the cluster of pathogenic variants within C3 (aa.485-503)^15, 32, 34^ discussed above.

Of note, STRUM performed superiorly to sequence based-algorithms alone, such as SIFT and PolyPhen-2, which had lower specificity and were unable to clinically risk stratify patients with HCM and a *MYBPC3* missense VUS. We also examined, CardoBoost a recently developed disease-specific variant classifier for pathogenicity of rare missense variants in inherited cardiomyopathies^6^. CardioBoost demonstrated slightly higher specificity and sensitivity compared to STRUM for known benign and pathogenic variants. These algorithms identified distinct subsets of known pathogenic variants, suggesting they could be complimentary approaches. Consistent with this, combining the algorithms resulted in improved sensitivity to 63% while maintaining a specificity of 93%. Notably, CardioBoost identified three additional HCM patients with a missense VUS that would not otherwise be identified by STRUM and should be prioritized for further investigation. However, CardioBoost only identified 16/39 HCM patients with *MYBPC3* STRUM+ VUS as CardioBoost +. This result highlights the added utility of STRUM to identify a subset of VUSs within *MYBPC3* that exhibit allelic loss-of-function via local subdomain misfolding and are more likely to be pathogenic.

Another important observation in this study is that patients carrying a STRUM-variant exhibit clinical outcomes similar to patients lacking a sarcomere gene variant (Sarc -). The lower clinical risk profile of Sarc-patients and patients carrying a STRUM– *MYBPC3* VUS suggests a non-Mendelian form of HCM triggered by environmental factors, polygenic variation, and/or hypomorphic genetic variants of intermediate effect size^8, 35^.

Although this study was limited by a moderate sample size of 105 patients with HCM, the comprehensive variant adjudication in SHaRe enabled strict inclusion of patients carrying a single VUS within *MYBPC3* to clearly discriminate genetic-clinical correlates in this population. This approach enabled us to discern a difference in a composite of adverse clinical outcomes between patients with STRUM+ and STRUM-variants. However, the sample size was insufficient for detecting differences in individual outcomes, such as arrhythmias or heart failure. For this reason, the *in silico* saturation mutagenesis will be particularly valuable in enabling stratification and prioritization of novel variants that arise commonly in clinical testing for additional patients with HCM, so STRUM can be continually assessed as a tool for variant interpretation and clinical risk prediction over time.

The approach of using STRUM as an adjunctive tool for decision making may also be applicable to other genes for which loss-of-function is a pathogenic mechanism. Approximately 50% of disease associated variants within Human Gene Mutation Database are nonsense, splice, and frameshift variants predicted to result in loss-of-function^11^. These genes, like *MYBPC3*, also have missense VUSs that may be evaluated for protein misfolding using STRUM. For example, there are several causal genes for hypertrophic, dilated and arrhythmogenic cardiomyopathies with truncating pathogenic variants, including lamin A/C (*LMNA*), desmoplakin (*DSP*), and plakophilin 2 (*PKP2*), Titin (*TTN*), and phospholamban (*PLN*)^11, 36^. This approach is best suited for non-enzymatic proteins where high-quality structural modeling can be performed, and the primary pathogenic mechanism has been established to be loss-of-function.

## Conclusions

We show that a computational approach which predicts protein structure stability in response to missense variation enables identification of patients carrying a *MYBPC3* VUS who are at higher clinical risk of adverse events. This approach also provides supportive evidence for pathogenicity, prioritizing variants for functional experimental studies and clinical familial segregation to improve *MYBPC3* variant adjudication. Finally, the approach highlighted here may be broadly applicable to variants in other genes for which loss-of-function is an established mechanism.

## Data availability

The data will not be made available to other researchers for purposes of reproducing the results or replicating the procedure because of constraints related to human subjects research.

## Acknowledgements

This work was supported by the following sources; Funding for SHaRe has been provided by an unrestricted research grant by Myokardia, Inc a startup company that is developing therapeutics that target the sarcomere. Myokardia, Inc had no role in the preparation of this manuscript or approving the content of this manuscript. The following individuals are supported by the National Heart, Lung, and Blood Institute at the National Institutes of Health; Dr. Thompson [T32 HL007853], Dr. Helms [K08HL130455], Dr. Day [R01 11572784], Dr. Ho [1P50HL112349] & [1U01HL117006]. Dr. Thompson is supported by the protein folding disease initiative and Michigan Biology of Cardiovascular Aging (M-BoCA) at the University of Michigan. Dr Ware is supported by the Wellcome Trust [107469/Z/15/Z], the Medical Research Council (United Kingdom), the British Heart Foundation, National Institute for Health Research (NIHR), Royal Brompton Cardiovascular Biomedical Research Unit, and the NIHR Imperial College Biomedical Research Centre. Dr Ingles a recipient of a National Health and Medical Research Council. Dr Semsarian is the recipient of a National Health and Medical Research Council (NHMRC) Practitioner Fellowship [#1154992]. Dr Olivotto is supported by the Italian Ministry of Health [RF-2013-02356787] and [NET-2011-02347173] and by the Tuscany Registry of Sudden Cardiac Death (ToRSADE) project [FAS-Salute 2014, Regione Toscana].

## Author Information

Conceptualization – A.D.T., S.M.D.; Data curation-A.S.H., N.K.L., S.G.W., A.C.P., D.L.J., S.D.C., E.A.A., S.S., J.S.W., J.I., C.S., M.M., F.M., I.O., C.Y.H.; Formal Analysis, Investigation, Methodology-A.D.T., A.K., J.Y., A.S.H.,; Writing-original draft – A.D.T., S.M.D.; Writing-review & editing-A.S.H.; N.K.L., S.G.W., A.C.P., D.L.J., S.D.C., E.A.A., S.S., J.S.W., J.I., C.S., M.M., F.M., I.O., C.Y.H.

## Ethics Declaration

The generation of the centralized SHaRe database has been previously described (8). Data for this study were exported from quarter 1 of 2019. Exported data was de-identified. This study complies with the Declaration of Helsinki, Institutional review board and ethics approval was obtained in accordance with policies applicable to each SHaRe site and informed consent was obtained from all participants as required.

## Expanded Methods

### A. Variant Interpretation

SHaRe non-truncating *MYBPC3* missense variants (Supplemental Tables 1,2) were classified as previously reported(1) in accordance with American College of Medical Genetics and Genomics (ACMGG) guidelines, leveraging available clinical and experimental data (1-5). Herein we provide additional detail regarding this variant interpretation process. Clinically-indicated genetic testing was performed at all sites. Initial site designations of Pathogenic/Likely Pathogenic (pathogenic), variant of unknown significance (VUS), benign/likely benign (benign) were reviewed for accuracy in accordance with American College of Medical Genetics and Genomics (ACMGG) guidelines(6, 7) This review included review of available clinical and experimental data such as variant frequency (ShaRe and gnomAD) and evidence of segregation within families either previously published or within SHaRe. Further, splice site variants were considered pathogenic if they affected critically conserved splice consensus sites (i.e. acceptor -1 or -2; donor +1 or +2) and/or if experimental evidence showed aberrant splicing (1, 3, 5, 6). All potential exonic splice variants were identified and were cross-referenced to recent *in silico* and mini-gene assay analysis of MYBPC3 variants predicted to impact splicing (1, 3, 5, 6). None of the variants included in this study had experimental evidence of splicing, as variants with proven evidence of splicing were classified as truncating. Variants were considered “benign/likely benign” if the population allele frequency exceeded 0.004 and the odds ratio between SHaRe and gnomAD was <10-fold over the gnomAD allele frequency. If the variant was absent in gnomAD a conservative upper bound population allele frequency of 4.5E-06 was used for the odds ratio calculation (1). Since variants in sarcomere gene present in gnomAD with allele frequencies of > 4E-05 and absent in SHaRe are unlikely to be independently pathogenic for HCM, these variants were included in our list of benign MYBPC3 variants(1).

We also reference previously established subgroups within the SHaRe registry Sarc+, SarcU, Sarc-(8-10). Sarc+ is defined as sarcomeric HCM or patients with HCM carrying a pathogenic or likely pathogenic variant within 8 sarcomere genes definitively associated with HCM (MYBPC3, MYH7, TNNT2, TNNI3, TPM1, MLY2, MYL3, ACTC) (8). SarcU (also referred to as SarcVUS in previous publications) is defined as HCM patients carrying a sarcomere variant of uncertain significant(s) and Sarc- is defined as non-sarcomeric HCM or patients with HCM who underwent genetic testing and had no pathogenic, likely pathogenic, or VUS variant identified within a sarcomere gene. Patients were excluded from Sarc- if they had pathogenic variants in genes encoding non-sarcomere proteins such as GLA or LAMP2.

### B. Computational Protein Folding Stability Predictive Modeling

For *MYBPC3*, which encodes protein MyBP-C, missense variants were analyzed using STRUM. STRUM calculates the effect of the missense variant on the Gibbs free energy of local subdomain folding (referred to as the ΔΔG, kcal/mol)(11). Subdomains were selected instead of the full-length protein to enable more accurate structural modeling (Supplemental Table 3). STRUM analysis of *MYBPC3* variants was performed using amino acid sequence inputs (Supplemental Table 3) and method 2, completed 05/2020. This enabled a full analysis of all possible MYBPC3 subdomain variants (Supplemental Table 6).

A negative ΔΔG value indicates the degree of reduced folding energy (kcal/mol) relative to the wild-type subdomain, or folding destabilization (11). Previous experimental validation of this algorithmƒ compared STRUM predictions to 3421 experimentally tested variants from 150 proteins and demonstrated a strong correlation with a Pearson’s correlation coefficient (PCC) of 0.79 and root mean square error (RMSE) of prediction of 1.2 kcal/mol (11). A value of ΔΔG ≤-1.2 kcal/mol was defined as the cut-off for destabilizing (deleterious) variants while a value of ΔΔG > -1.2 was defined as the cut-off for variants that did not significant destabilize subdomain folding (non-deleterious) variants.

### C. Computational Structural Modeling Information

The STRUM algorithm utilizes I-TASSER structural models in calculating ΔΔG. I-TASSER (Iterative Threading ASSEmbly Refinement) is a hierarchical approach to protein structure and function prediction(12-14). MyBP-C is made up of immunoglobulin and fibronectin subdomains(15). Structural predictions of MyBP-C (the protein encoded by *MYBPC3*) subdomains C0-C10 were obtained using I-TASSER and are depicted using PyMOL (Supplemental Figure 1). Two linker regions were modeled. The first is a proline rich region between C0 and C1. The second is the M-domain linker between C1 and C2. For variants within these linker regions, a structural model of two adjacent subdomains and the linker region of interest were established (Supplemental Figure 2). Linker regions were otherwise assumed to consistent of short flexible unstructured linkers and were not analyzed using STRUM (Supplemental Figure 3, Supplemental Table 4).

High quality I-TASSER models were produced for each sub-domain (Supplemental Figure 1-3, Supplemental Table 3)(12, 13). To describe quality metrics in more detail; the C-score is a confidence score for estimating the quality of predicted models by I-TASSER It is calculated based on the significance of threading template alignments and the convergence of parameters of the structure assembly simulations. C-score is typically in the range of [-5,2], where a C-score of higher value signifies a model with high confidence. Subdomains C0-C10 exhibited C-scores ranging from 0.22-0.98. Another measure of model quality is TM-score, which is sensitive to local error. A TM-score > 0.5 indicates a model with correct topology. Our models exhibited TM-scores ranging from 0.74-0.84. Finally, we report a root mean square deviation (RMSD) compared to known MyBP-C subdomain structures, which ranged from 0.29-0.44 Angstroms(16-23), exhibiting good agreement between experimental and modeling data.

With the exception of two linker regions, individual subdomains are connected by short flexible linkers which were not modeled (Supplemental Figure 3). The two larger linker regions present in MyBP-C are; the proline rich region between C0 and C1 domains and the M-domain between C1 and C2 domains(15). These linker regions were modeled in the presence of their two adjacent subdomains; C0-Proline rich linker-C2 (aa 1-260) and C1-M domain-C2 (aa 152-452) using I-TASSER (Supplemental Figure 2). This revealed a flexible proline rich linker with minimal secondary structure and a structured M- domain made up of beta-sheets. The M-domain model is distinct from a truncated region of M-domain that was evaluated connected to C2 domain by NMR (319-451) which suggested a triple helix within this C-terminal portion of the M-domain (PDB 5k6p.pdb) (21). These models were of acceptable quality with C-score of -2.01 and -0.72 and TM score 0.47 and 0.62 respectively, albeit lower quality than individual subdomains. As such these models were only utilized to evaluate variants within the linker region. All depictions of I-TASSER models and alignments with known MyBP-C structures were performed using PyMOL (Supplemental Figure 1-3).

Finally, a model of full length *MYBPC3* based on axial-radial model (24) using these individual subdomains was assembled (Supplemental Figure 3). This model serves to illustrate the well-defined subdomain immunoglobulin and fibronectin folds within *MYBPC3* and assumes independent motion of individual subdomains connected by flexible linker regions. The exceptions to this assumption are the proline-rich (PR) linker between CO and C1 and the M-domain linker between C1 and C2 which were modeled as described above. There are multiple proposed models of MyBP-C quaternary structure within the sarcomere based on electron microscopy data (24-27). Further experimental work is needed to more accurately model orientation between subdomains, potential interdomain interactions, and *MYBPC3* quaternary structure.

### D. Clinical Outcomes Analysis

Only HCM patients with a single *MYBPC3* variant were included in clinical outcomes analysis to avoid confounding from multiple cases with multiple sarcomere gene variants (Figure 3). To expand on this inclusion criterion further, we began by identifying 426 SarcU patients within SHaRe. As previously described(8) SarcU patients are HCM patients whom have undergone clinical genetic testing and lack pathogenic or likely pathogenic variants within sarcomere genes or other potentially pathogenic genes such as GLA or LAMP2 but do carry a VUS within a sarcomere gene. We next limited our analysis to patients with a single VUS [VUS variants (all genes): one], excluding patients that carry multiple sarcomere VUSs. Finally, we limited our analysis to patients carrying a single missense MYBPC3 VUS [VUS variants (MYBPC3): STRUM] (Figure 3). For comparison groups (Figure 4) we evaluated HCM patients with a single MYBPC3 pathogenic variant (MYBPC3-Path-all). To identify this group we utilized the previously defined Sarc+ group (8), limiting our evaluation to patients with single pathogenic sarcomere variant (P/LP variants: One) and excluding patients carrying another VUS (VUS variants: zero). Our final comparison group was the previously defined Sarc-group (8).

### E. Defining Clinical outcomes

Composite clinical outcomes evaluated herein as previously defined(8) are;

- Ventricular arrhythmic composite: first occurrence of sudden cardiac death, resuscitated cardiac arrest, or appropriate implantable cardioverter-defibrillator therapy
- Heart Failure composite: first occurrence of cardiac transplantation, LV assist device implantation, LV ejection fraction <35%, or New York Heart Association class III/IV symptoms
- Overall composite: first occurrence of any component of the ventricular arrhythmic or heart failure composite end point (without inclusion of LV ejection fraction), all-cause mortality, atrial fibrillation (AF), stroke, or death

### F. Statistical Analysis

Odds ratio(OR), sensitivity, and specificity were calculated to evaluate the association between computational algorithms (STRUM, SIFT, PolyPhen-2) STRUM predictions of deleterious variants and known *MYBPC3* pathogenic and benign variants as follows;

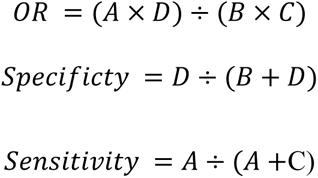

where

*A = variants predicted the be deleterious and known to be pathogenic,*

*B = variants predicted to be deleterious and known to be benign,*

*C= variants predicted to be non-deleterious and known to be pathogenic,*

*D= variants predicted to be non-deleterious and known to be benign*.

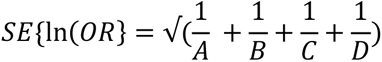

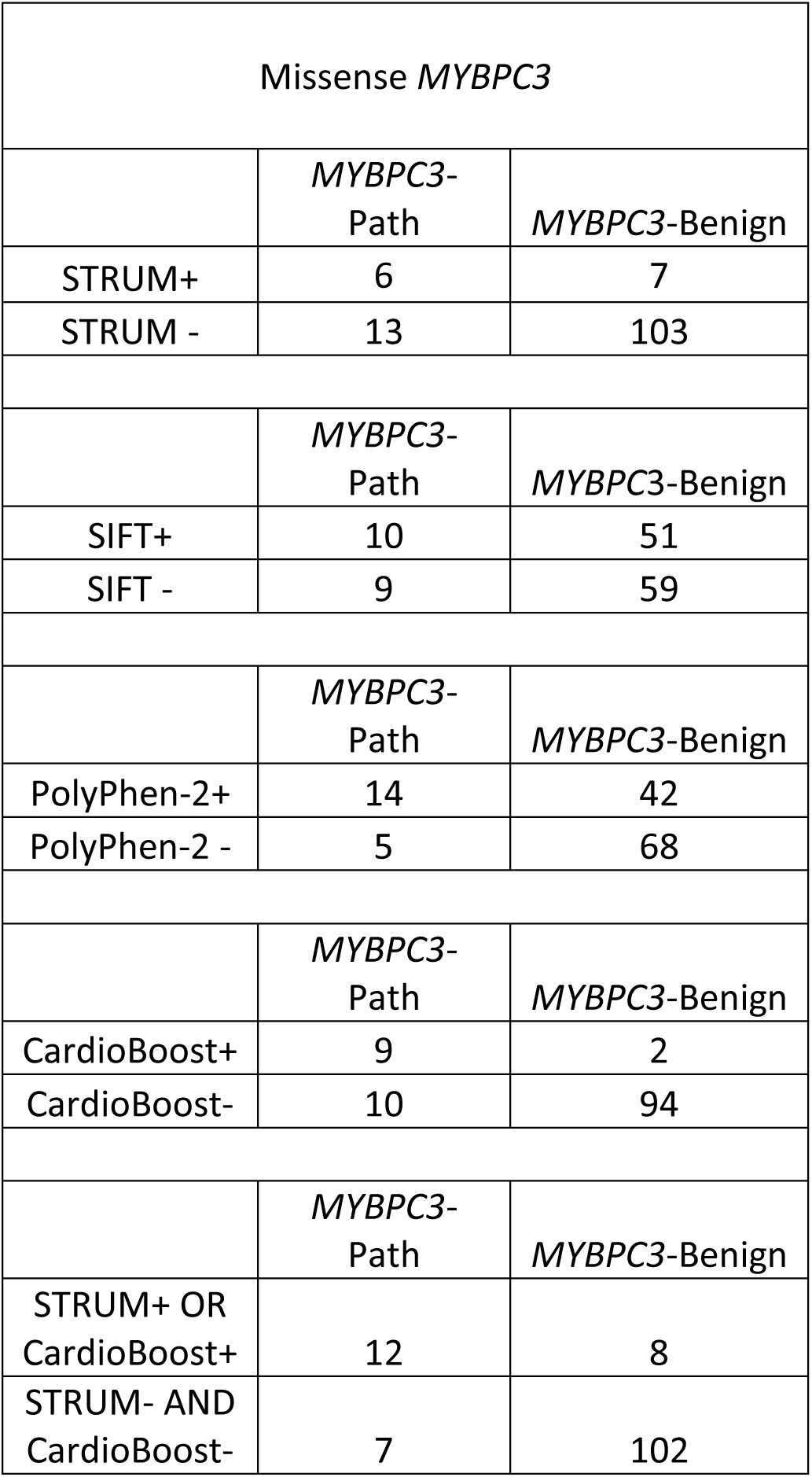

| Missense MYBPC3 |  |  |
| --- | --- | --- |
|  | MYBPC3-Path | MYBPC3-Benign |
| STRUM+ | 6 | 7 |
| STRUM - | 13 | 103 |
|  | MYBPC3-Path | MYBPC3-Benign |
| SIFT+ | 10 | 51 |
| SIFT - | 9 | 59 |
|  | MYBPC3-Path | MYBPC3-Benign |
| PolyPhen-2+ | 14 | 42 |
| PolyPhen-2 - | 5 | 68 |
|  | MYBPC3-Path | MYBPC3-Benign |
| CardioBoost+ | 9 | 2 |
| CardioBoost- | 10 | 94 |
|  | MYBPC3-Path | MYBPC3-Benign |
| STRUM+ OR CardioBoost+ | 12 | 8 |
| STRUM- AND CardioBoost- | 7 | 102 |

### G. Figures

**Figure S1:**
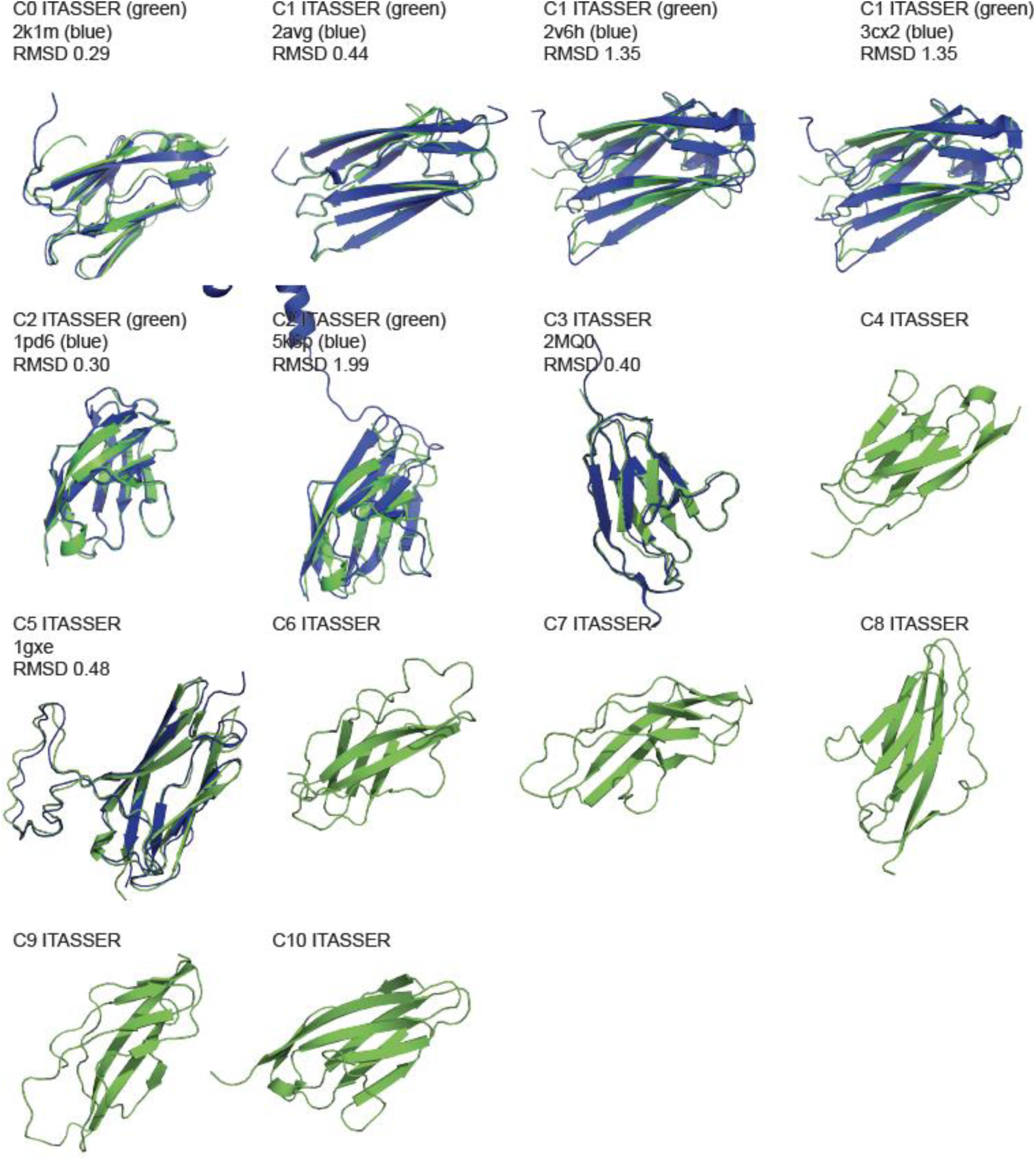
MyBP-C subdomain I-TASSER models. I-TASSER models were derived for MyBP-C subdomains C0-C10 from sequence inputs as detailed in Supplemental Table 3 are shown in green (PyMOL, cartoon). These structures were aligned with all available known MyBP-C subdomain structures shown in blue (PyMOL, cartoon) (16-23). For each alignment the pdb code of the experimentally derived structure and the root mean square deviation (RMSD) of the alignment is reported. Subdomains C4, C6, C7, C8, C9, C10 did not have available structural data for comparison.

**Figure S2:**
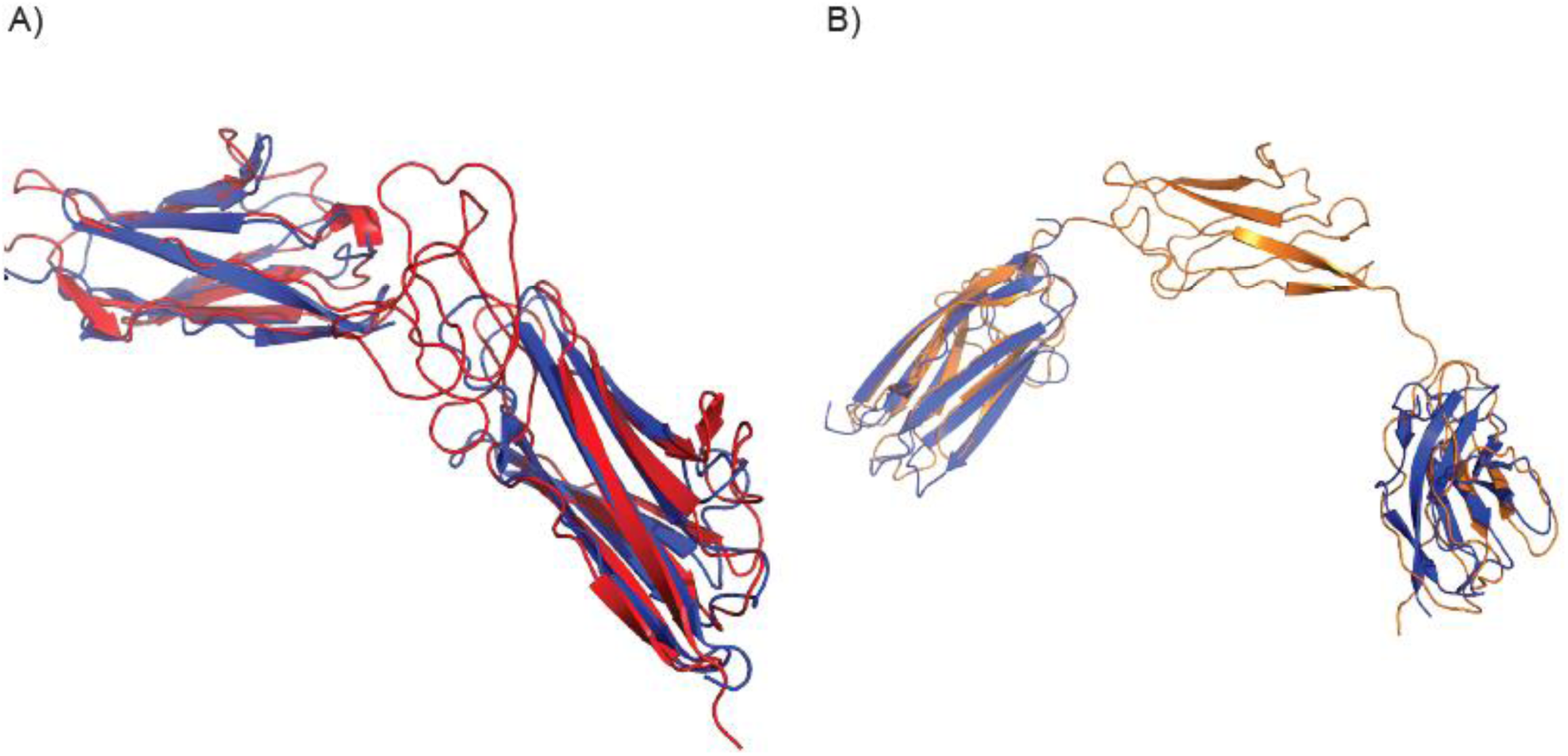
I-TASSER Models of proline rich and M-domain linkers. I-TASSER models of these two linker regions were made by modeling the linker regions with two adjacent domains. A) The proline rich (PR) linker model [C0-PR-C1 (aa 1-260)] is shown in red (PyMOL, cartoon). This model was aligned with known MyBP-C C0 and C1 subdomain structures shown in blue (PyMOL, cartoon) (2k1m.pdb and 2avg.pdb respectively). B) The M-domain linker model [C1-M domain-C2 (aa 152-452)] is shown in orange (PyMOL, cartoon). This model was aligned with known MyBP-C C1 and C2 subdomain structures shown in blue (PyMOL, cartoon). (2avg.pdb, and 1pd6.pdb respectively) (16, 17)

**Figure S3:**
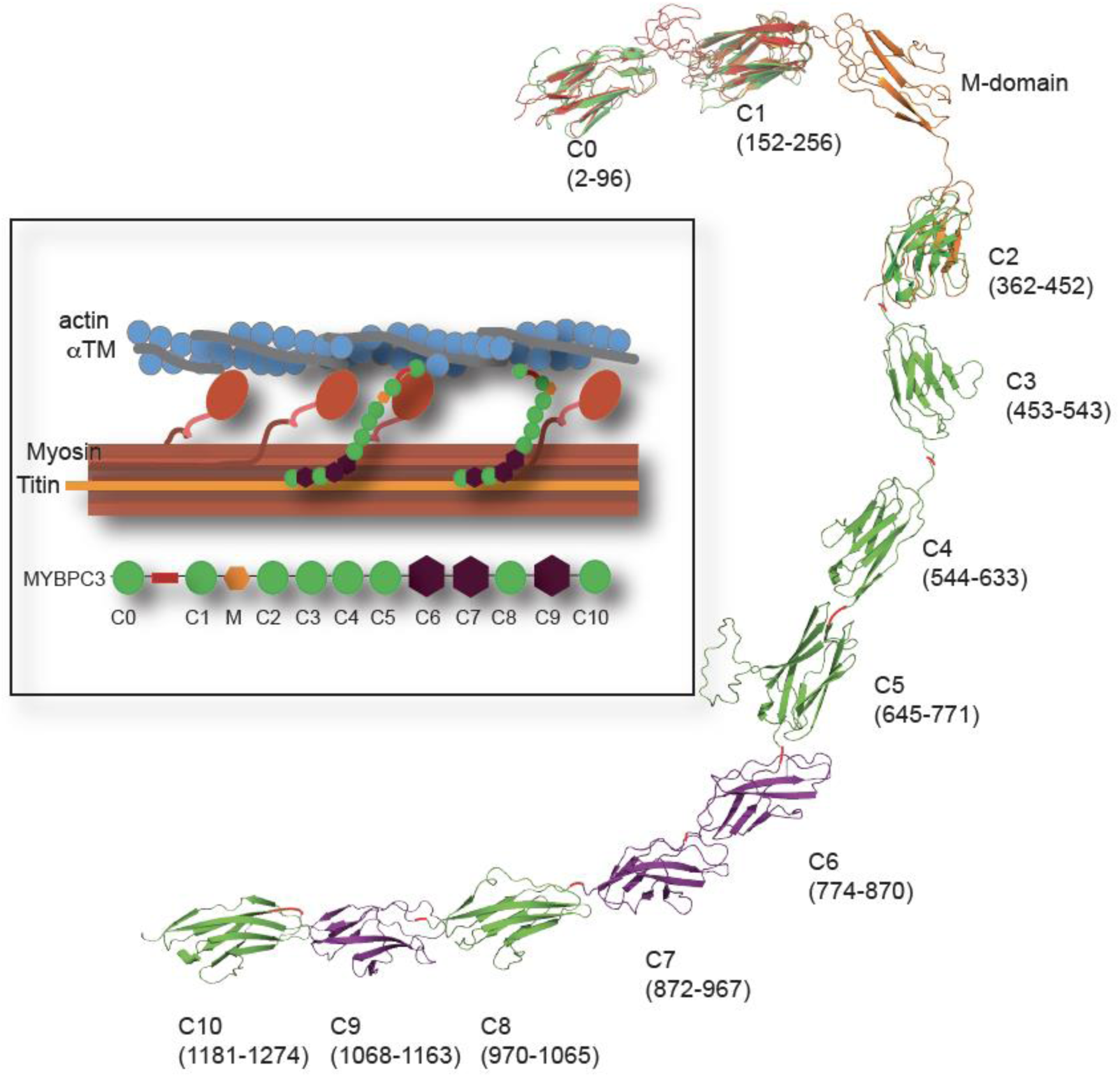
Structural model of MyBP-C. MyBP-C (the protein encoded by *MYBPC3*) is made up of immunoglobin (green circles) and fibronectin domains (purple hexagons) connected by short flexible linkers. Two larger linker regions exist; the proline-rich (PR) linker (red rectangle) between C0 and C1 domains and the M-domain linker (small orange hexagon) between C1 and C2 domains. MyBP-C is positioned in an anti-parallel fashion within the A-band of sarcomere. The N-terminus (C0-C2) interacts with actin and myosin in a dynamic fashion while the C-terminus (C7-C10) interacts with thick filament and Titin. A structural model was built using I-TASSER subdomain models. Individual immunoglobulin (green PyMOL cartoon) and fibronectin (purple PyMOL cartoon) I-TASSER models are arranged based on previously proposed quaternary structural model(24). In addition, the C0-PR-C1 (red, PyMOL cartoon) and C1-Mdomain-C2 (orange PyMOL cartoon) I-TASSER models were aligned with the individual I-TASSER models for these subdomains. This model otherwise assumes independent motion of individual subdomains connected by flexible linker regions (red lines).

**Figure S4:**
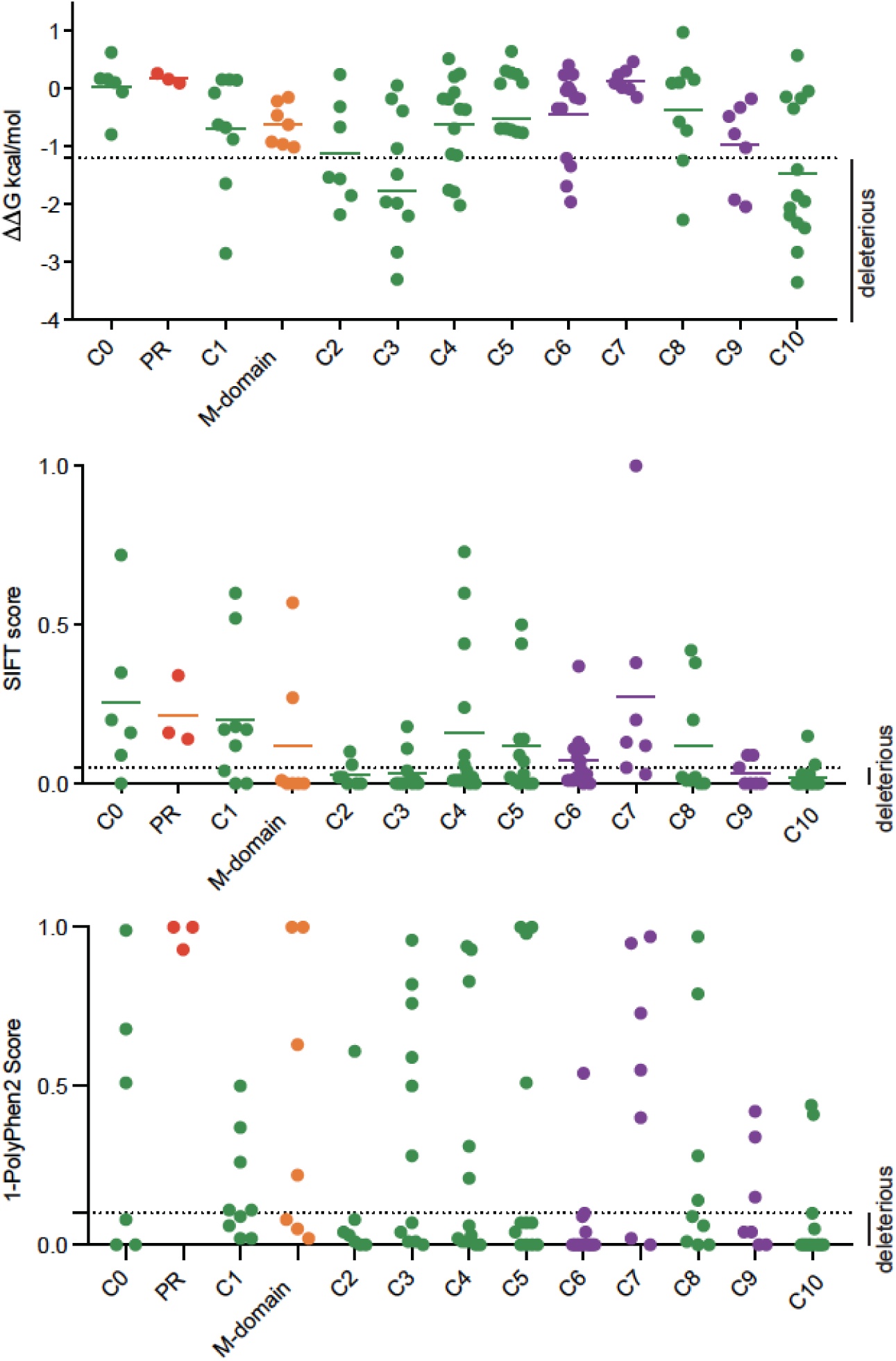
Computational analysis of missense *MYBPC3* VUSs. Missense *MYBPC3* VUS within SHaRe (Supplemental Table 2) were analyzed by STRUM (top panel), SIFT (middle panel) and PolyPhen-2 (lower panel). Results are grouped by individual subdomains with immunoglobulin domains (green; C0, C1, C2, C3, C4, C5, and C8) fibronectin domains (purple; C6, C76, C9). The linker regions - proline rich (PR, red) and M-domain (orange) linkers. The cut-off for likely deleterious variants (SIFT < or = 0.05, 1- PolyPhen-2 < 0.10) is designated by dotted line with region below representing variants predicted to be deleterious. In the STRUM analysis, the proportion of *MYBPC3* VUSs predicted to be deleterious varied by subdomain. For example, a higher proportion of VUSs were predicted to be deleterious within the C3 (63%) C2 (57%) and C10 domains (38%) compared to other immunoglobulin domains. High quality ITASSER models were obtained for each subdomain (Supplemental Figure 1). Of note very few variants within SHaRe were present within linker regions outside of defined subdomains (4%) (Supplemental Table 1,2). Two linker domains within MYBP-C accounted for the majority of linker region variants (22/24) and were modeled (Supplemental Figure 2-3). The PR domain was predicted to be a flexible and largely unstructured linker whereas the M-domain was predicted to exhibit beta-sheet secondary structure (Supplemental Table 3, Supplemental Figure 2). *MYBPC3* VUSs within the PR linker were consistently predicted to be non-deleterious not only by STRUM but also by the sequence-based algorithms Polyphen-2 and SIFT (Supplemental Figure 4).

**Figure S5:**
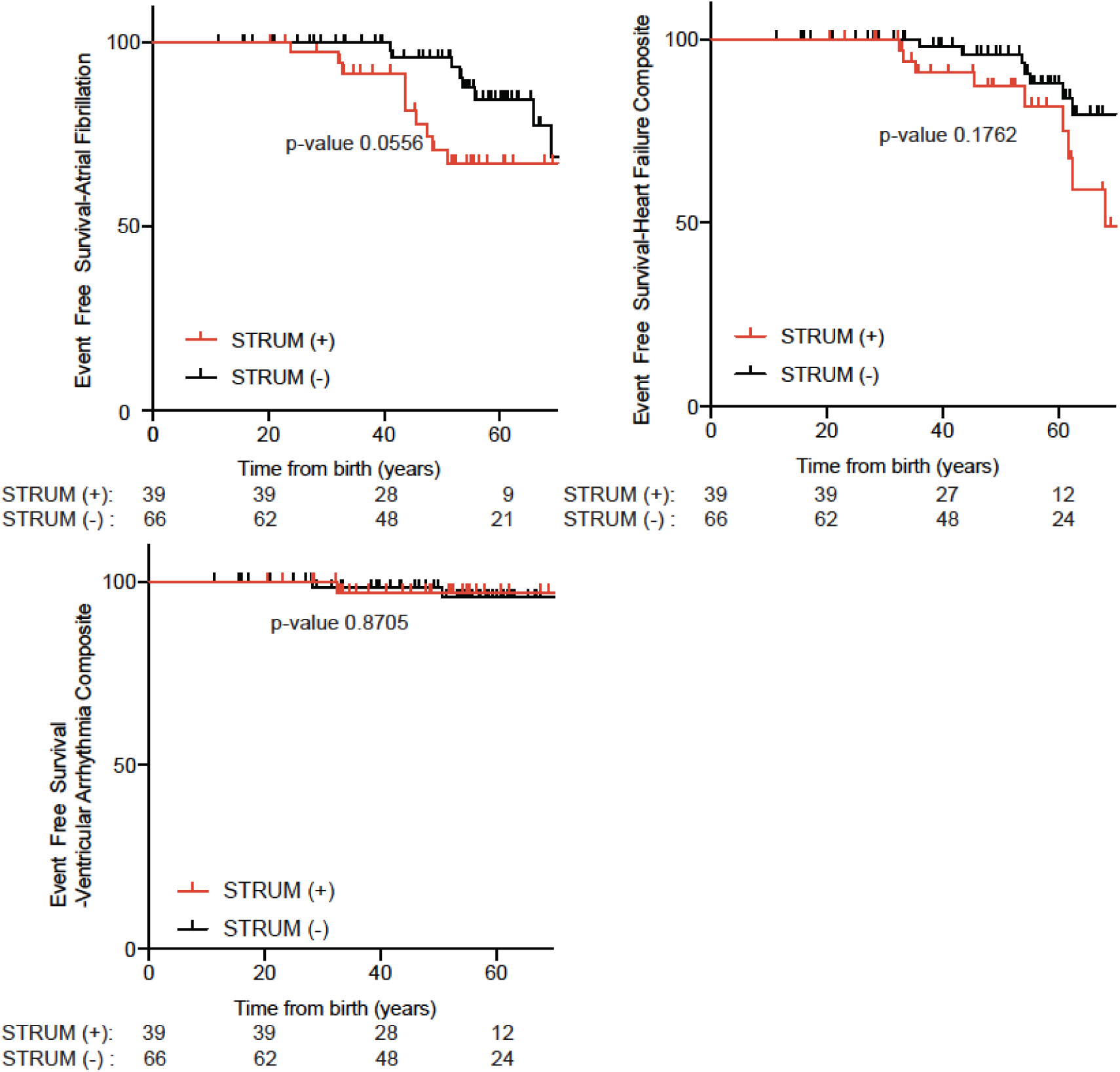
Risk stratification of individual composite outcomes. Variants predicted to be deleterious by STRUM (STRUM+, ΔΔG ≤ -1.2 kcal/mol, red, n =39) did exhibit higher rates of atrial fibrillation and heart failure composite than variants not predicted to be deleterious STRUM-, black, n=66). However, we could not exclude the null hypothesis (p -value 0.0559 and 0.1762). No difference in ventricular arrhythmia composite was observed. As previous described(8) composite outcomes reported are defined as follows;

**Figure S6:**
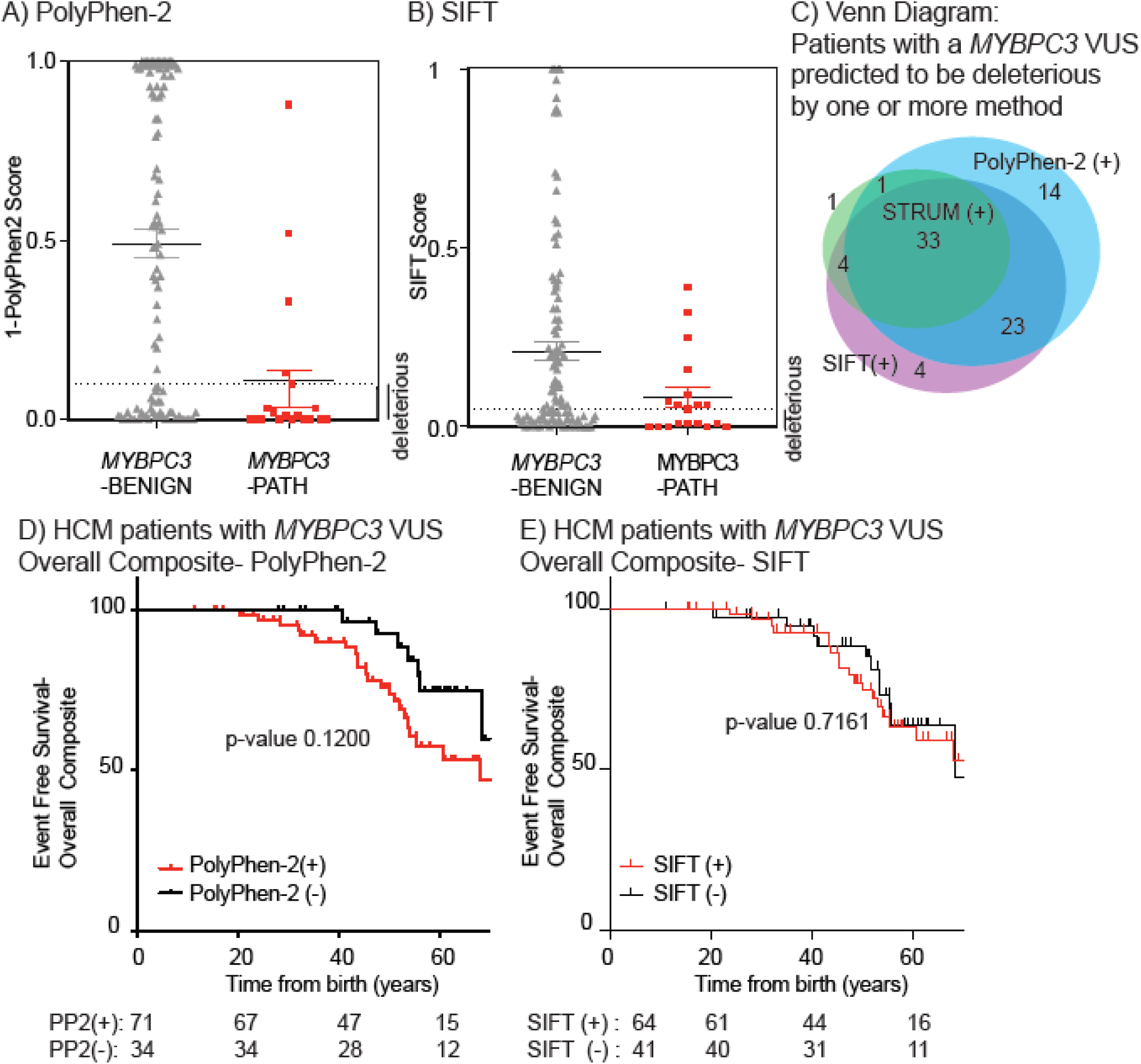
Sequence-based algorithms were exhibit decreased specificity compared to STRUM and are unable to identify a subgroup of HCM patients with a *MYBPC3* VUS at increased risk for HCM-related adverse clinical outcomes. Results of computational analysis for each unique *MYBPC3*-Benign (grey triangles, n= 110) and *MYBPC3*-Path (red circles, n =19) variant are shown. Mean and SEM for each group depicted. The cut-off for deleterious variants were (A) Polyphen-2 > 0.90 (graphed as 1-Polyphen-2 score < 0.10) and (B) SIFT ≤ 0.10. Both SIFT and Polyphen-2 demonstrated decreased specificity 62% and 54% respectively compared to STRUM 93% (Figure 3). (C) We next evaluated patients with HCM and a MYBPC3 VUS. A ven diagram is shown for patients with HCM and a MYBPC3 VUS predicted to be deleterious by one or more of the following algorithms; SIFT, PolyPhen-2, STRUM. (D) Variants predicted to be deleterious by PolyPhen-2 (PolyPhen-2 > 0.90, red, n =71)(28) did exhibit higher rates of adverse HCM-related clinical outcomes (Overall composite) compared to variants not predicted to be deleterious (black, n = 34) by Kaplan Meier event free survival analysis. However, we could not exclude the null hypothesis (p-value 0.1200). As defined previously, the overall composite (8) is defined as the first occurrence of any component of the ventricular arrhythmic or heart failure composite end point (without inclusion of LV ejection fraction), all-cause mortality, atrial fibrillation (AF), stroke, or death. (E) Variants predicted to be deleterious by SIFT(29) (SIFT score ≤0.05, red, n =64) and non-deleterious (black, n =41) exhibited similar rates of adverse HCM-related clinical outcomes.

**Figure S7:**
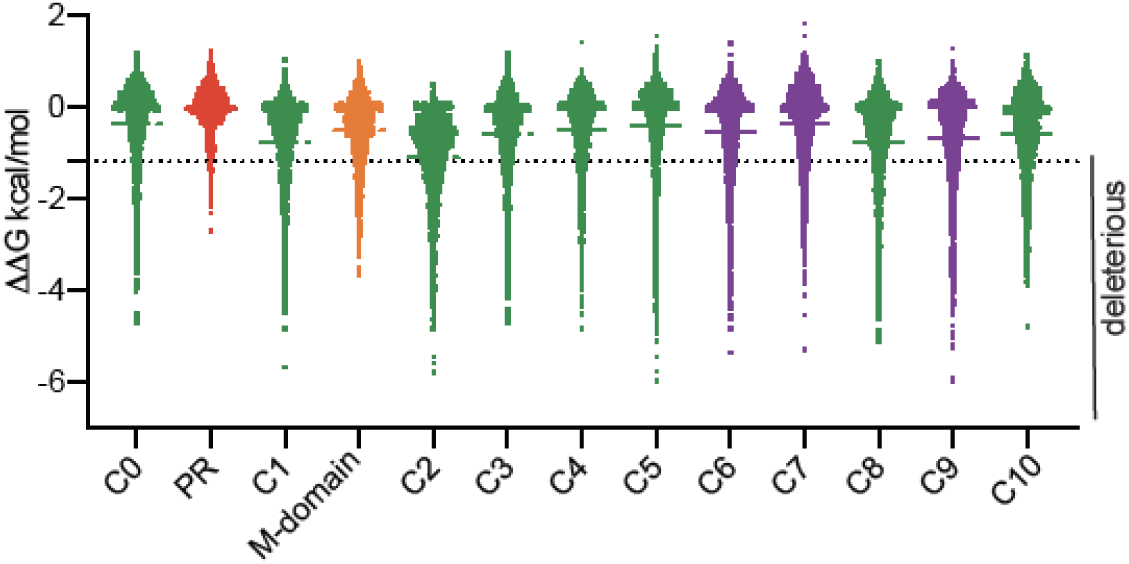
STRUM Analysis of *MYBPC3* VUSs. All possible MyBP-C missense and synonymous variants were analyzed by STRUM by performing *in silco* saturation mutagenesis (Supplemental Table 5). Subdomains of MyBP-C (the protein encoded by *MYBPC3*) are labeled and colored by domain type - immunoglobulin (C0, C1, C2, C3, C4, C5, and C8) in green, fibronectin (C6, C6, C9) in purple, and the proline rich (PR) and M-domain linkers in red and orange respectively. Horizontal line indicates mean for each group. Variants with ΔΔG ≤-1.2 kcal/mol are predicted to be deleterious. The cluster of variants at zero represent largely synonymous variants.

**Supplemental Table 1:** Computation analysis of *MYBPC3* non-synonymous missense variants-Benign and Path

| ***MYBPC3*-Likely Benign/Benign (Benign) Variants (SHaRe, and GnomAD)** |  |  |  |  |  |  |
| --- | --- | --- | --- | --- | --- | --- |
| **Amino Acid Variant** | **MyBP-C Domain** | **SHaRe Number of Cases** | **STRUM(11)**<br>**(deleterious $\leq -1.2$)** | **SIFT(29)**<br>**(deleterious $\leq 0.05$)** | **Polyphen-2(28)**<br>**(deleterious $> 0.90$)** | **CardioBoost(10)**<br>**(deleterious $> 0.90$)**<br>**(Benign $< 0.10$)** |
| p.Gly5Arg | 0 | 2 | 0.57 | 0.03 | 0.10 | 0.006 |
| p.Arg17Gln | 0 | 1 | 0.25 | 0.06 | 0.21 | 0.030 |
| p.Ser25Asn | 0 | 0 | 0.17 | 0.04 | 0.72 | 0.040 |
| p.Val28Met | 0 | 0 | -0.27 | 0.02 | 0.58 | 0.105 |
| p.Glu32Lys | 0 | 1 | -0.77 | 0.33 | 1.00 | 0.040 |
| p.Arg44His | 0 | 0 | -0.36 | 0.28 | 0.95 | 0.028 |
| p.Thr62Pro | 0 | 1 | -0.53 | 0.16 | 0.16 | 0.003 |
| p.Arg63Trp | 0 | 0 | 0.05 | 0.02 | 0.79 | 0.012 |
| p.Pro108Leu | Proline rich | 0 | 0.05 | 0.20 | 0.02 | 0.002 |
| p.Pro121Leu | Proline rich | 1 | 0.50 | 0.02 | 0.00 | 0.002 |
| p.Ser138Gly | Proline rich | 0 | -0.28 | 0.03 | 0.02 | 0.002 |
| p.Pro147Leu | Proline rich | 0 | 0.24 | 0.02 | 0.09 | 0.003 |
| p.Ile154Thr | 1 | 1 | -0.83 | 0.88 | 0.02 | 0.017 |
| p.Val158Met | 1 | 3 | -0.50 | 0.03 | 0.45 | (-) |
| p.Arg160Trp | 1 | 1 | 0.23 | 0.00 | 0.98 | (-) |
| p.Glu165Asp | 1 | 1 | -0.07 | 0.58 | 0.45 | 0.158 |
| p.Val168Ala | 1 | 0 | -0.44 | 0.41 | 0.00 | 0.0004 |
| p.Val168Met | 1 | 0 | -0.06 | 0.07 | 0.46 | 0.018 |
| p.Arg177His | 1 | 3 | -0.65 | 0.01 | 0.63 | (-) |
| p.Pro186Leu | 1 | 2 | -0.01 | 0.21 | 0.68 | 0.036 |
| p.Val189Ile | 1 | 4 | -0.62 | 0.43 | 0.37 | (-) |
| p.Gln208His | 1 | 1 | 0.19 | 0.12 | 0.99 | 0.375 |
| p.Arg215Cys | 1 | 0 | 0.16 | 0.02 | 0.98 | 0.005 |
| p.Ala216Thr | 1 | 3 | -0.30 | 0.92 | 0.00 | 0.001 |
| p.Ser217Gly | 1 | 9 | -0.05 | 0.10 | 0.61 | (-) |
| p.Glu223Lys | 1 | 0 | -0.47 | 0.00 | 1.00 | 0.303 |
| p.Asp228Asn | 1 | 0 | 0.01 | 0.10 | 0.07 | 0.003 |
| p.Asp228Glu | 1 | 0 | 0.02 | 0.88 | 0.00 | 0.005 |
| p.Ser236Gly | 1 | 5 | -1.18 | 0.71 | 0.00 | (-) |
| p.Arg238His | 1 | 0 | -0.73 | 0.01 | 1.00 | 0.960 |
| p.Asn253Ser | 1 | 0 | -0.38 | 0.23 | 0.02 | 0.007 |
| p.Gly263Arg | M | 1 | -0.74 | 0.04 | 0.10 | 0.002 |
| p.Arg272Cys | M | 0 | -0.12 | 0.00 | 0.77 | 0.193 |
| p.Gly278Glu | M | 1 | -0.60 | 0.00 | 0.16 | (-) |
| p.Arg281Gln | M | 0 | -0.41 | 0.04 | 0.01 | 0.008 |
| **p.Val321Met** | **M** | **4** | **-1.36** | **0.01** | **0.52** | **0.676** |
| p.Arg326Gln | M | 4 | -0.55 | 0.30 | 0.01 | (-) |
| **p.Glu334Lys** | **M** | **2** | **-1.38** | **0.06** | **0.33** | **0.112** |
| p.Ala364Val | 2 | 0 | -0.94 | 0.41 | 0.97 | 0.036 |
| p.Arg382Trp | 2 | 0 | -1.17 | 0.02 | 0.99 | (-) |
| **p.Leu383Val** | **2** | **1** | **-2.55** | **0.02** | **0.98** | **0.2466** |
| p.Ile415Leu | 2 | 0 | -0.6 | 0.38 | 0.00 | 0.006 |
| p.Gly416Ser | 2 | 1 | -1.04 | 0.00 | 0.98 | 0.069 |
| p.Ala429Val | 2 | 1 | -0.30 | 0.36 | 0.52 | 0.003 |
| p.Glu441Lys | 2 | 7 | -0.57 | 0.02 | 0.91 | 0.042 |
| p.Thr457Met | 3 | 0 | 0.02 | 0.06 | 0.92 | 0.010 |
| p.Arg458His | 3 | 0 | -0.04 | 0.30 | 0.04 | 0.044 |
| p.Gly490Arg | 3 | 7 | -1.06 | 0.00 | 1.00 | 0.411 |
| p.Gly507Arg | 3 | 6 | -0.86 | 0.00 | 1.00 | 0.169 |
| p.Asn515Ser | 3 | 1 | 0.09 | 0.97 | 0.07 | 0.021 |
| p.Ala522Thr | 3 | 1 | -0.51 | 0.18 | 0.01 | 0.006 |
| p.Gly596Arg | 4 | 2 | -0.84 | 0.00 | 0.99 | 0.247 |
| p.Asp605Asn | 4 | 3 | 0.34 | 0.27 | 0.07 | 0.064 |
| p.Glu619Lys | 4 | 12 | 0.04 | 0.08 | 0.43 | 0.029 |
| p.Val639Ile | Linker C4-C5 | 0 | NA | NA | NA | NA |
| p.Ile655Thr | 5 | 0 | -0.81 | 1.00 | 0.00 | 0.004 |
| p.Leu667Val | 5 | 0 | -0.99 | 0.70 | 0.60 | 0.381 |
| p.Arg668His | 5 | 2 | -0.49 | 0.05 | 0.98 | 0.124 |
| p.Asp709Asn | 5 | 0 | 0.77 | 0.40 | 0.01 | 0.004 |
| p.Arg726Cys | 5 | 0 | -0.13 | 0.01 | 0.99 | 0.003 |
| p.Arg733Cys | 5 | 1 | -0.09 | 0.21 | 0.03 | 0.009 |
| p.Ser734Gly | 5 | 0 | -0.66 | 0.05 | 0.80 | 0.411 |
| p.Val748Ile | 5 | 0 | -0.2 | 0.11 | 0.01 | 0.009 |
| p.Val757Met | 5 | 0 | 0.13 | 0.04 | 0.86 | 0.235 |
| p.Pro794Leu | 6 | 1 | 0.38 | 0.01 | 0.99 | 0.024 |
| p.Ala833Thr | 6 | 3 | -1.41 | 0.42 | 0.91 | (-) |
| p.Ala833Val | 6 | 0 | -0.89 | 1.00 | 0.97 | (-) |
| p.Arg835Leu | 6 | 0 | 0.07 | 0.03 | 1.00 | 0.666 |
| p.Ala851Val | 6 | 1 | 0.52 | 0.12 | 0.92 | 0.234 |
| p.Met854Val | 6 | 0 | 0.46 | 1.00 | 0.02 | 0.011 |
| p.Gly868Ser | 6 | 0 | 0.20 | 0.20 | 0.47 | 0.014 |
| p.Glu872Lys | 7 | 2 | -0.10 | 0.17 | 0.98 | 0.081 |
| p.Pro873His | 7 | 3 | -0.80 | 0.00 | 1.00 | 0.335 |
| p.Pro873Leu | 7 | 0 | -1.07 | 0.00 | 1.00 | 0.374 |
| p.Thr885Met | 7 | 0 | -0.02 | 0.02 | 0.99 | 0.167 |
| p.Arg895His | 7 | 0 | 0.45 | 0.22 | 0.32 | 0.609 |
| p.Val896Met | 7 | 8 | -0.09 | 0.04 | 0.09 | (-) |
| p.Pro910Thr | 7 | 2 | 0.36 | 0.26 | 0.00 | 0.002 |
| p.Gln921Glu | 7 | 1 | -0.02 | 0.54 | 0.01 | 0.006 |
| p.Gln921Leu | 7 | 0 | 0.4 | 0.21 | 0.00 | 0.004 |
| p.Thr924Ile | 7 | 0 | 0.24 | 0.57 | 0.01 | 0.0002 |
| p.Leu934Pro | 7 | 0 | -2.72 | 0.01 | 1.00 | 0.967 |
| p.Thr957Ser | 7 | 4 | -0.01 | 0.39 | 0.00 | 0.001 |
| p.Thr958Ile | 7 | 3 | 0.58 | 0.18 | 0.01 | 0.003 |
| p.Pro961Leu | 7 | 1 | 0.11 | 0.02 | 0.39 | 0.002 |
| p.Arg972Trp | 8 | 1 | -0.24 | 0.01 | 0.01 | 0.021 |
| p.Arg980Cys | 8 | 0 | -0.13 | 0.00 | 1.00 | 0.085 |
| p.Gln998Glu | 8 | 5 | 0.32 | 0.02 | 0.95 | (-) |
| p.Arg1002Trp | 8 | 3 | 0.05 | 0.00 | 1.00 | 0.387 |
| p.Glu1017Lys | 8 | 0 | -0.4 | 1.00 | 0.00 | 0.012 |
| p.Arg1036Cys | 8 | 1 | -0.50 | 0.04 | 0.60 | 0.0004 |
| p.Arg1036His | 8 | 0 | -0.20 | 0.11 | 0.00 | 0.002 |
| p.Glu1050Lys | 8 | 0 | -0.27 | 0.06 | 0.94 | 0.006 |
| p.Met1052Val | 8 | 0 | -0.68 | 0.53 | 0.01 | 0.006 |
| p.Trp1078Arg | 9 | 0 | -0.69 | 0.66 | 1.00 | 0.687 |
| p.Val1092Ile | 9 | 0 | -0.49 | 0.04 | 0.00 | 0.003 |
| p.Lys1108Thr | 9 | 0 | 0.21 | 0.01 | 0.98 | 0.278 |
| p.Thr1109Ile | 9 | 2 | 0.10 | 0.01 | 1.00 | 0.082 |
| p.Glu1111Lys | 9 | 0 | -0.12 | 0.51 | 1.00 | 0.821 |
| p.Arg1121His | 9 | 0 | 0.09 | 0.33 | 0.04 | 0.004 |
| p.Glu1128Asp | 9 | 0 | 0.27 | 0.89 | 0.01 | 0.011 |
| p.Ile1131Thr | 9 | 5 | -0.18 | 0.03 | 0.54 | 0.003 |
| **p.Arg1138His** | **9** | **2** | **-1.56** | **0.00** | **1.00** | **(-)** |
| **p.Arg1138Cys** | **9** | **0** | **-1.43** | **0.00** | **1.00** | **0.626** |
| **p.Val1139Ile** | **9** | **0** | **-1.39** | **0.13** | **0.30** | **0.052** |
| p.Val1158Ile | 9 | 0 | 0.17 | 0.12 | 0.00 | 0.007 |
| p.Glu1179Lys | C9-C10<br>linker | 3 | NA | NA | NA | NA |
| p.Arg1190His | 10 | 0 | 0.18 | 0.20 | 1.00 | 0.036 |
| p.Ala1194Val | 10 | 1 | 0.29 | 0.08 | 0.95 | 0.005 |
| p.Arg1226Cys | 10 | 0 | 0.09 | 0.00 | 0.99 | 0.012 |

| p.Arg1228Cys | 10 | 1 | -0.27 | 0.02 | 0.99 | 0.020 |
| --- | --- | --- | --- | --- | --- | --- |
| p.Arg1263Trp | 10 | 1 | -0.18 | 0.02 | 0.20 | 0.013 |
| **MYBPC3- Likely Path/Pathogenic (Path) Variants (SHaRe)** |  |  |  |  |  |  |
| **Amino Acid Variant** | **MyBP-C Domain** | **SHaRe Number of Cases** | **STRUM**<br>(deleterious<br>< or = -1.2) | **SIFT**<br>(deleterious<br>< or = 0.05) | **Polyphen-2**<br>(deleterious<br>> 0.90) | **CardioBoost**<br>(deleterious<br>> 0.90)<br>(Benign < 0.10) |
| p.Pro161Ser | 1 | 10 | -0.97 | 0.32 | 0.98 | 0.718 |
| **p.Tyr237Ser** | **1** | **2** | **-4.06** | **0.00** | **1.00** | **0.564** |
| p.Pro371Arg | 2 | 11 | -0.45 | 0.00 | 0.99 | 0.596 |
| p.Arg495Gly | 3 | 13 | -0.11 | 0.25 | 0.87 | 0.767 |
| p.Arg495Trp | 3 | 3 | -0.32 | 0.06 | 0.99 | 0.625 |
| p.Arg495Gln | 3 | 33 | 0.04 | 0.39 | 0.90 | 0.984 |
| p.Arg502Trp | 3 | 67 | -0.23 | 0.01 | 0.48 | 0.934 |
| p.Arg502Gln | 3 | 46 | -0.17 | 0.09 | 0.97 | 0.957 |
| p.Phe503Leu | 3 | 2 | -0.93 | 0.06 | 0.12 | 0.685 |
| p.Gly531Arg | 3 | 13 | -0.55 | 0.05 | 1.00 | 0.378 |
| p.Arg589Gly | 4 | 6 | -0.26 | 0.01 | 0.67 | 0.916 |
| **p.Trp792Arg** | **6** | **16** | **-1.28** | **0.00** | **1.00** | **0.990** |
| **p.Arg810His** | **6** | **18** | **-1.22** | **0.06** | **1.00** | **0.984** |
| p.Arg810Leu | 6 | 6 | -1.12 | 0.07 | 1.00 | 0.983 |
| p.Arg820Gln | 6 | 4 | 0.04 | 0.16 | 0.97 | 0.956 |
| **p.Ala848Glu** | **6** | **4** | **-2.33** | **0.01** | **1.00** | **0.425** |
| p.Ser858Asn | 6 | 7 | -0.73 | 0.01 | 1.00 | 0.685 |
| **p.Leu1238Pro** | **10** | **1** | **-2.89** | **0.00** | **1.00** | **0.996** |
| **p.Asn1257Lys** | **10** | **3** | **-1.45** | **0.00** | **1.00** | **0.374** |

NA indicates variants not amenable to analysis by STRUM due to be located outside domains analyzed by STRUM
(Supplemental Table 1). If a variant was not amenable to analysis by STRUM it was not analyzed by other methods (also
denoted in NA). A subset of variants did not yield a result in CardioBoost and were also indicated by (-). MYBPC3
HGNC:7551, NP\_000247.2, NM\_000256.3.

**Supplemental Table 2:**
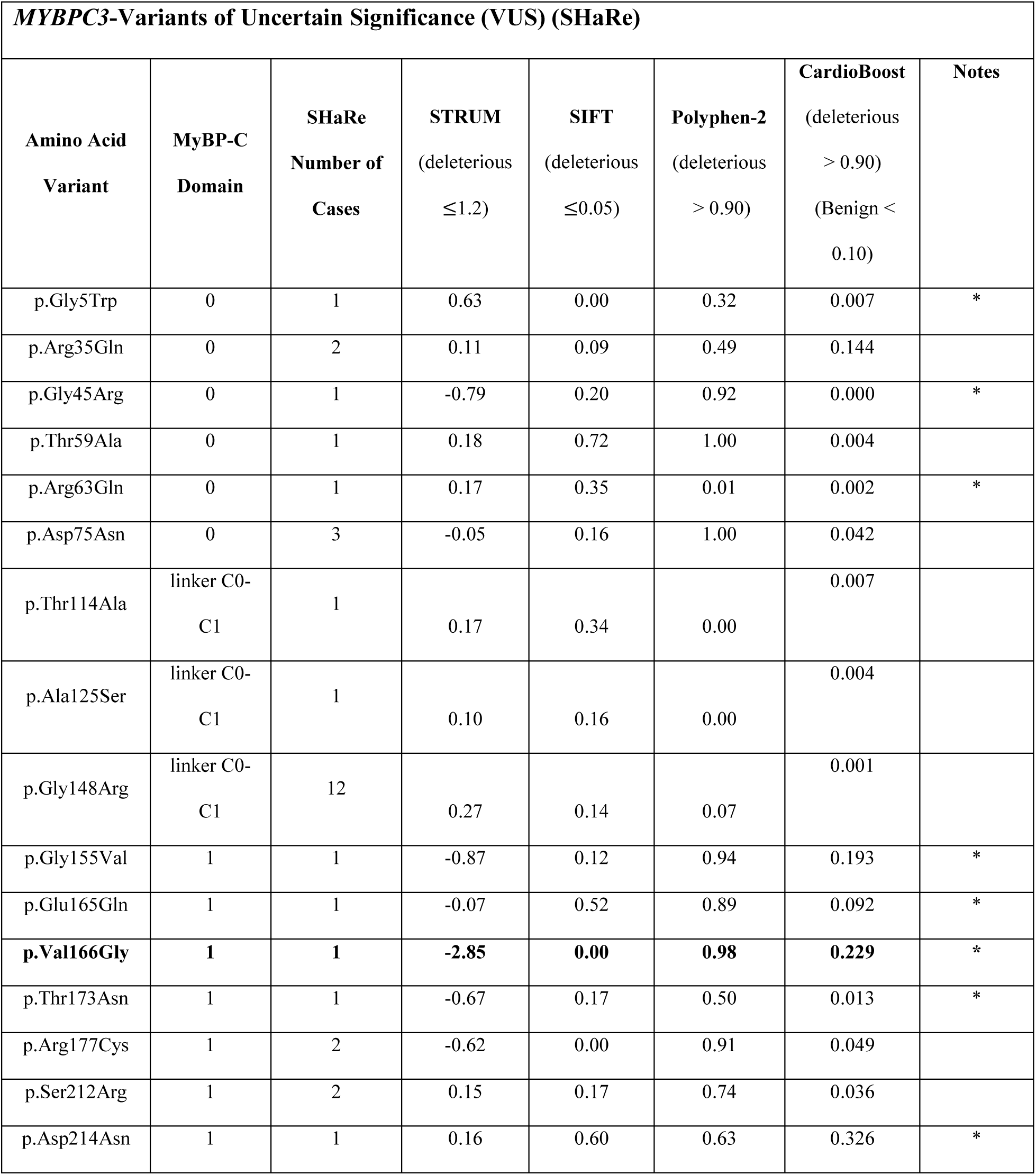
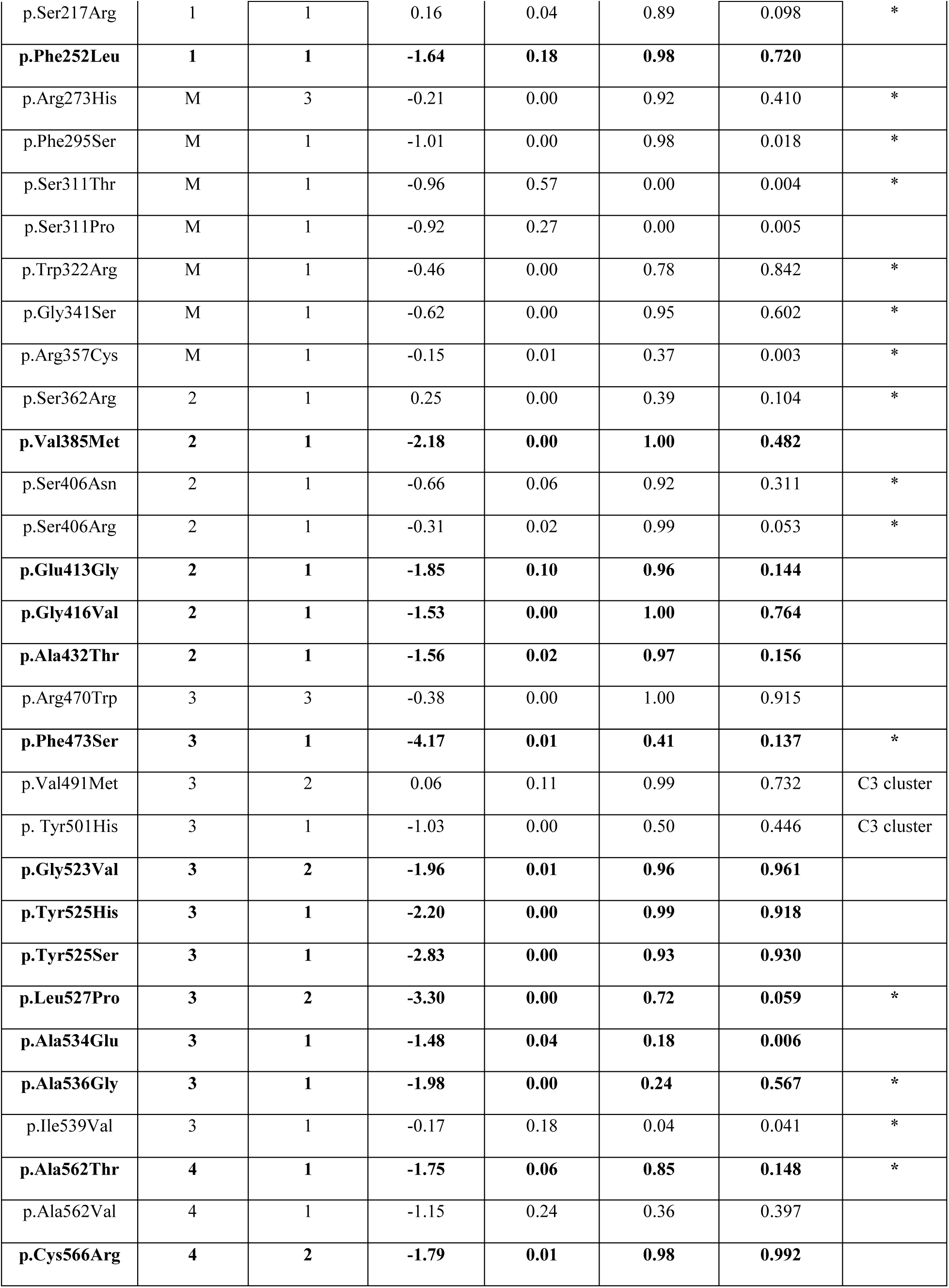
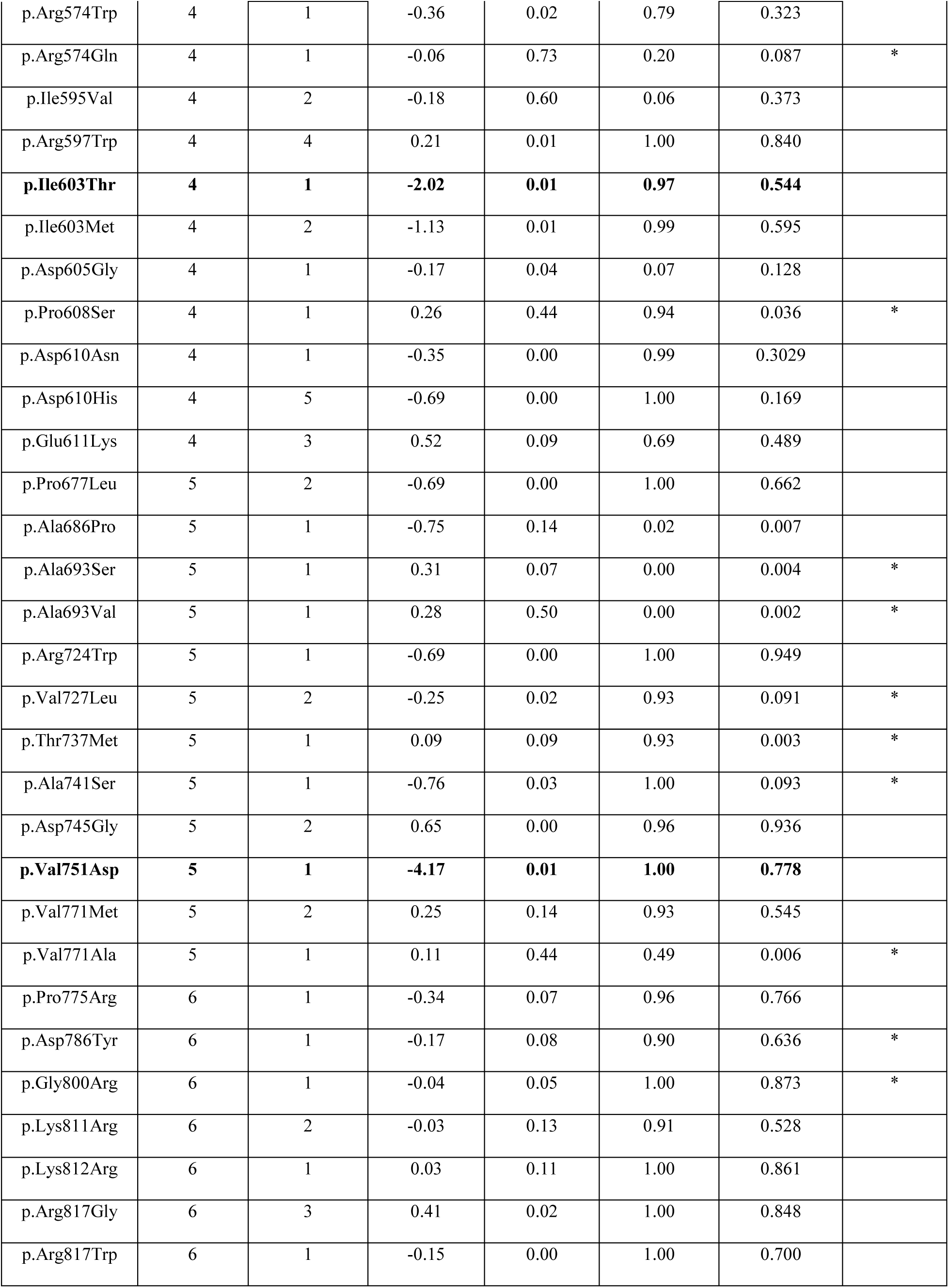
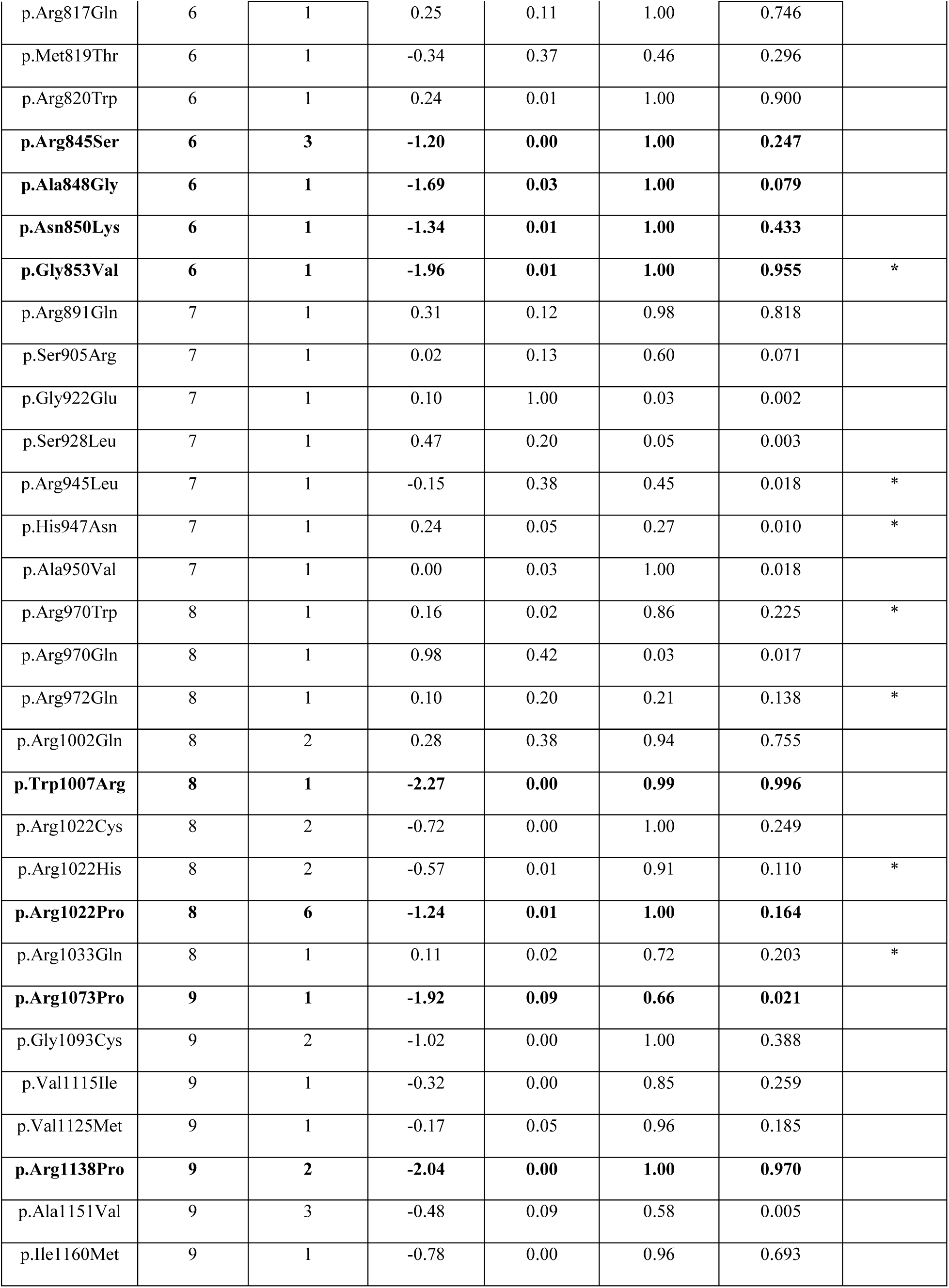
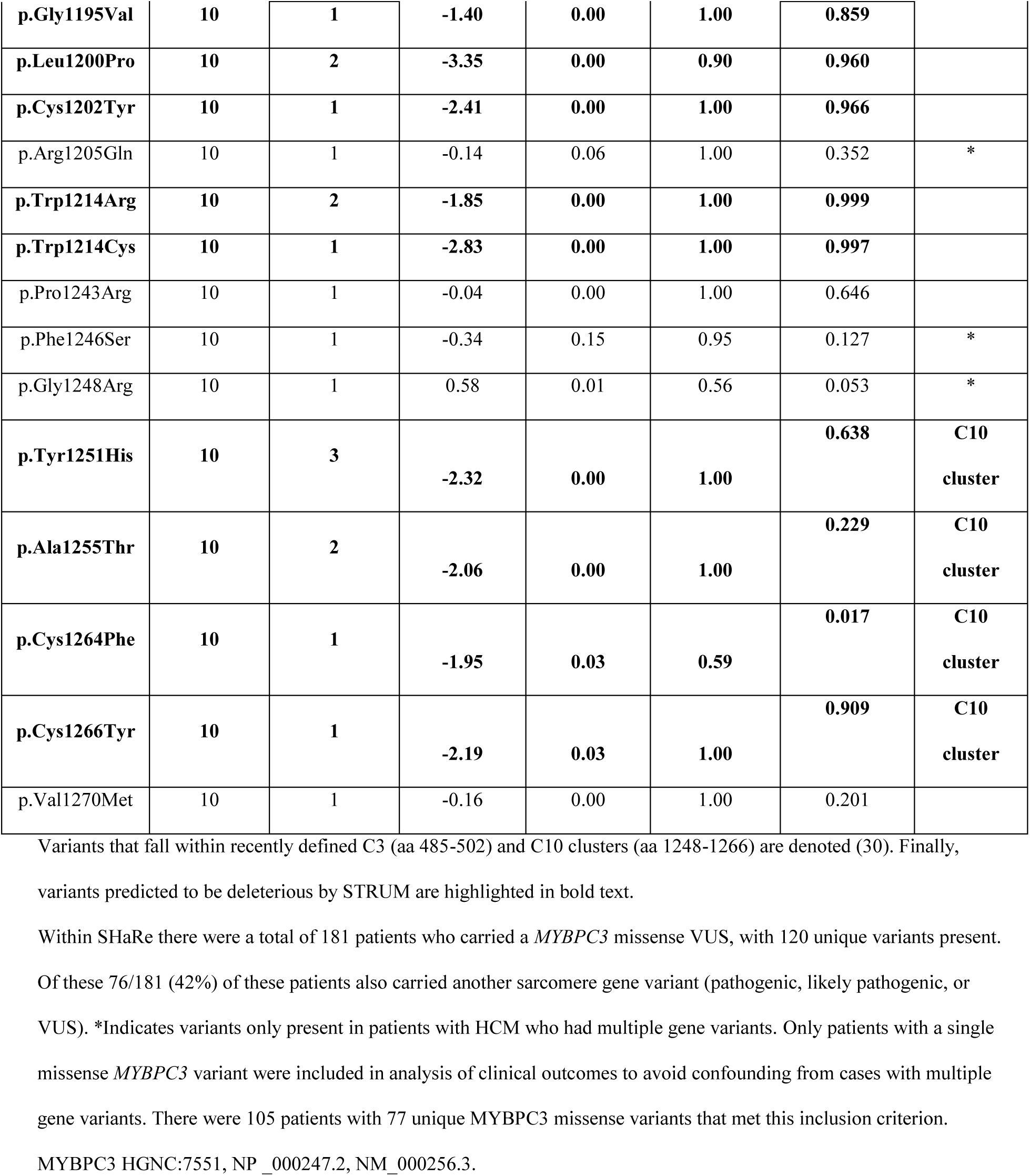
Computation analysis of *MYBPC3* non-synonymous missense variants-VUS

**Supplemental Table 3:** Summary of STRUM inputs and resulting I-TASSER models

| MyBP-C Domain | Model Input Sequence | Domain Type | STRUM A.A. | Available Structures | C-score | Estimated TM score | RMSD |
| --- | --- | --- | --- | --- | --- | --- | --- |
| C0 | 2 to 96 | Ig-like C2 | 2-96 | 2K1M (NMR) (18) | 0.32 | 0.76+/-0.10 | 0.29 |
| C1 | 153 to 256 | Ig-like C2 | 153-256 | 2AVG (NMR) (17) | 0.22 | 0.74+/- 0.11 | 0.44 |
|  |  |  |  | 3CX2 (Xray) (20) |  |  | 1.35 |
|  |  |  |  | 2V6H (Xray) (19) |  |  | 1.35 |
| C2 | 362 to 452 | Ig-like C2 | 362-452 | 1PD6 (NMR)(16) | 0.42 | 0.77+/-0.10 | 0.30 |
|  |  |  |  | 5K6P (NMR) (21) |  |  | 1.99 |
| C3 | 453-543 | Ig-like C2 | 453-543 | 2MQ0 (NMR)(22) | 0.59 | 0.79+/- 0.09 | 0.40 |
| C4 | 542-633 | Ig-like C2 | 544-633 | none | 0.4 | 0.77+/-0.10 | NA |
| C5 | 645-771 | Ig-like C2 | 645-771 | 1GXE (NMR) (23) | 0.34 | 0.76+/- 0.10 | 0.48 |
| C6 | 774-870 | Fibronectin | 774-870 | none | 0.90 | 0.84+/- 0.08 | NA |
| C7 | 872-967 | Fibronectin | 872-967 | none | 0.98 | 0.85+/-0.08 | NA |
| C8 | 970-1065 | Ig-like C2 | 970-1065 | none | 0.44 | 0.77+/-0.10 | NA |
| C9 | 1068-1163 | Fibronectin | 1068-1163 | none | 0.60 | 0.79+/- 0.09 | NA |
| C10 | 1181-1274 | Ig-like C2 | 1181-1274 | none | 0.77 | 0.82+/-0.09 | NA |
| C0-PR-C1 | 1 to 260 | NA | 97-152 | 2K1M (NMR ) | -2.01 | 0.47+/-0.15 | 1.39 |
|  |  |  |  | 2AVG (NMR) |  |  | 1.61 |
|  |  |  |  | 3CX2 (Xray) |  |  | 1.41 |
|  |  |  |  | 2V6H (Xray) |  |  | 1.20 |
| C1-M-C2 | 152 to 452 | NA | 257-361 | 2AVG (NMR) | -0.72 | 0.62+/-0.14 | 2.02 |
|  |  |  |  | 3CX2 (Xray) |  |  | 1.07 |
|  |  |  |  | 2V6H (Xray) |  |  | 1.19 |
|  |  |  |  | 1PD6 (NMR) |  |  | 2.78 |
|  |  |  |  | 5K6P (NMR) |  |  | 3.38 |

The C-score range is from [-5,2] where a C-score of higher value signifies a model with high confidence. Another measure of model quality is TM-score, that is sensitive to local error. A TM-score > 0.5 indicates a model of correct topology. Our models exhibit TM-score of 0.74- 0.84. Finally, we report the root mean square deviation (RMSD) in Angstroms comparing I-TASSER models to known *MYBPC3* subdomain structures. STRUM A.A. indicates the amino acids (A.A) whom utilized the a given model input sequence to execute the STRUM analysis.

**Supplemental Table 4.** STRUM results by subdomain

| MyBP-C | | $\Delta\Delta G$ kcal/mol | | Variants | % deleterious |
| --- | --- | --- | --- | --- | --- |
| Domain | Domain Type | Mean | Stdev | Analyzed | $\Delta\Delta G \leq -1.2$ kcal/mol |
| C0 | IgG-like C2 | -0.37 | 0.90 | 1805 | 15.79 |
| PR | linker | -0.04 | 0.47 | 1120 | 2.50 |
| C1 | IgG-like C2 | -0.78 | 0.96 | 2080 | 25.34 |
| M domain | linker | -0.51 | 0.74 | 2100 | 17.05 |
| C2 | IgG-like C2 | -1.11 | 0.99 | 1820 | 35.49 |
| C3 | IgG-like C2 | -0.59 | 0.89 | 1820 | 19.51 |
| C4 | IgG-like C2 | -0.50 | 0.87 | 1800 | 16.78 |
| C5 | IgG-like C2 | -0.43 | 1.01 | 2540 | 16.50 |
| C6 | Fibronectin | -0.53 | 0.95 | 1940 | 19.48 |
| C7 | Fibronectin | -0.34 | 0.89 | 1920 | 15.83 |
| C8 | IgG-like C2 | -0.79 | 1.06 | 1920 | 26.61 |
| C9 | Fibronectin | -0.69 | 1.09 | 1920 | 22.40 |
| C10 | IgG- like C2 | -0.58 | 0.93 | 1880 | 21.28 |

**Supplemental Table 5:**
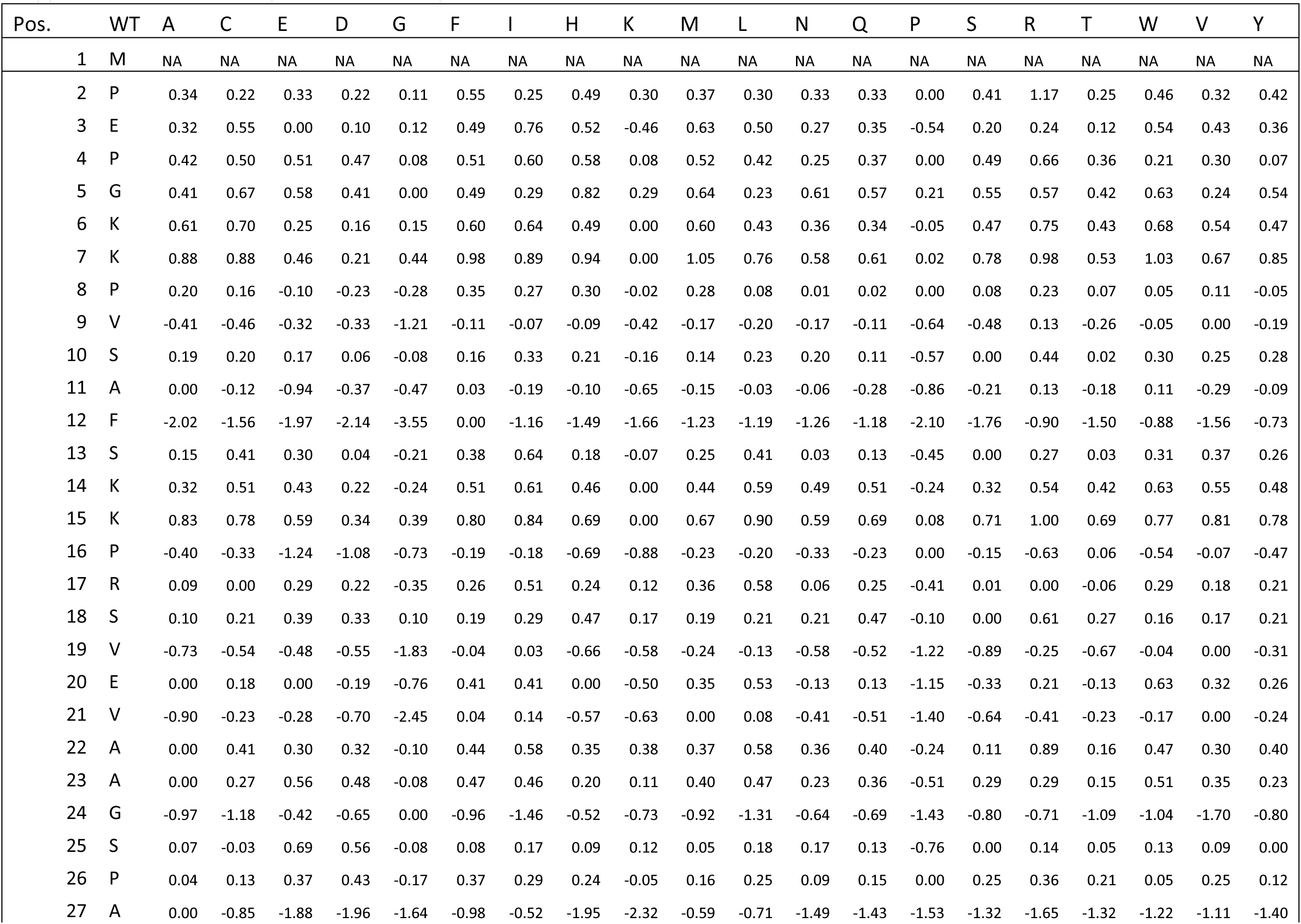
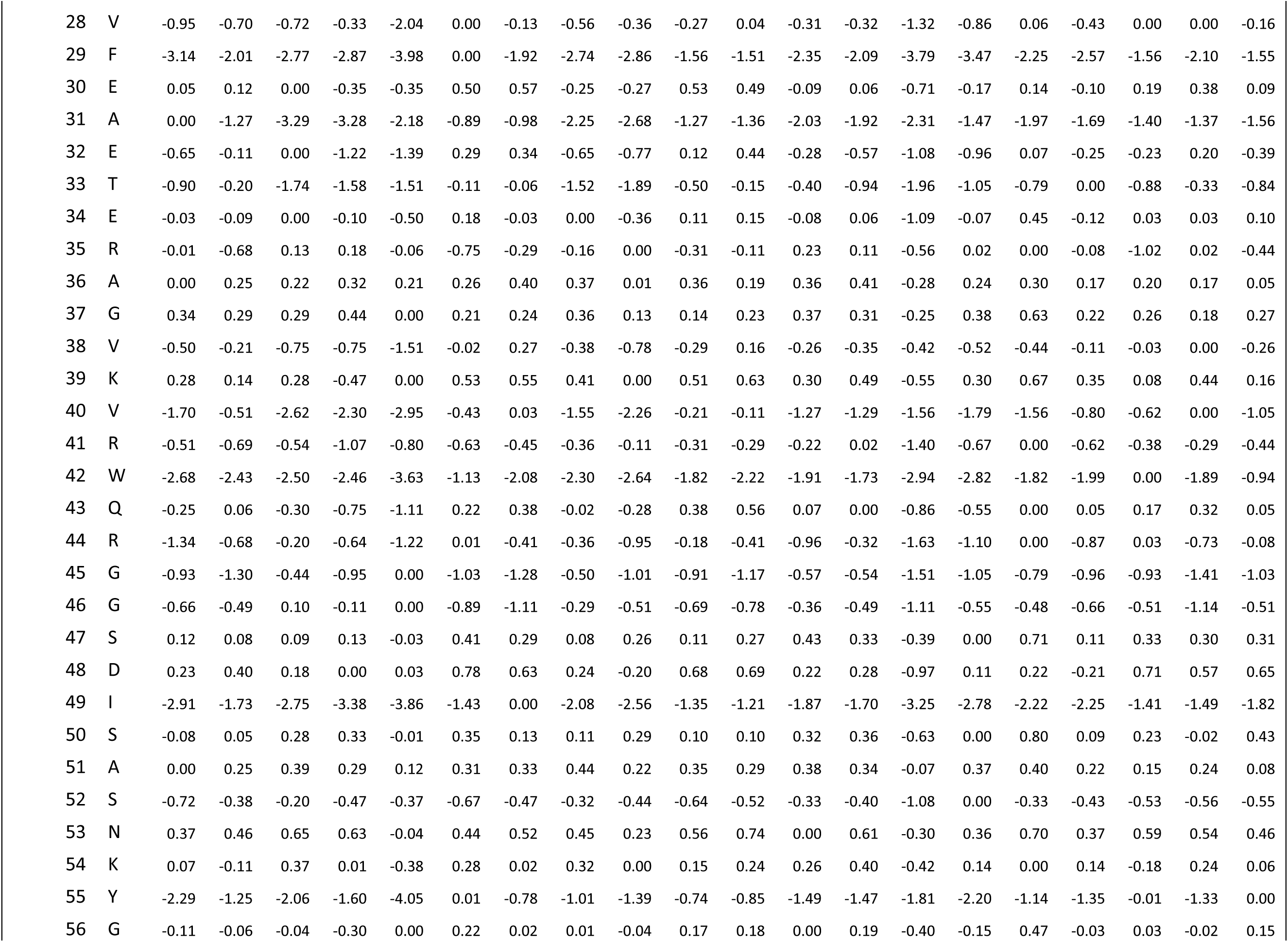
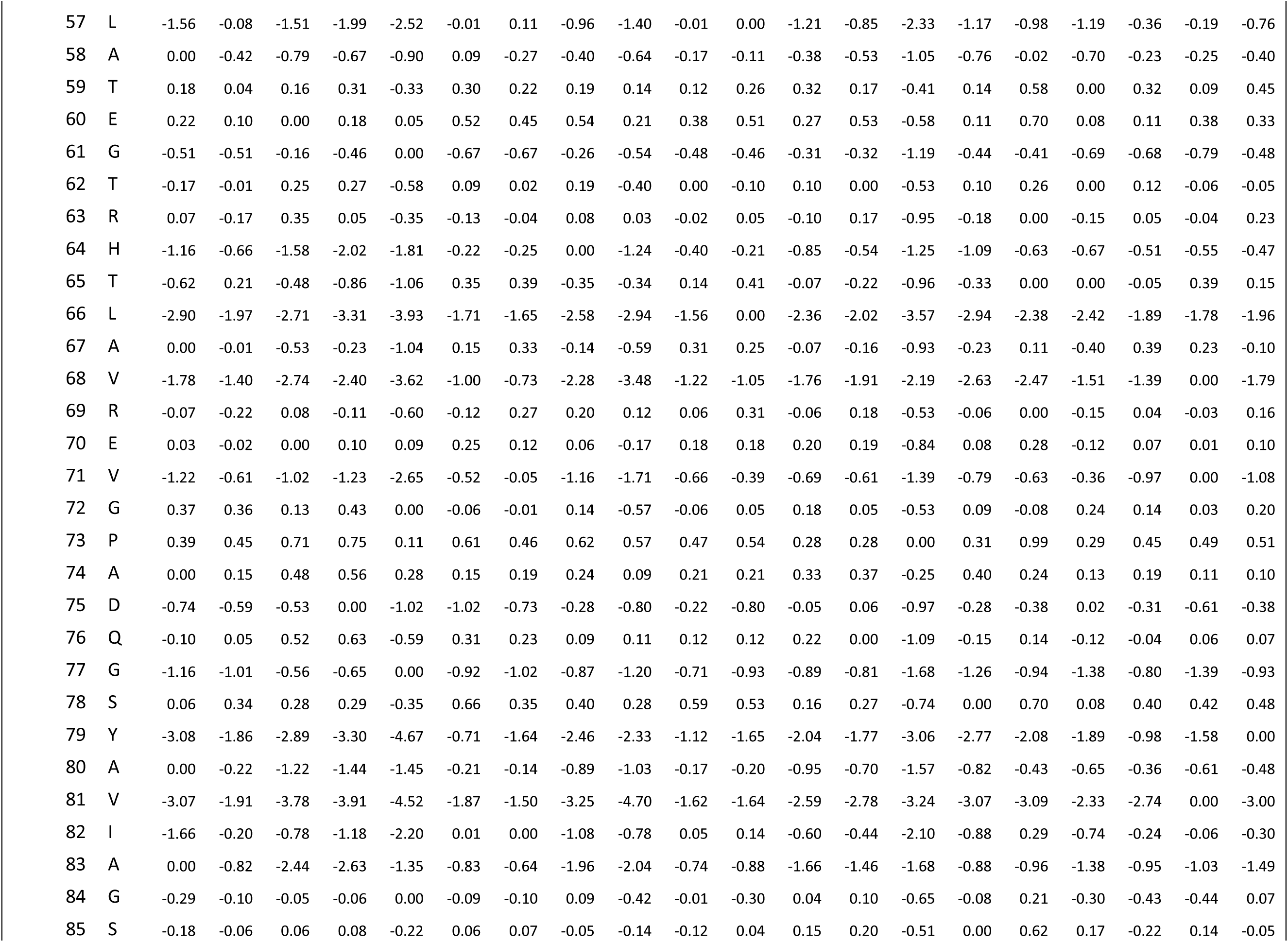
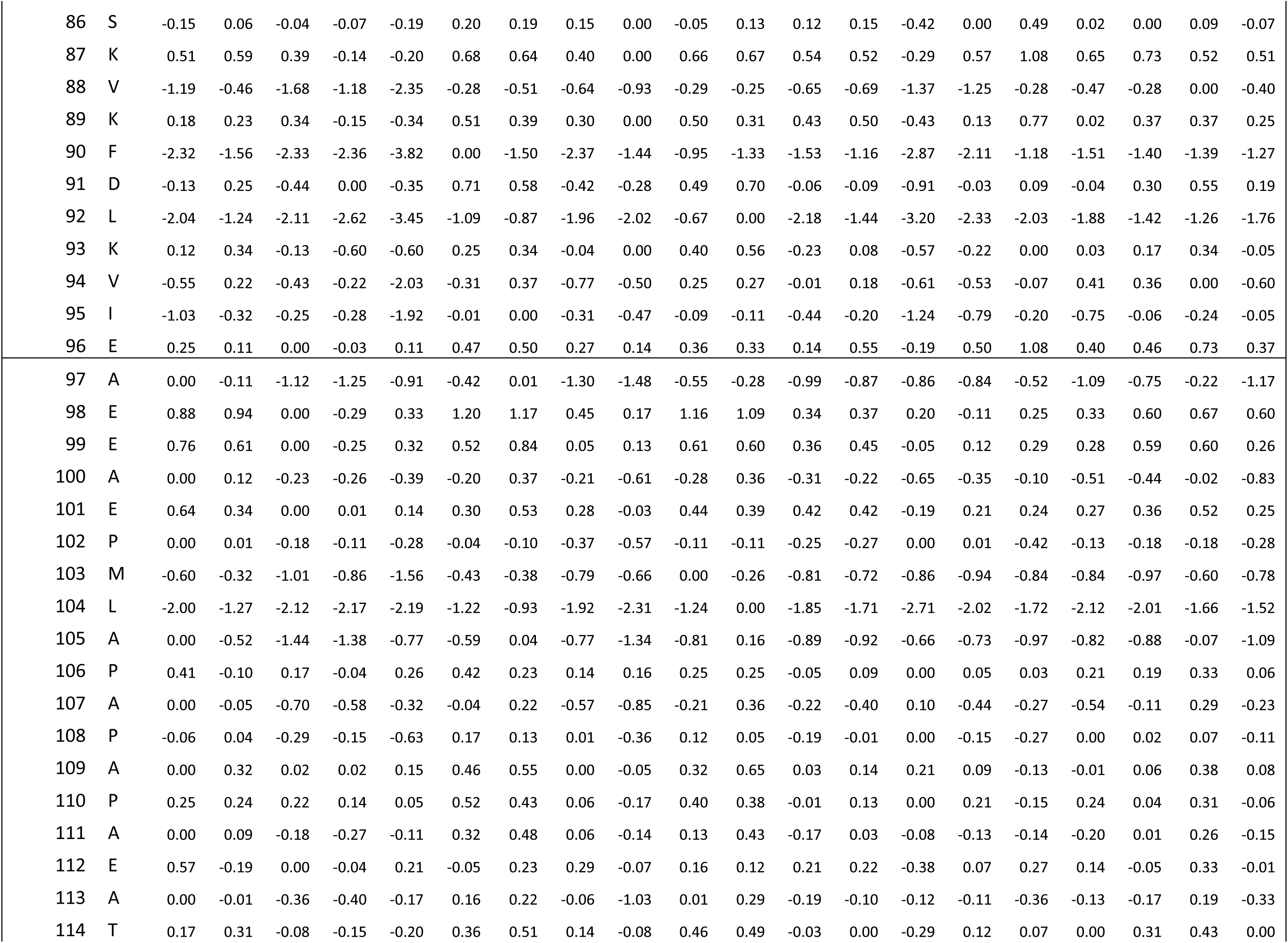
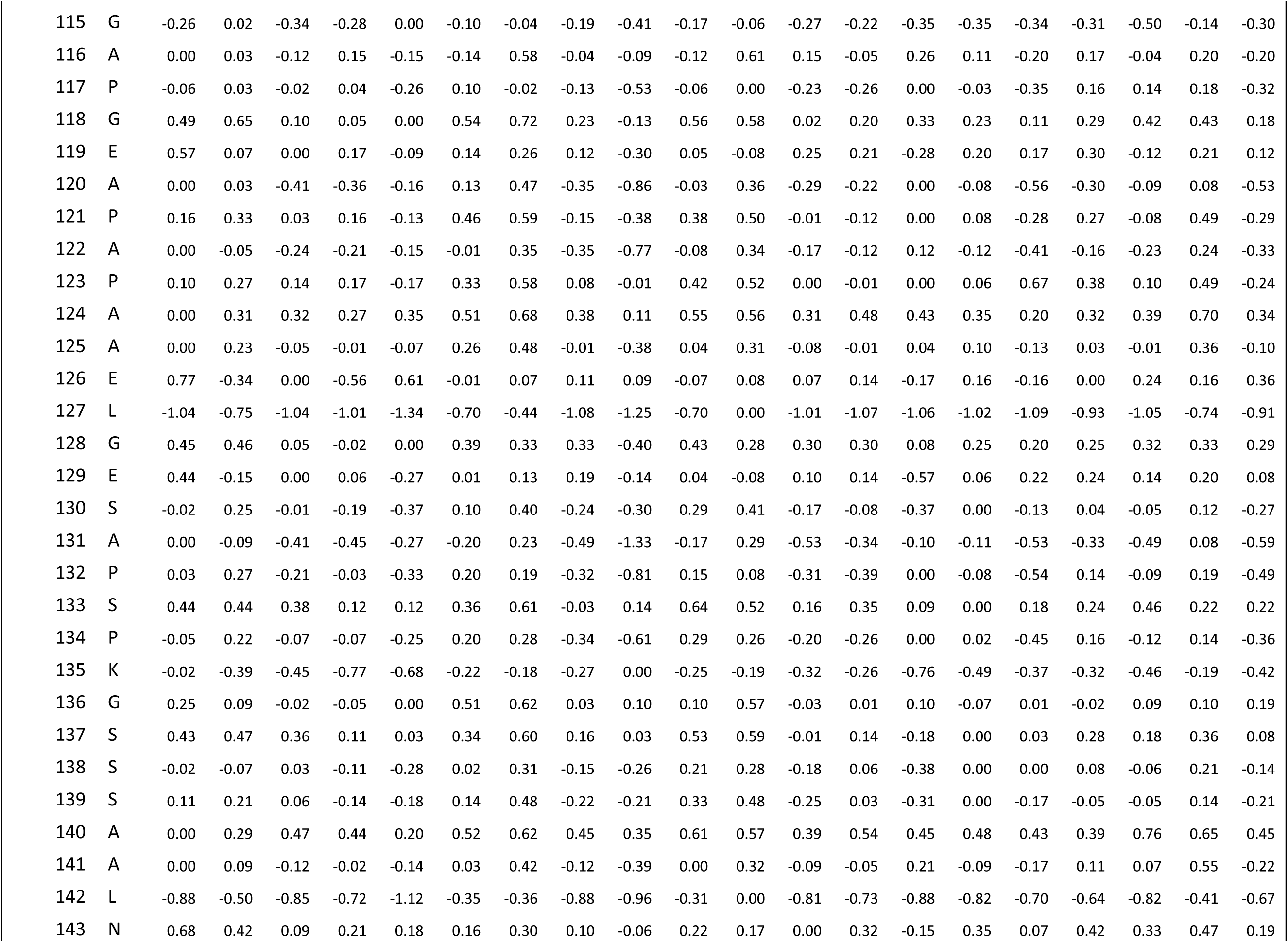
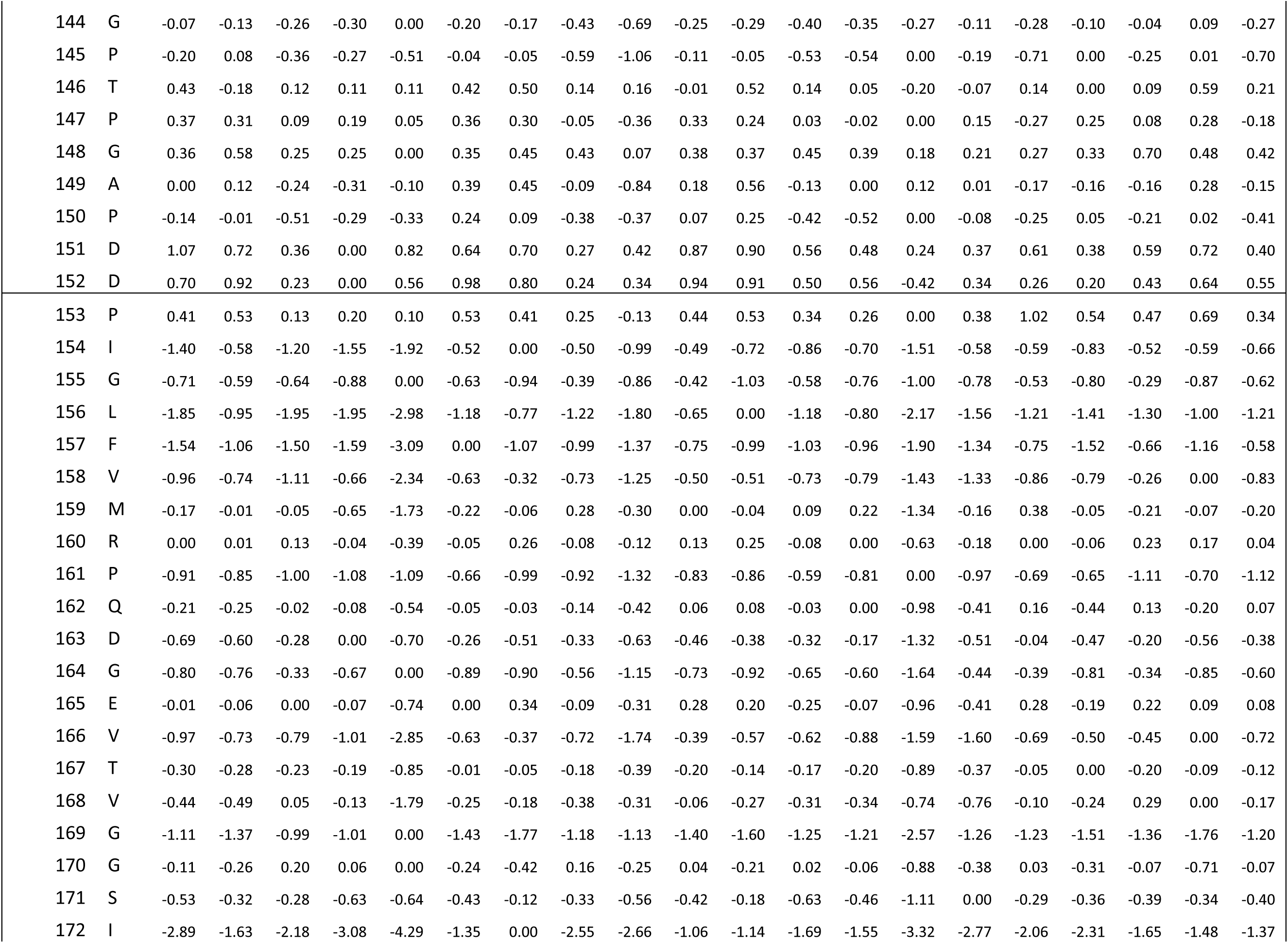
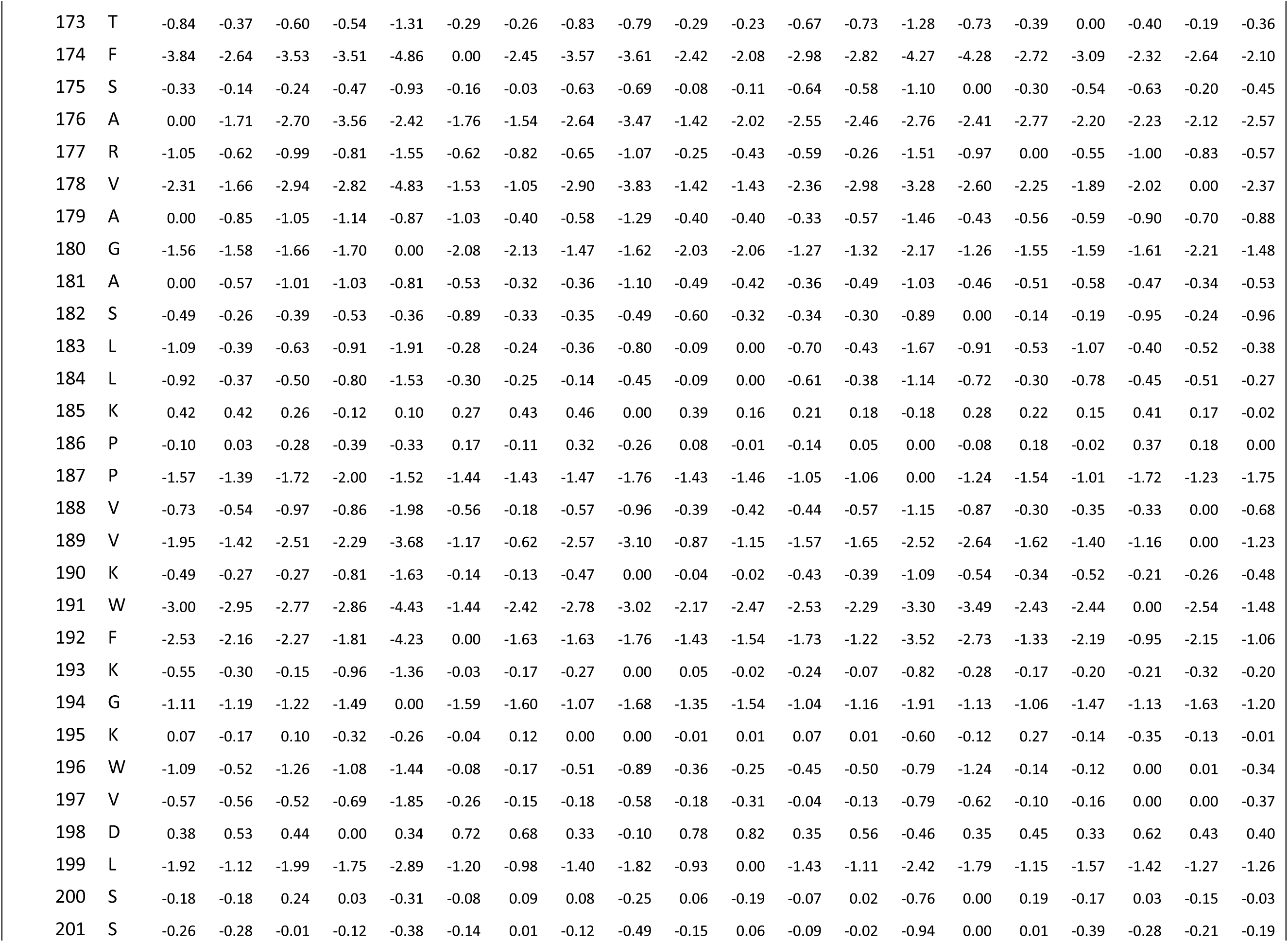
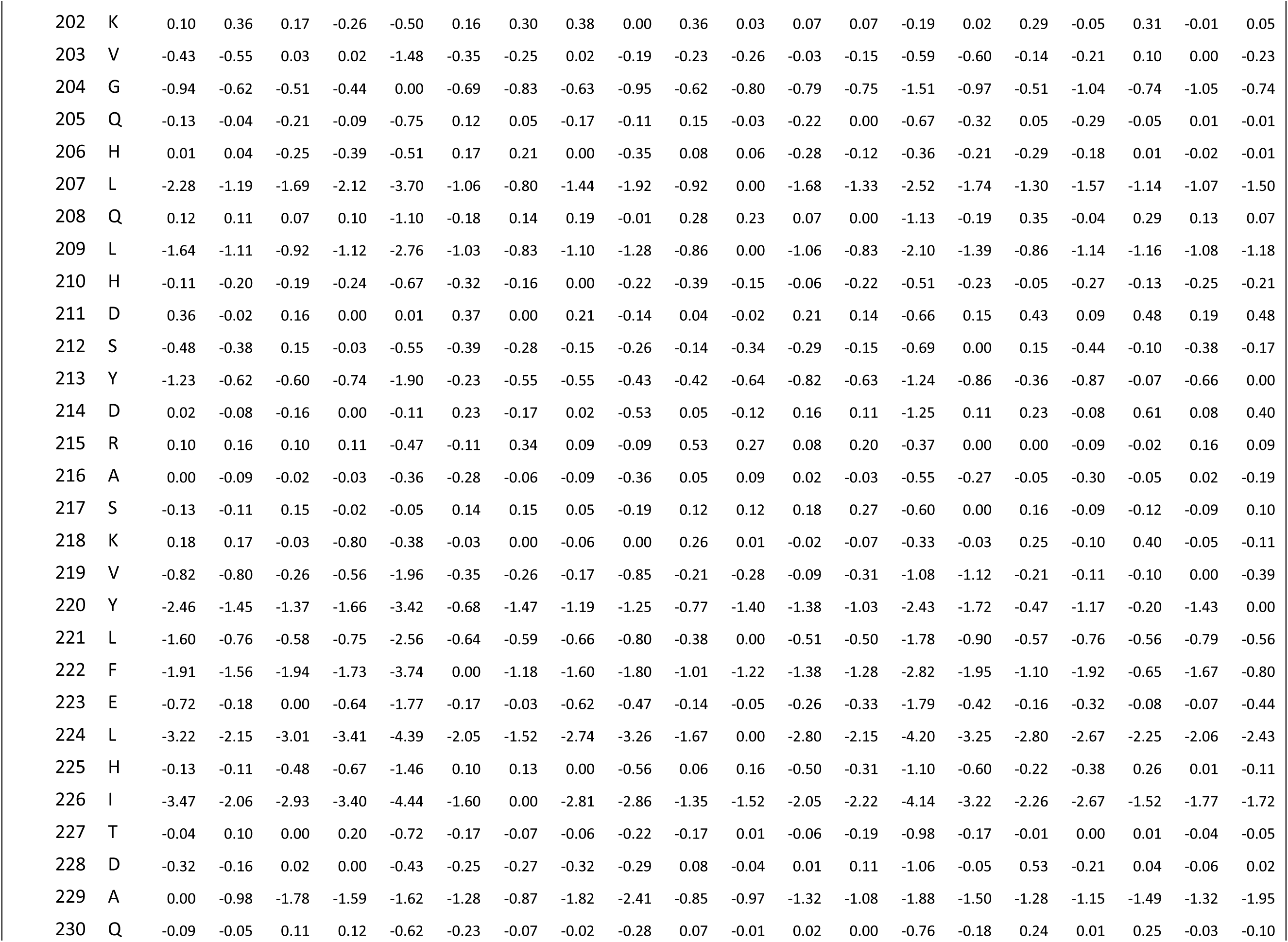
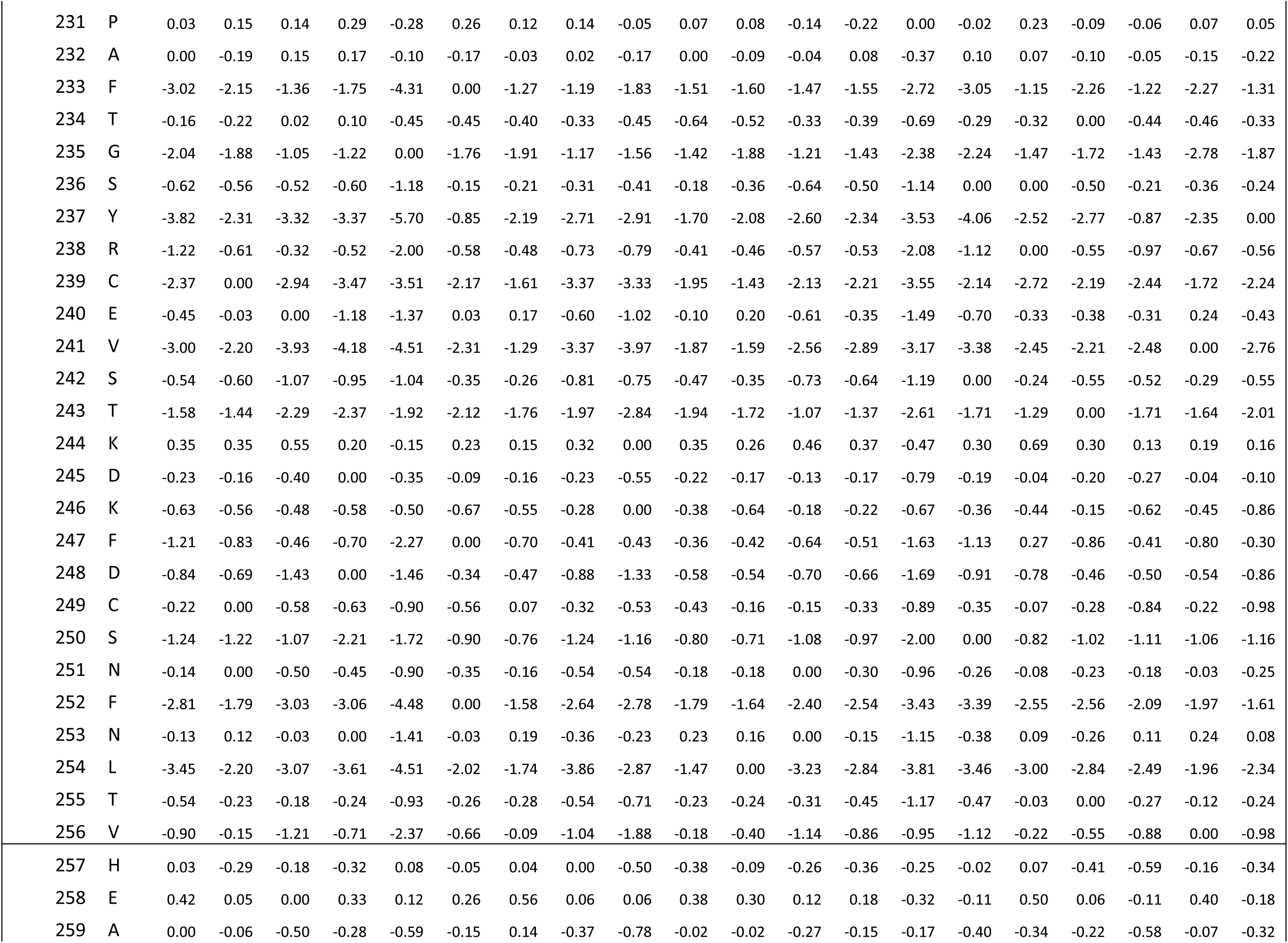
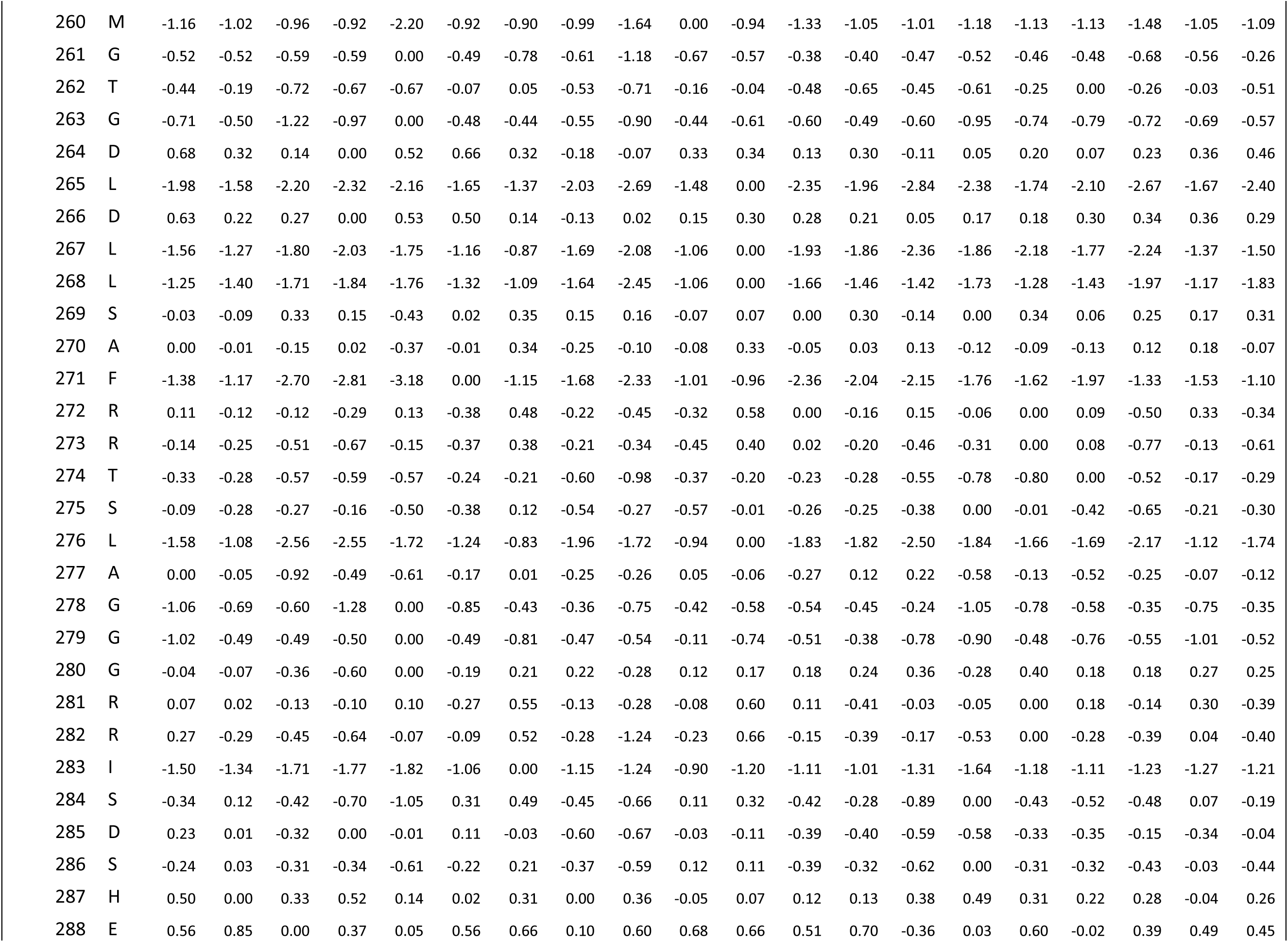
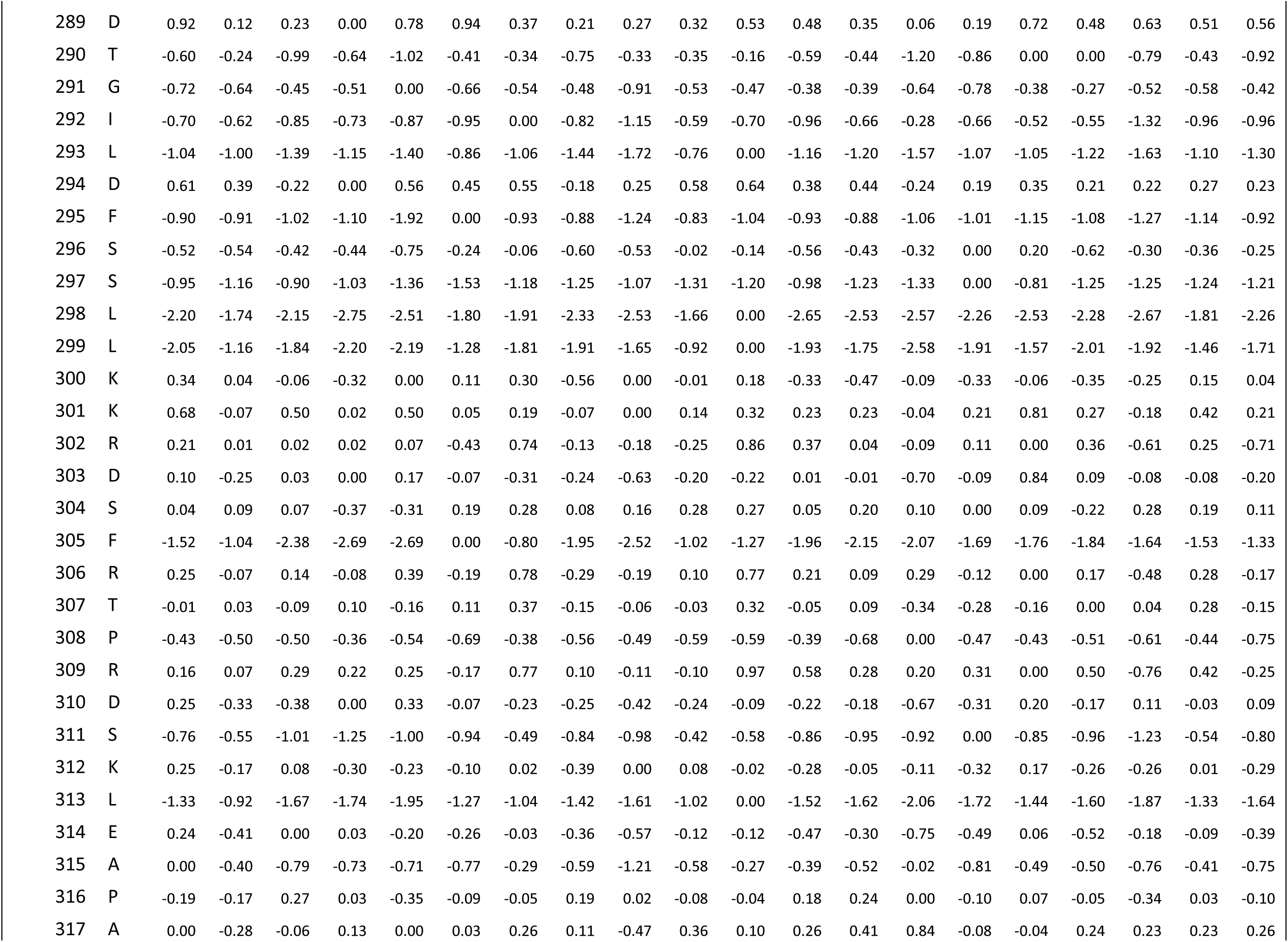
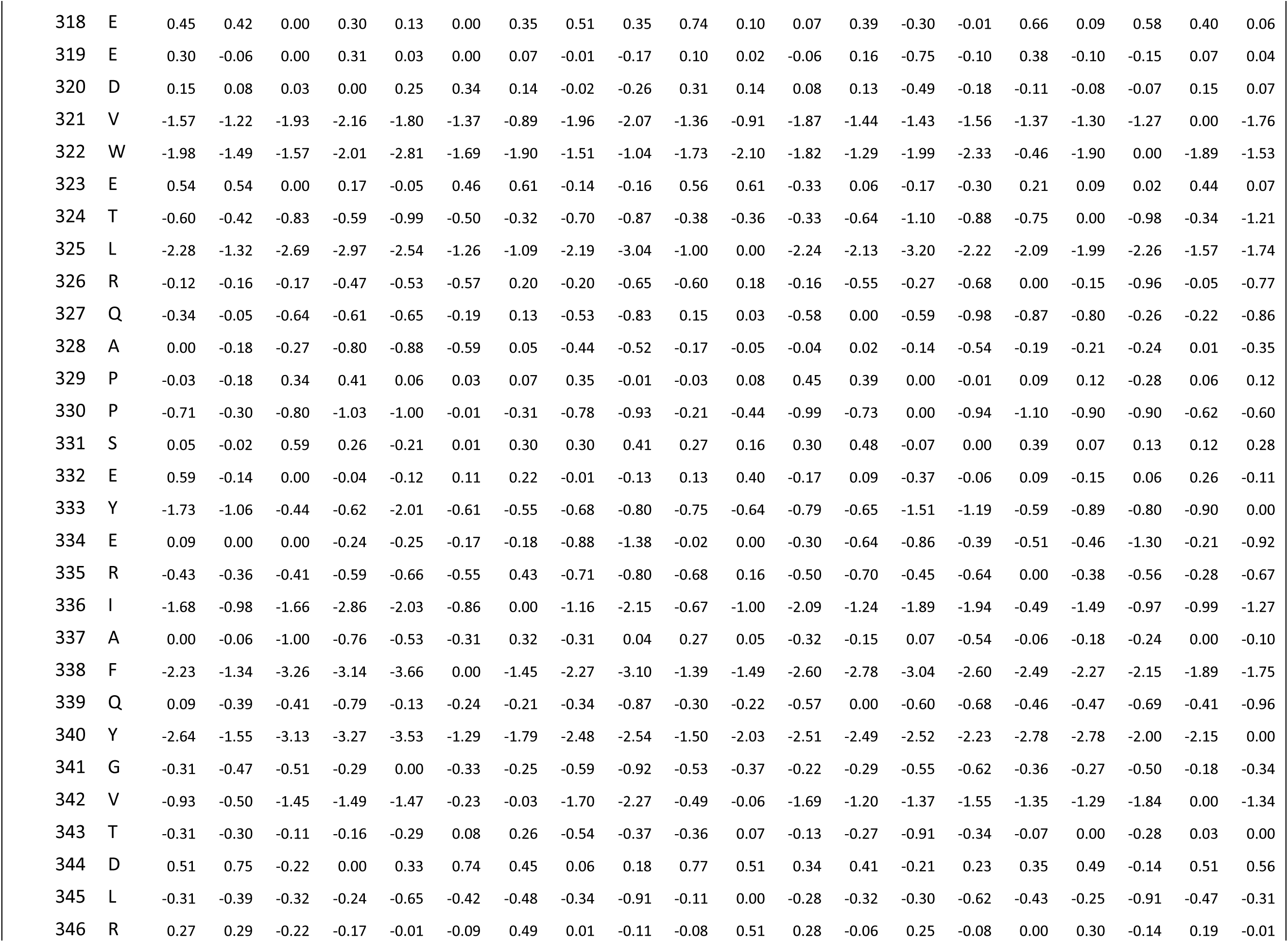
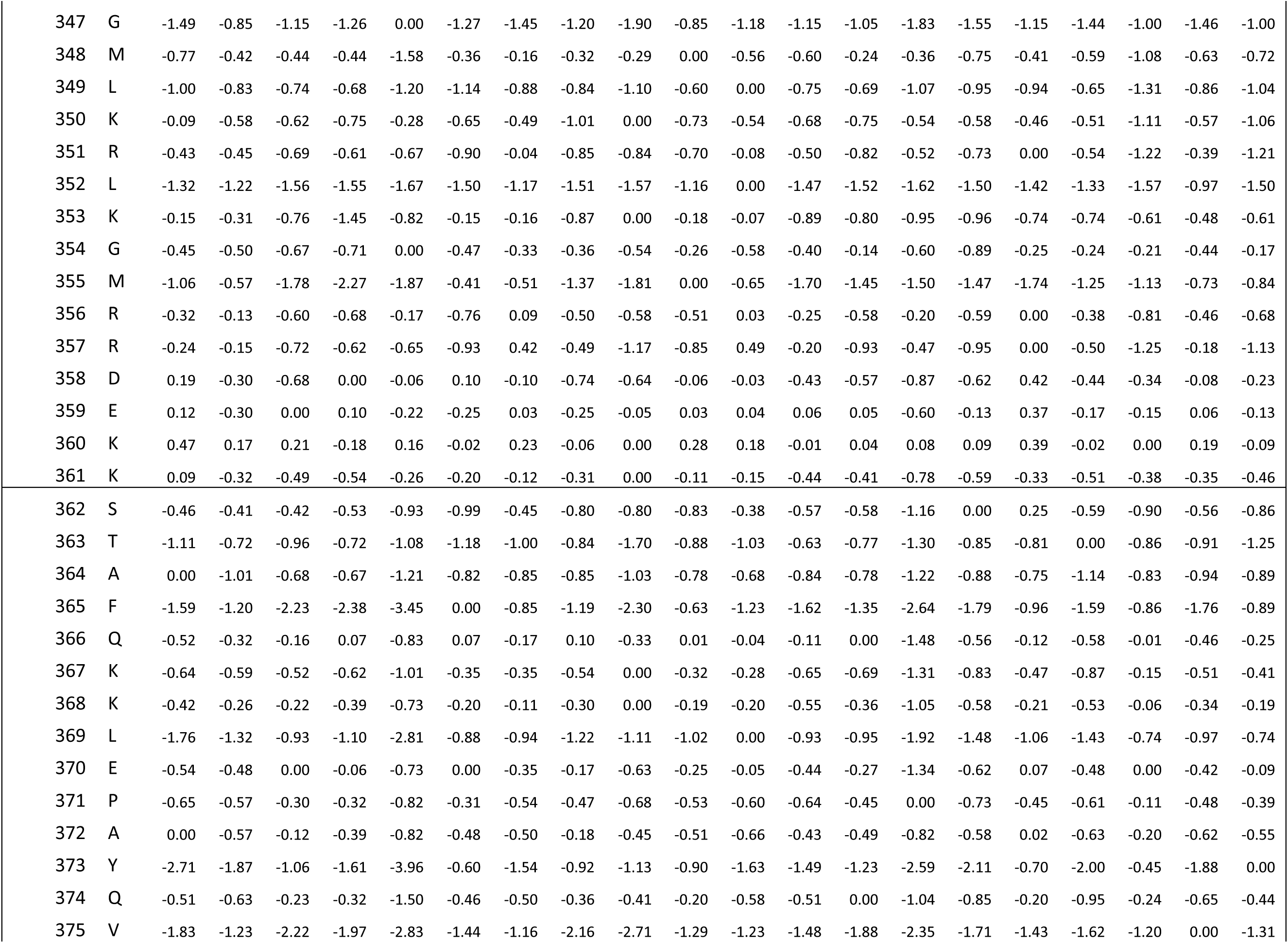
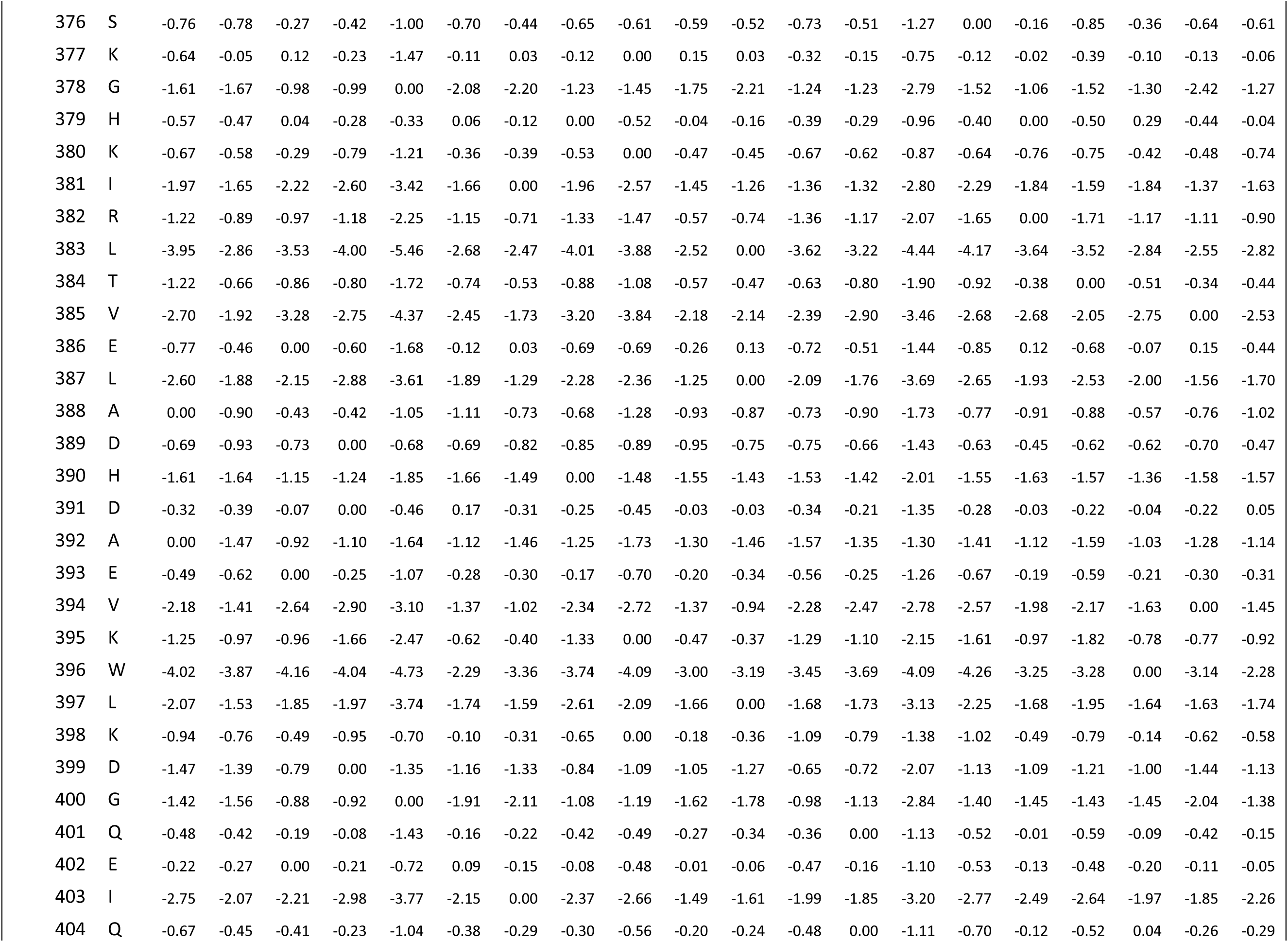
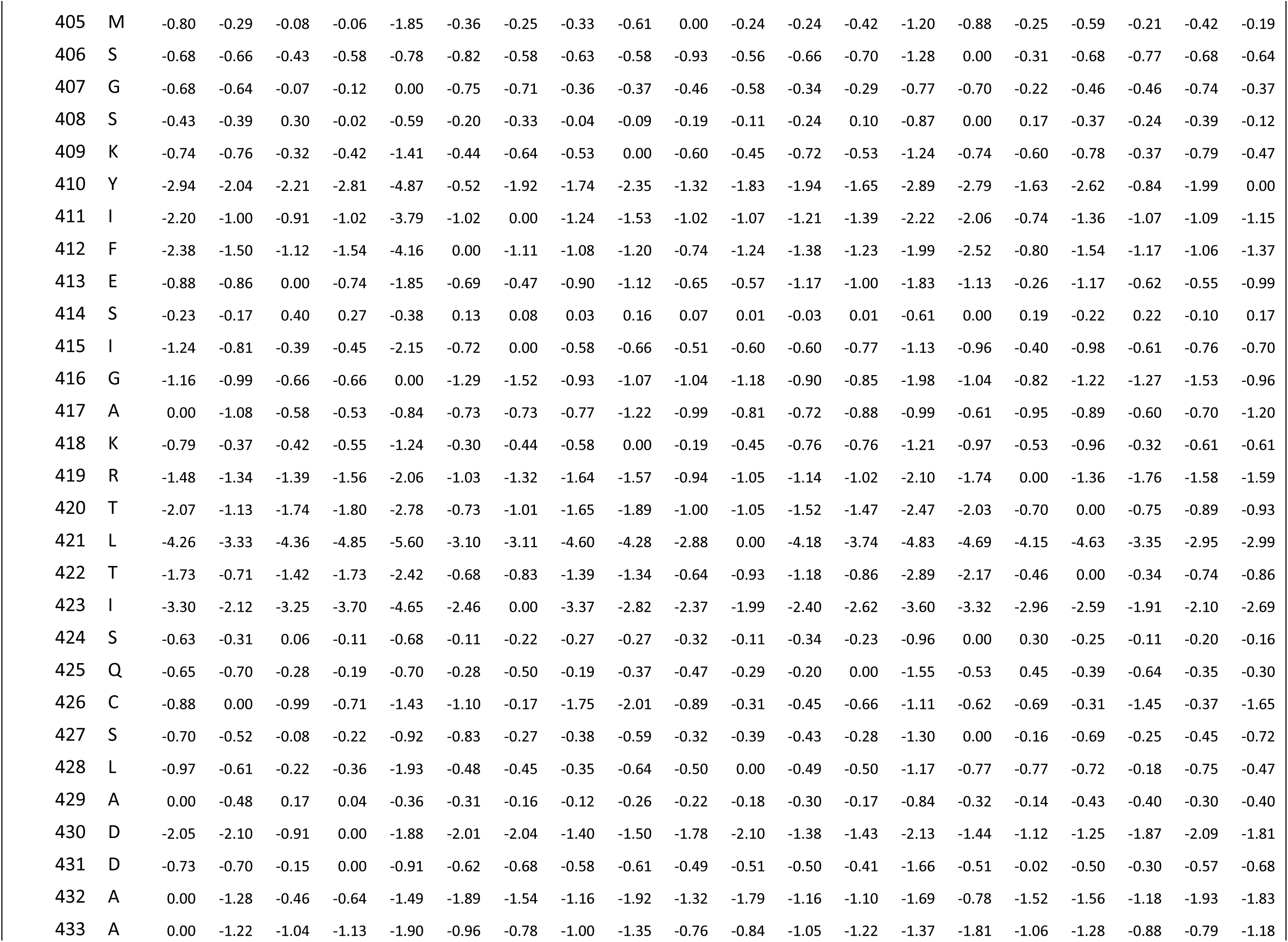
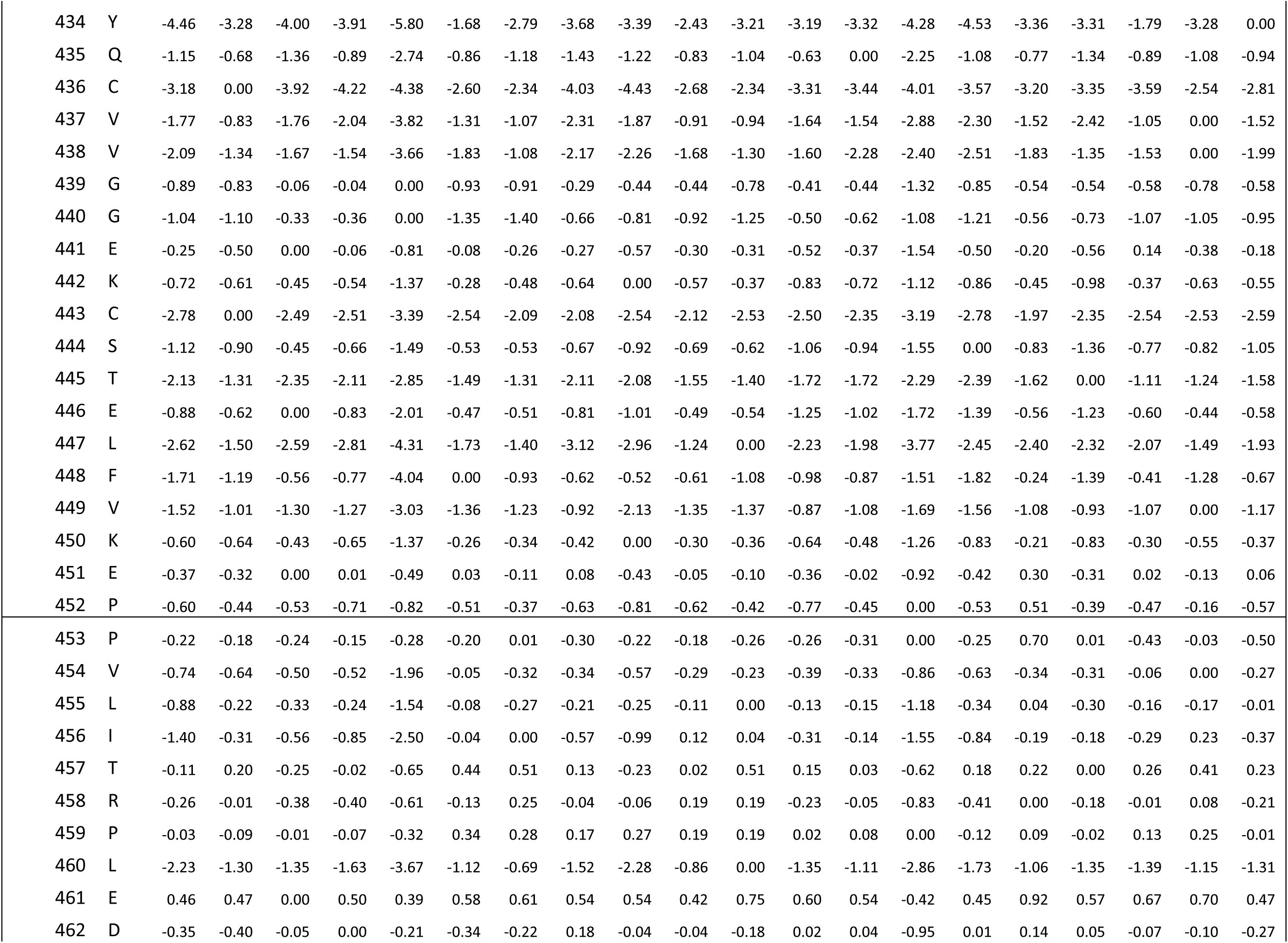
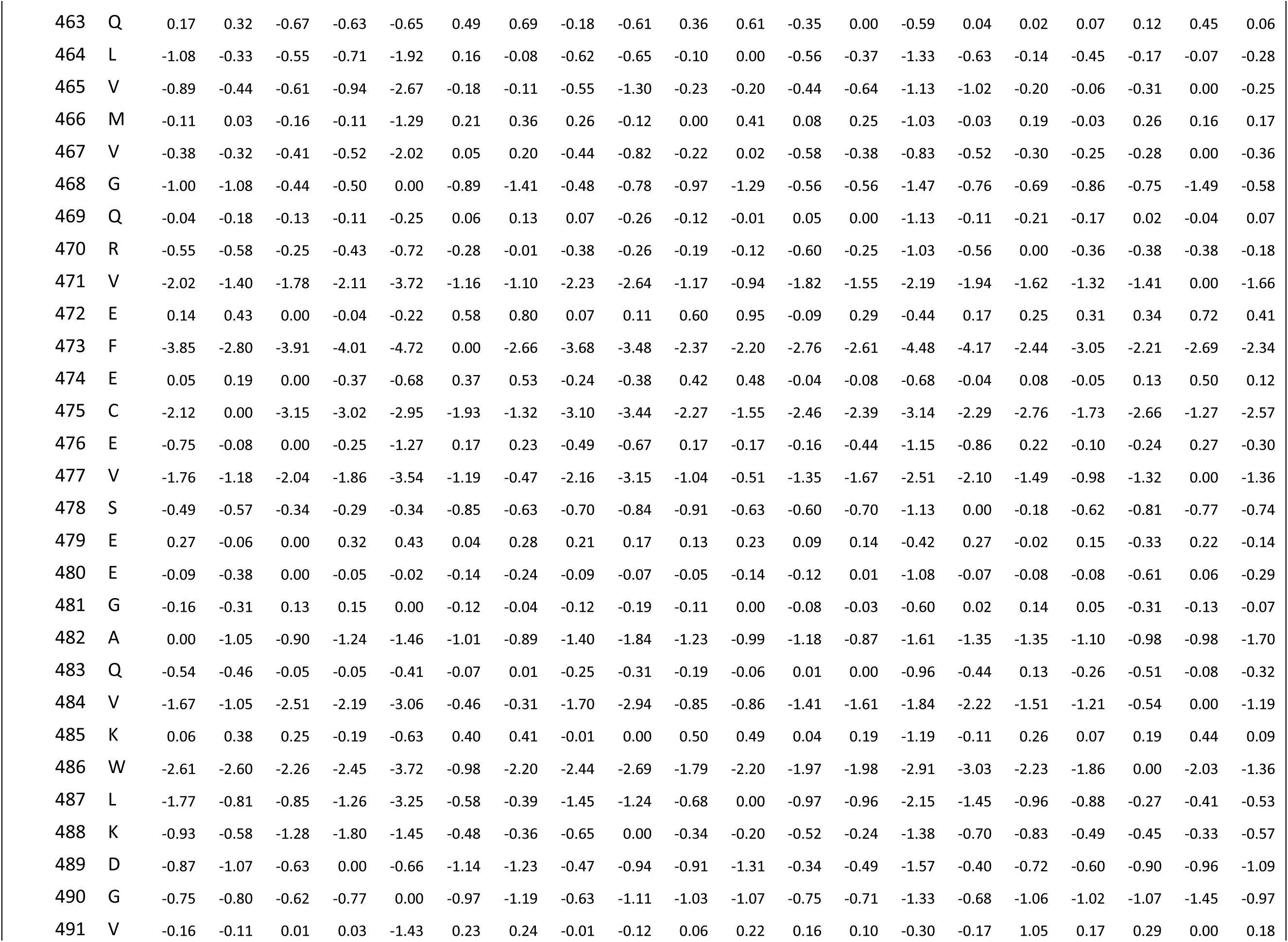
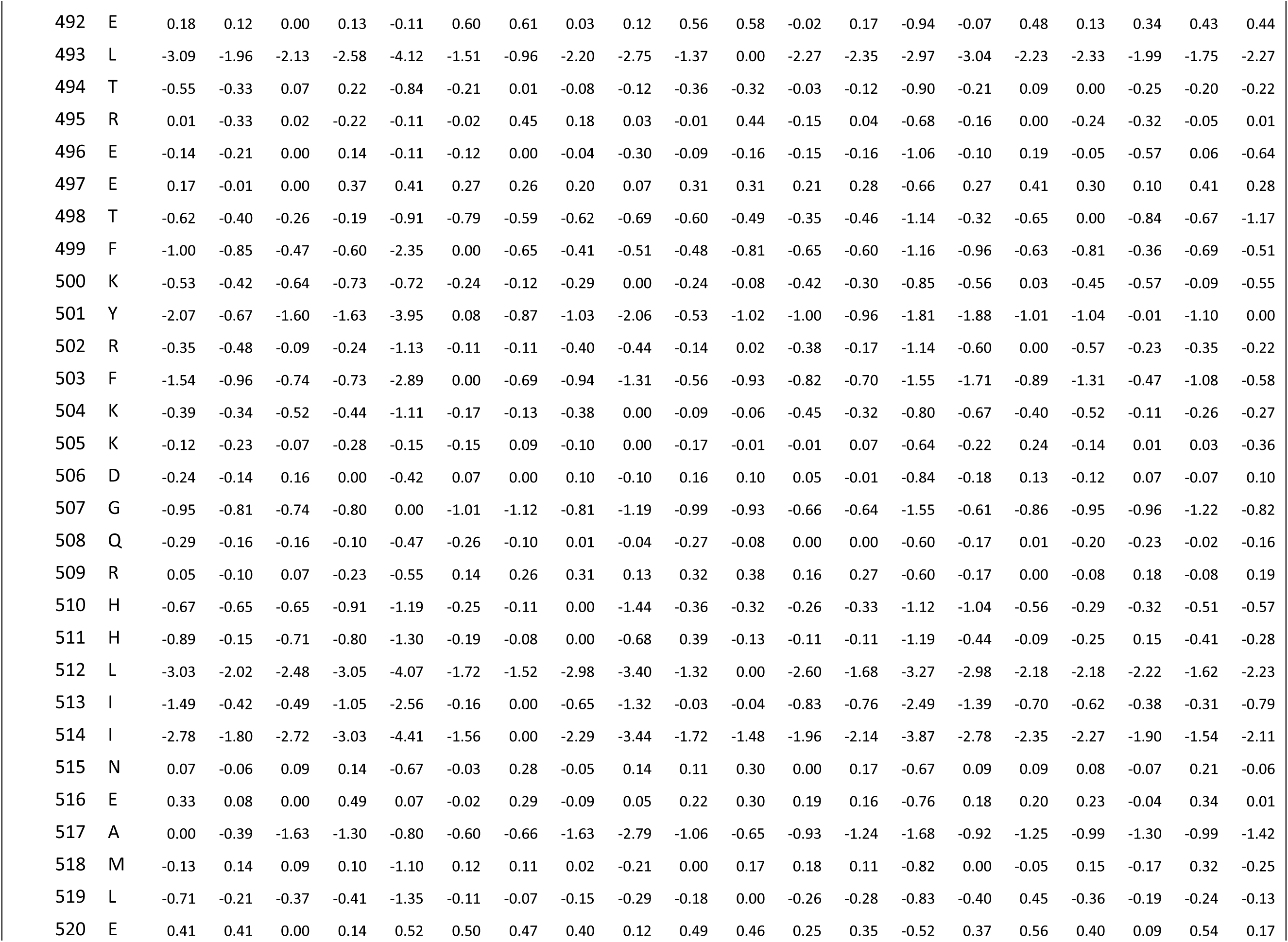
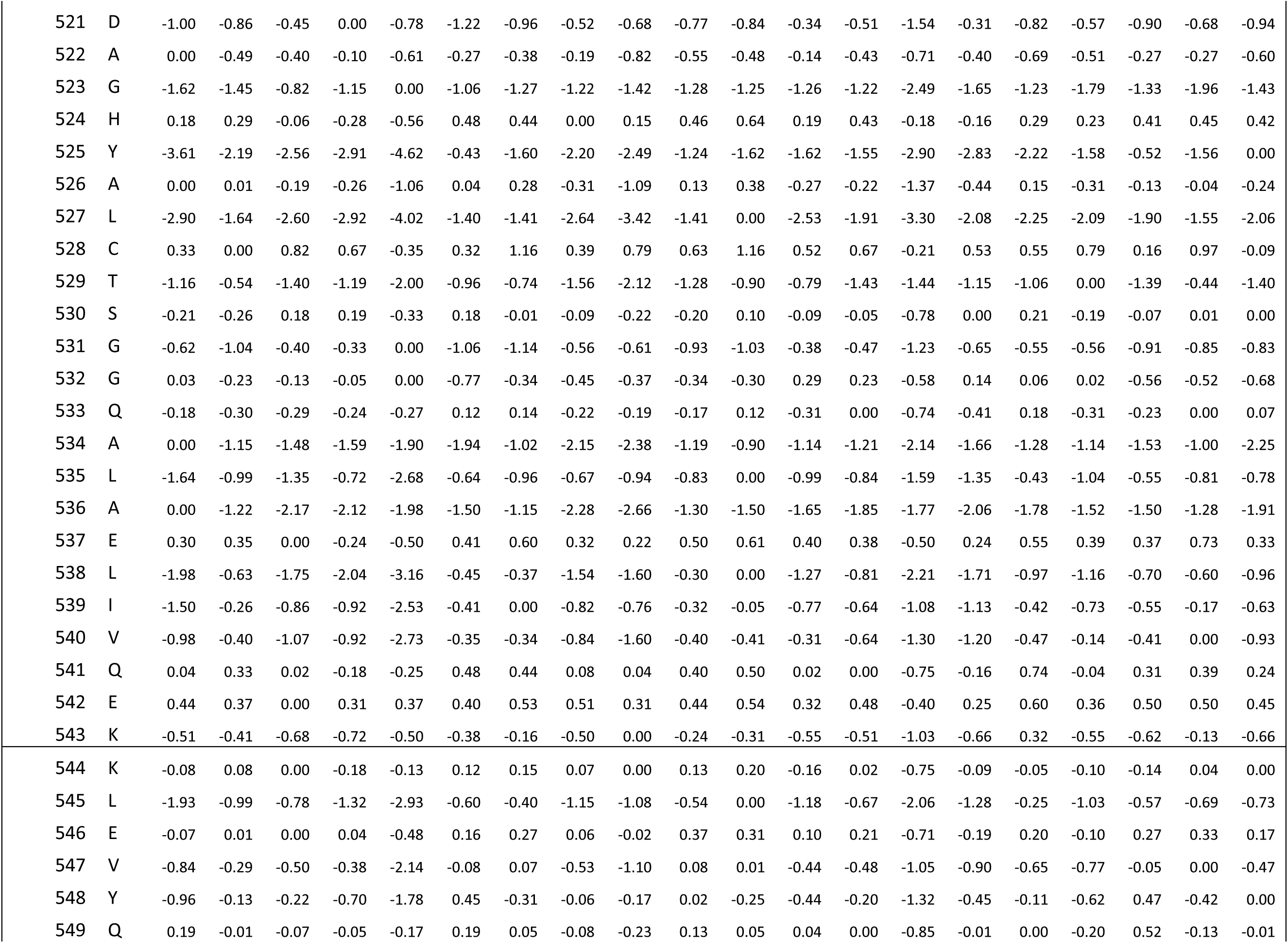
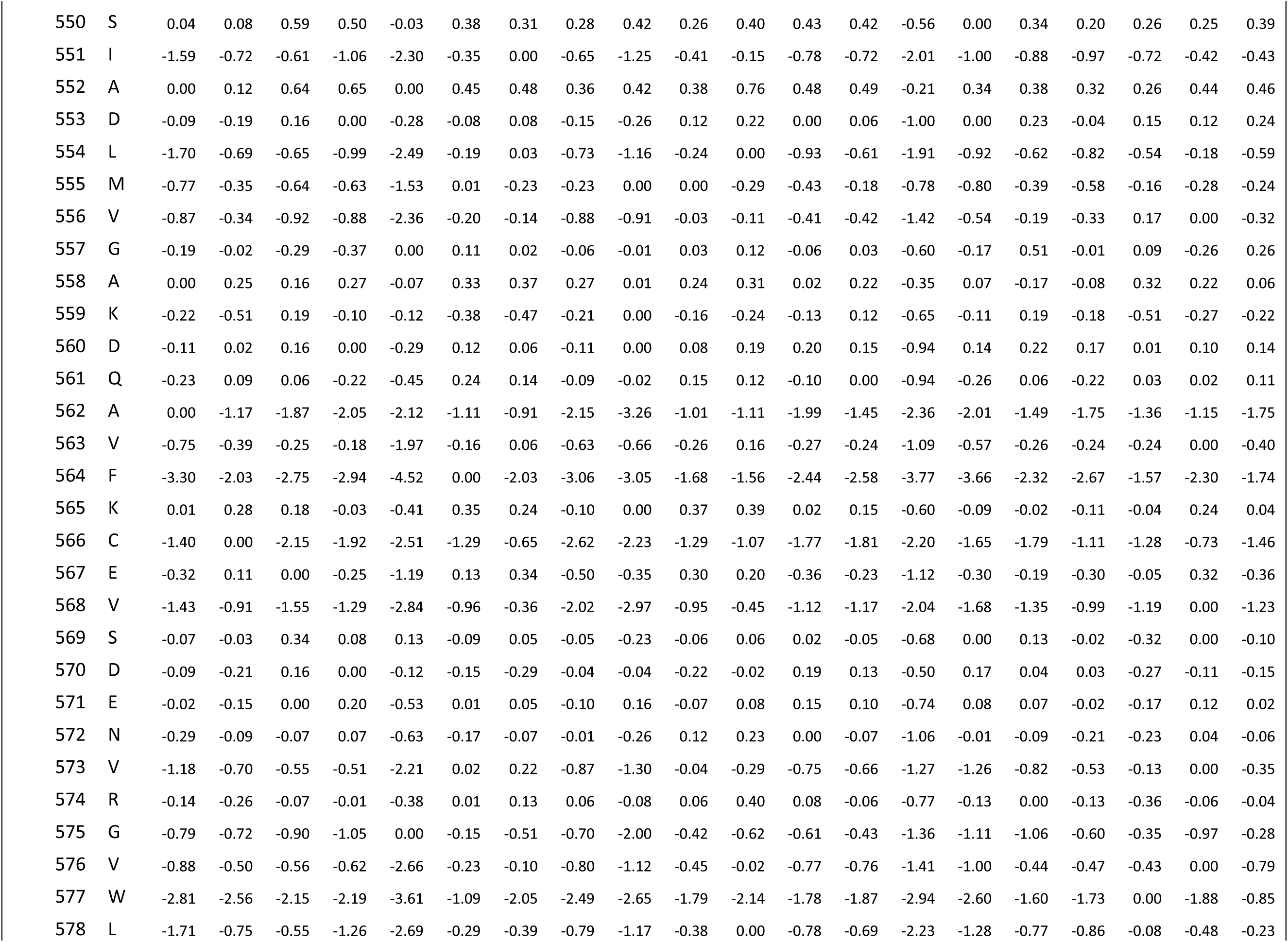
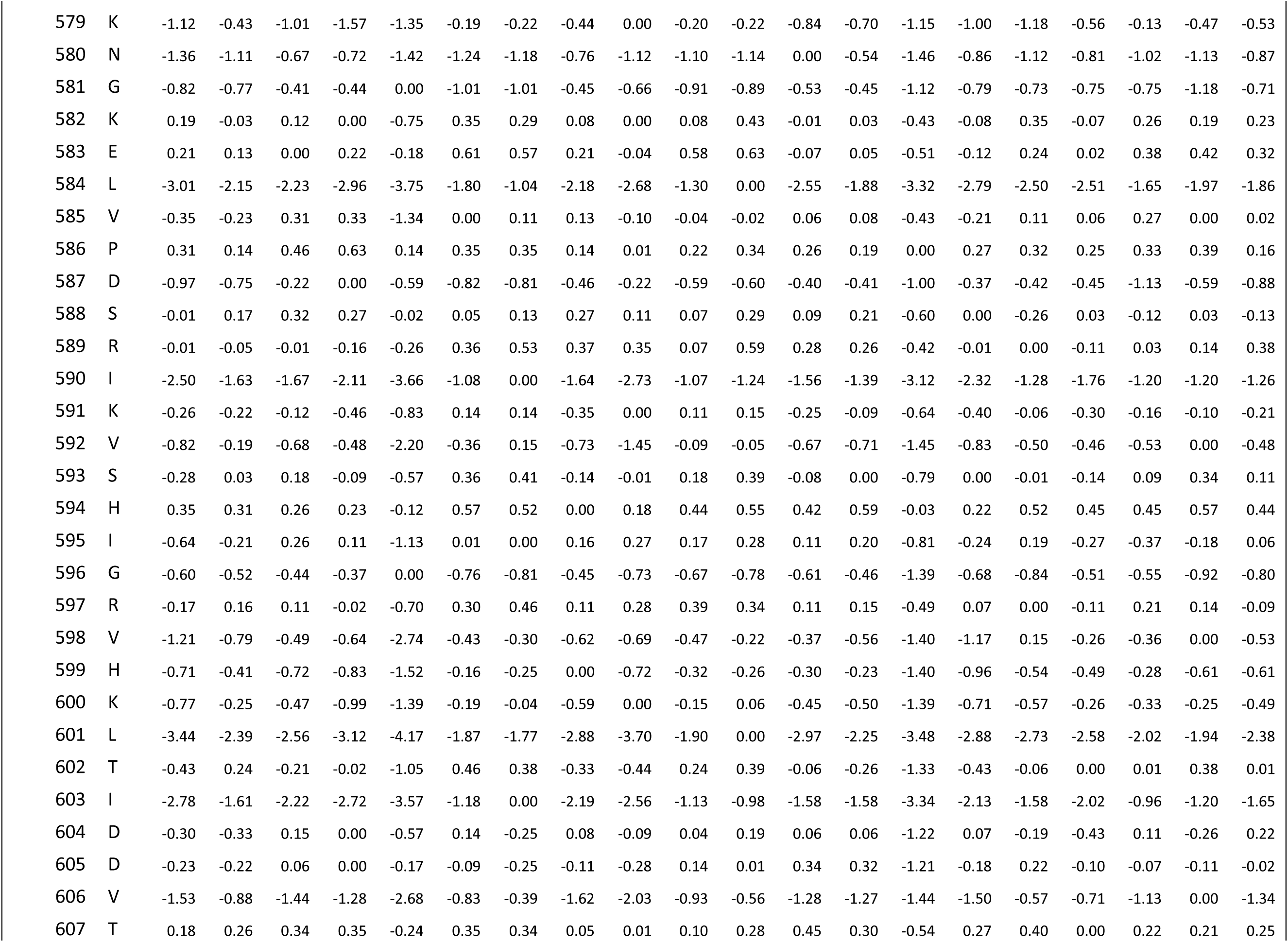
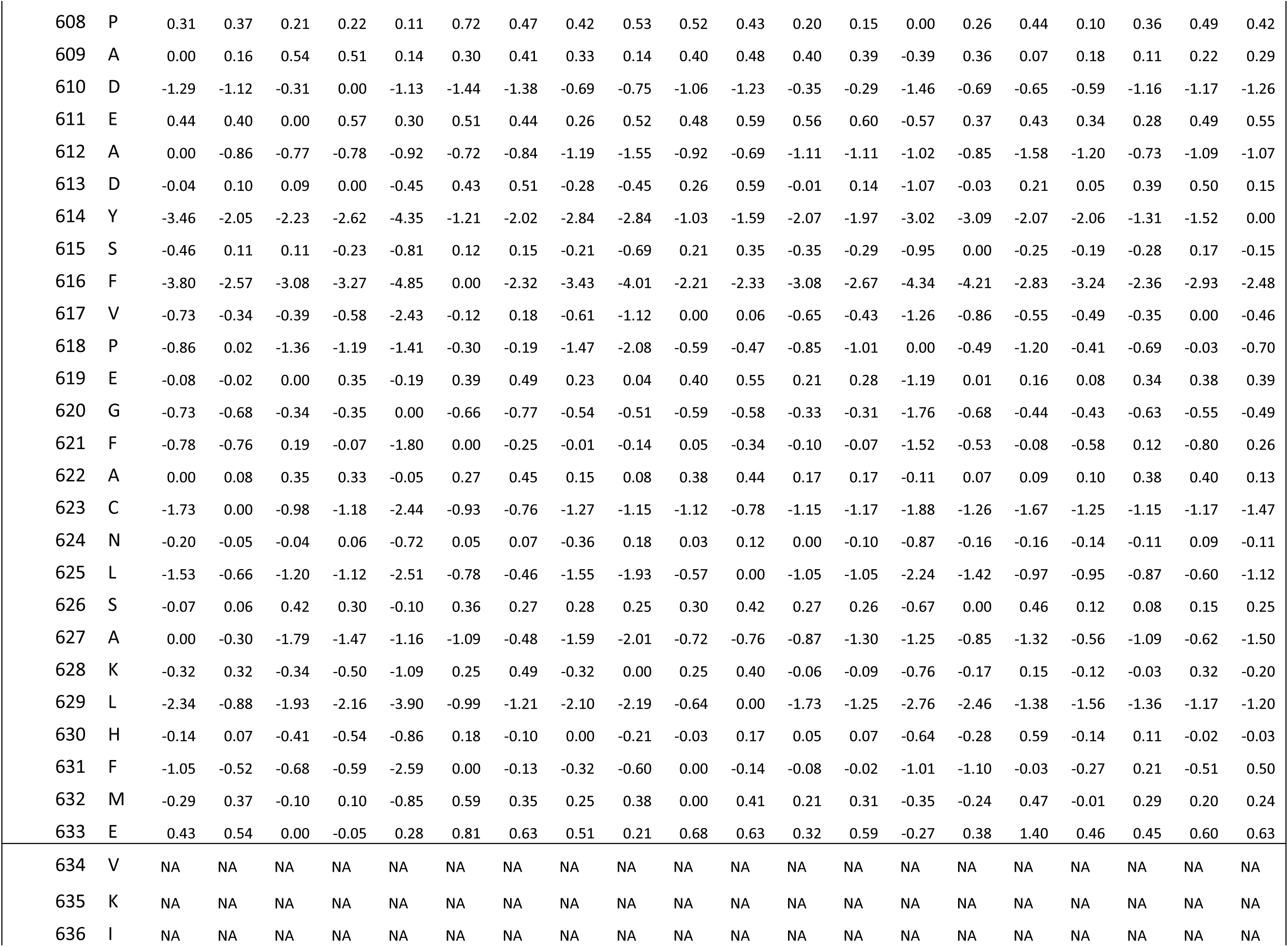
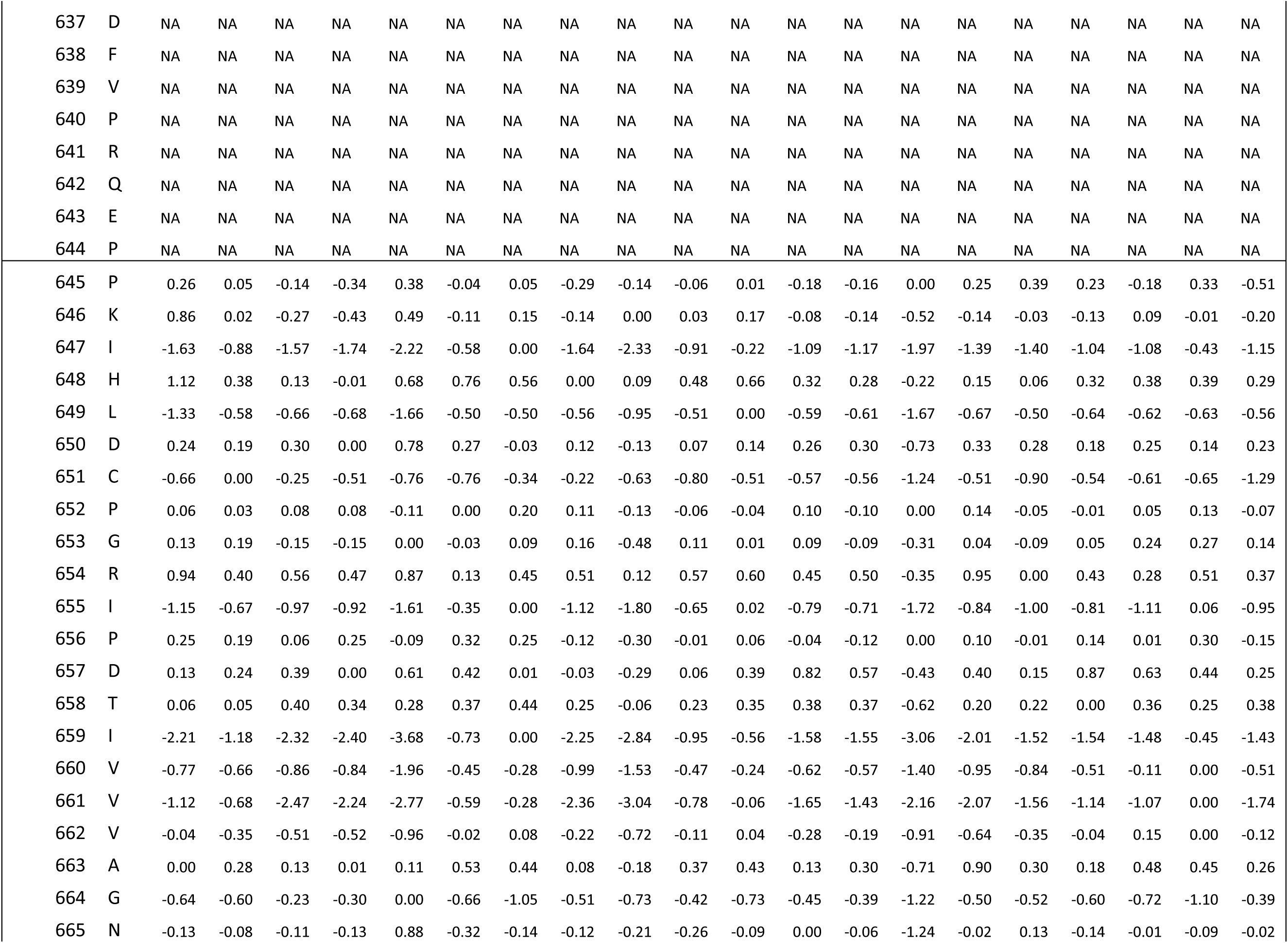
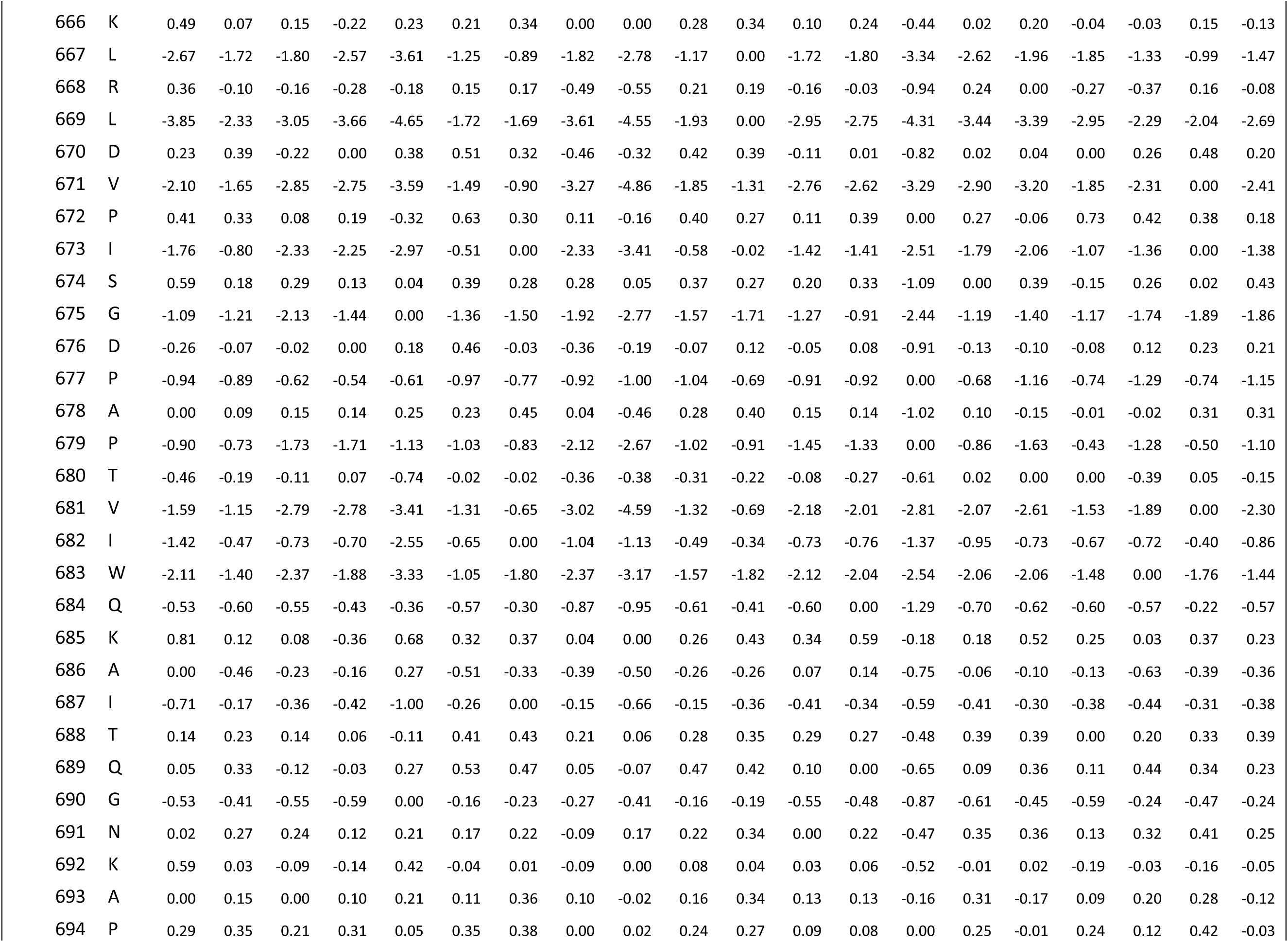
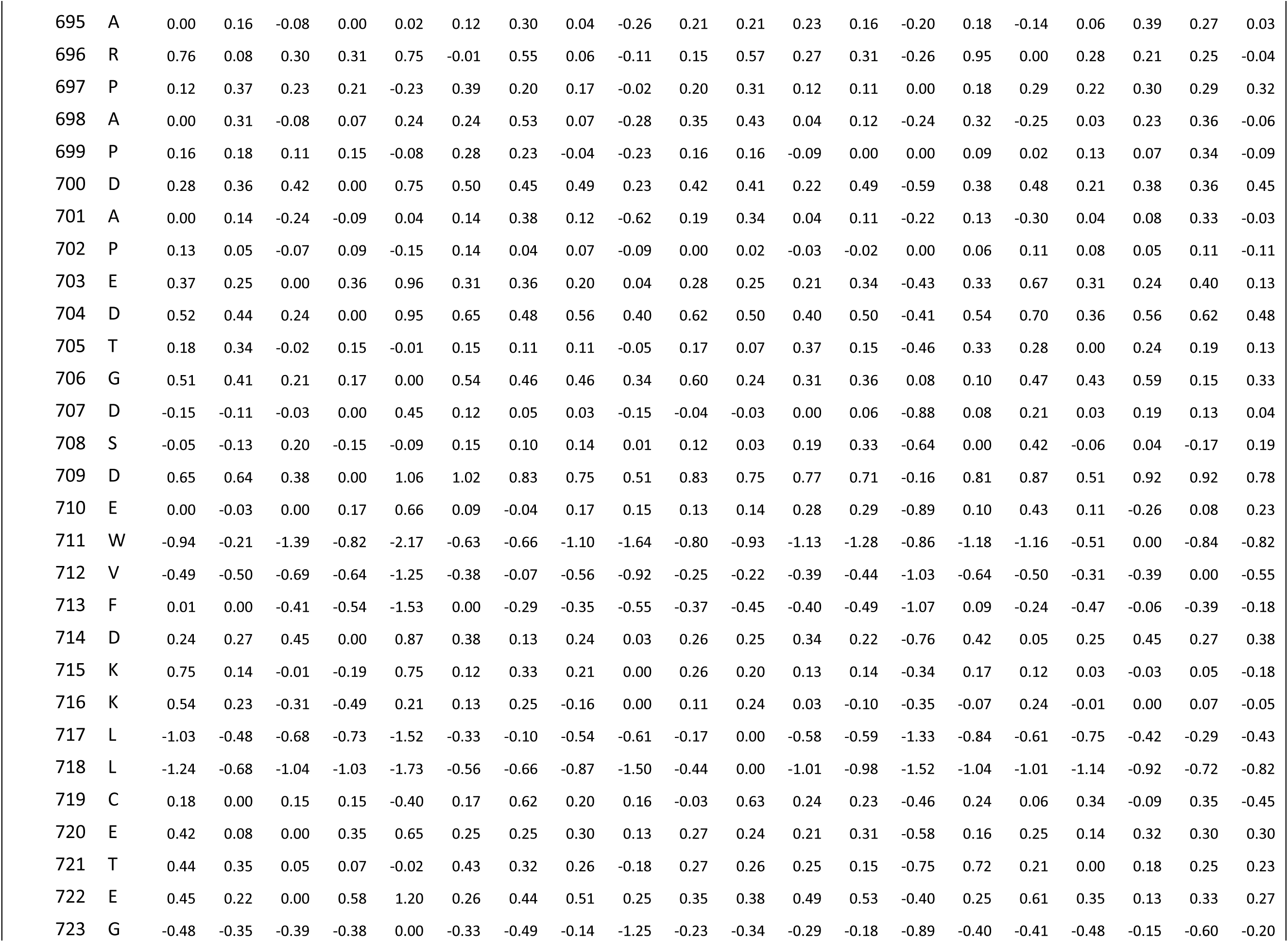
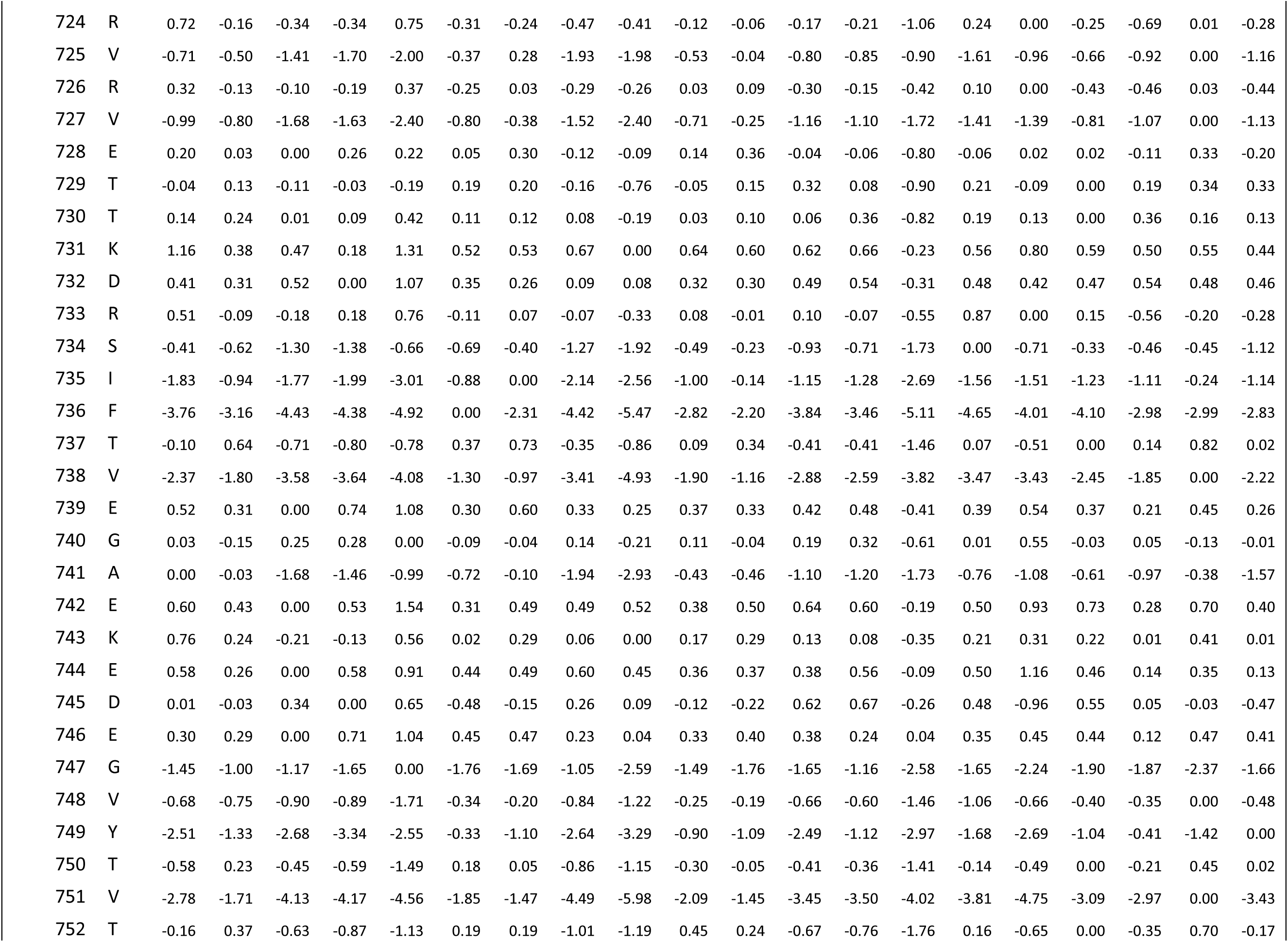
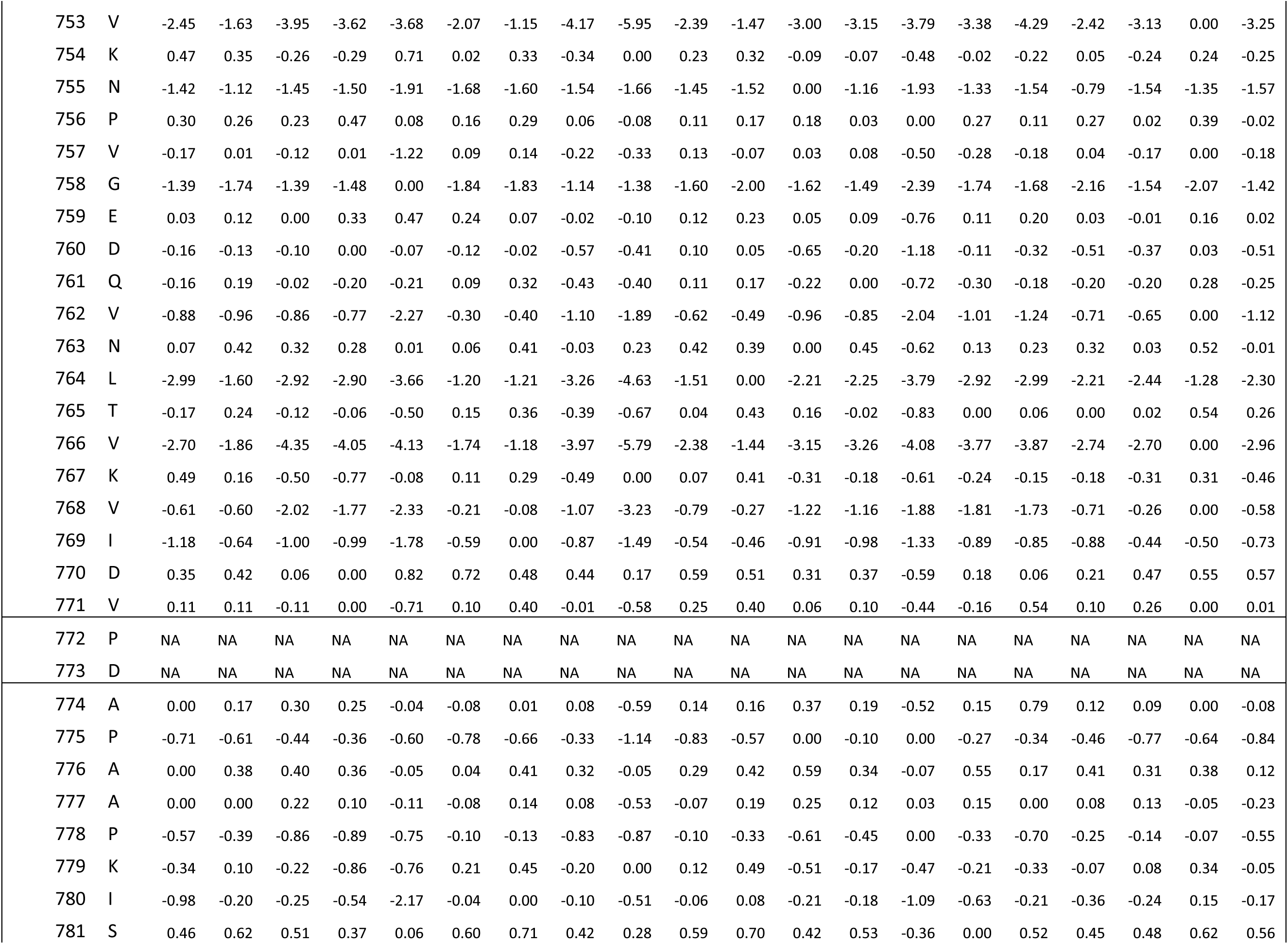
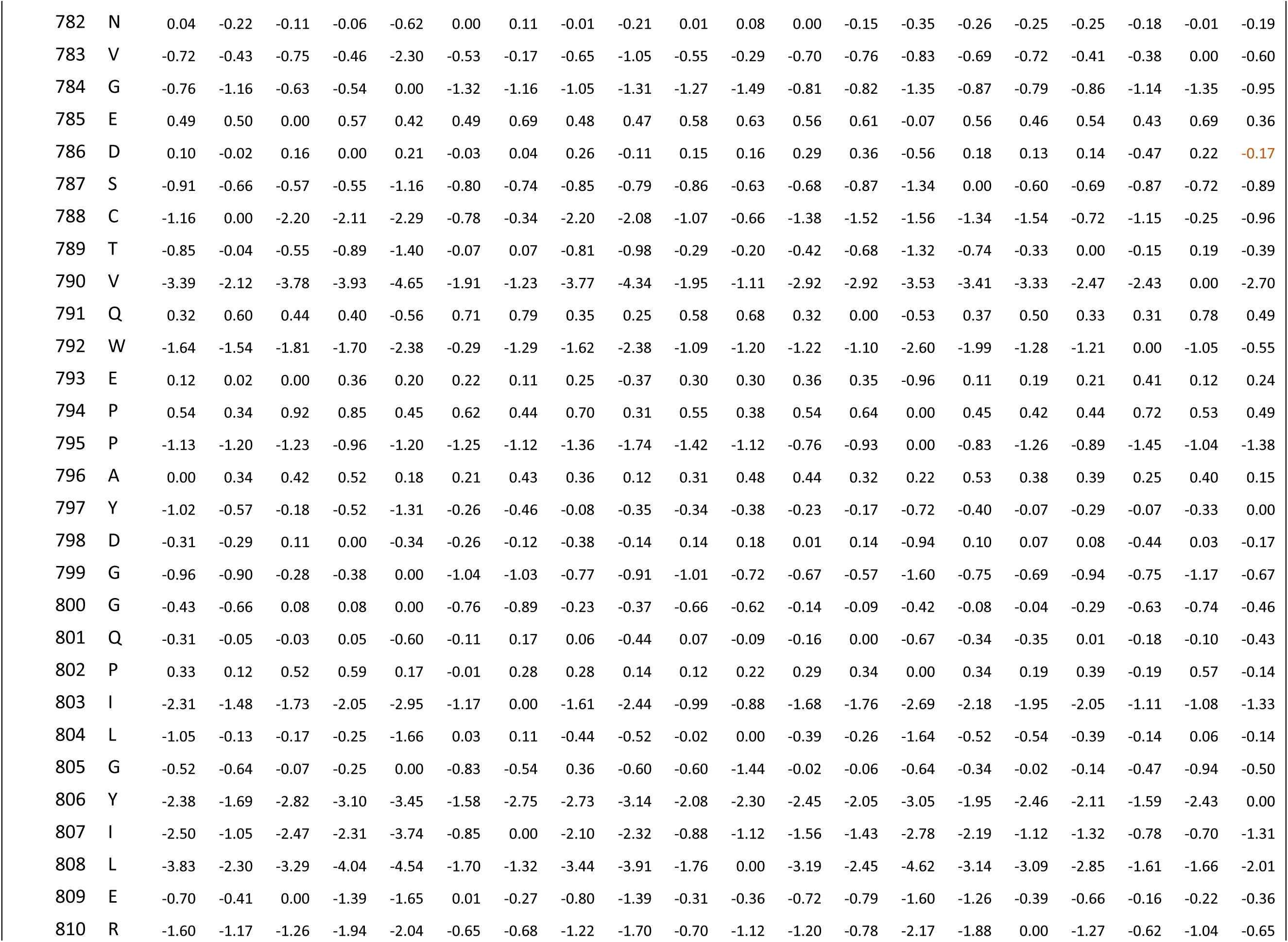
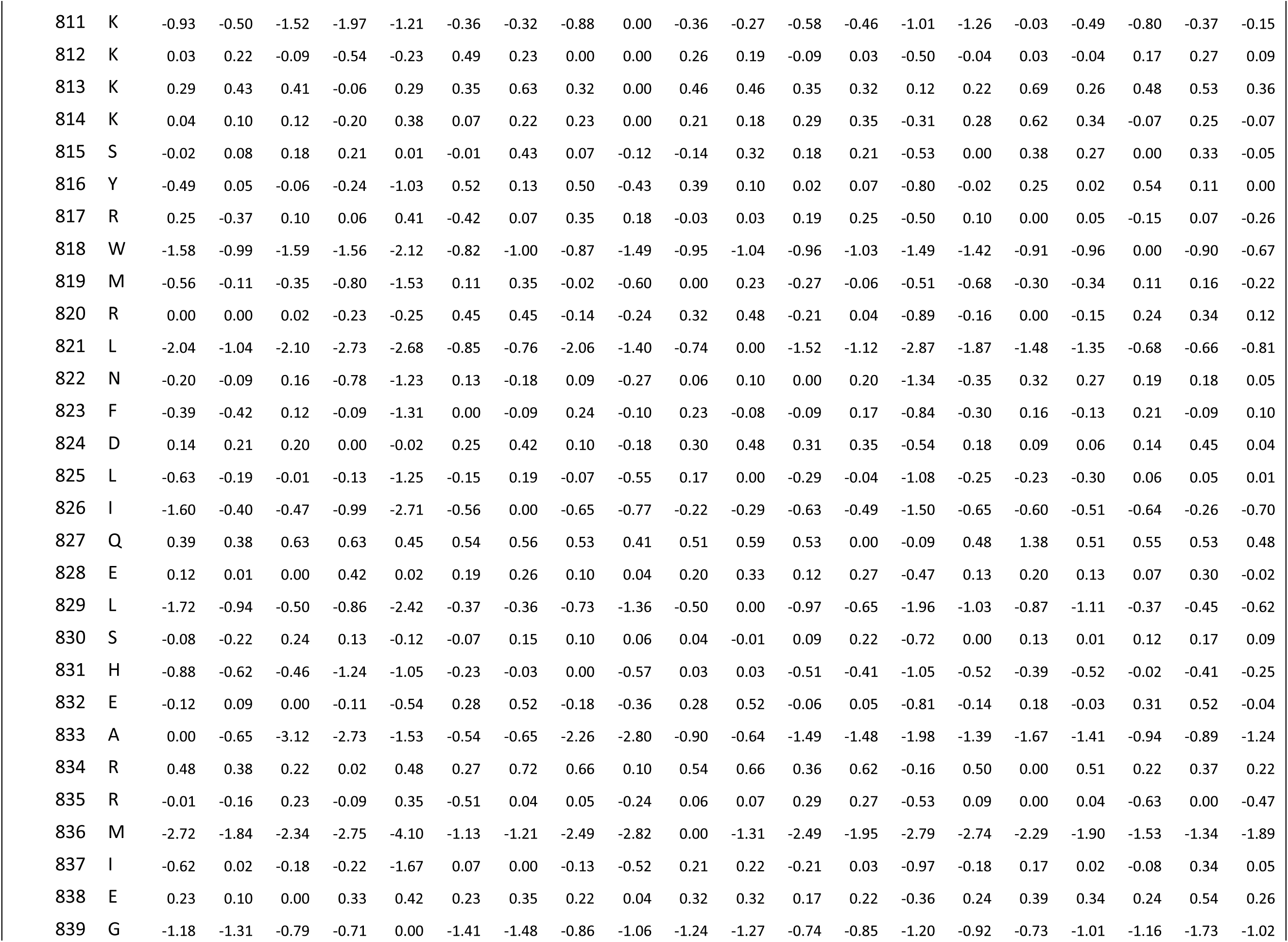
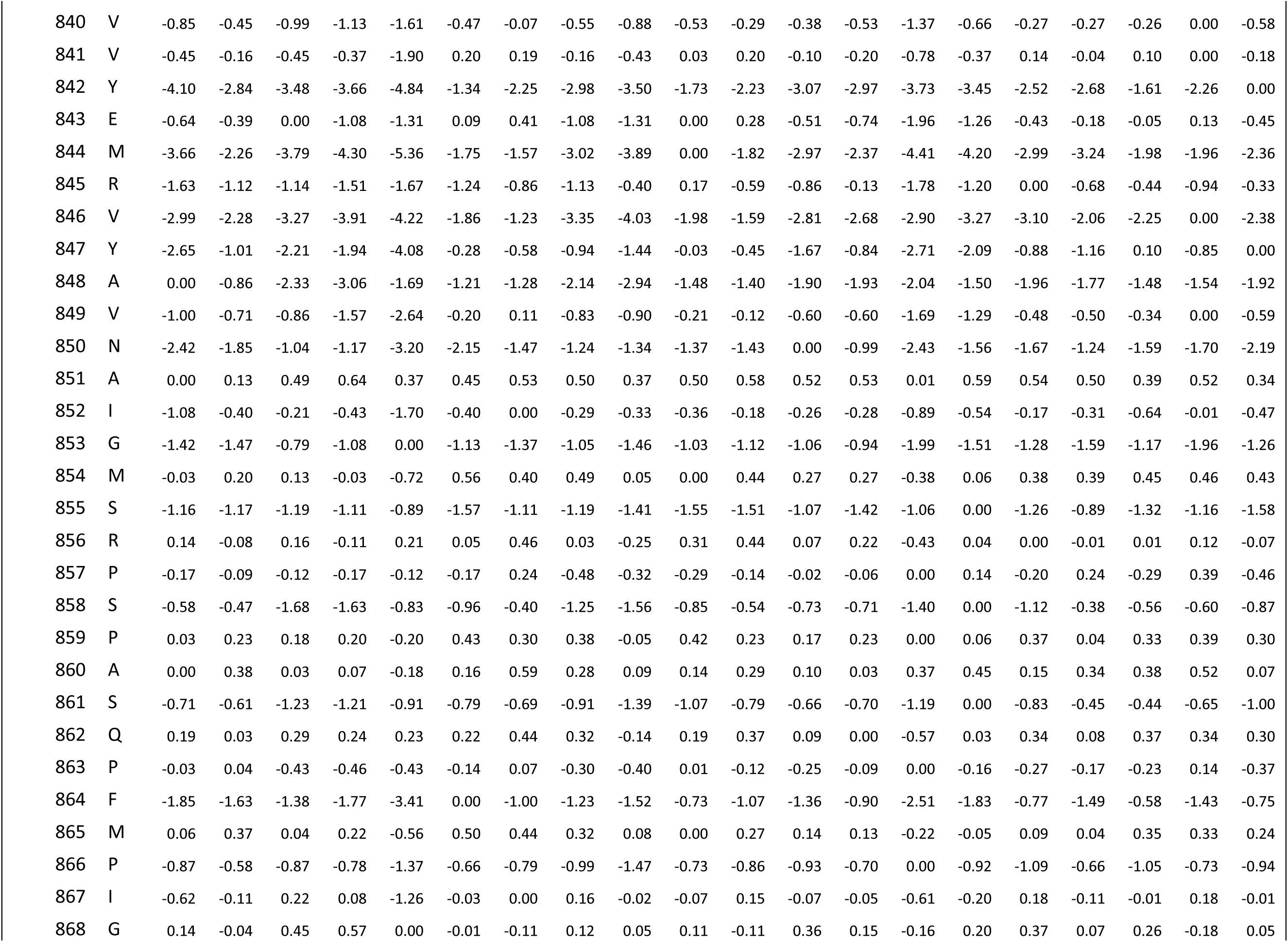
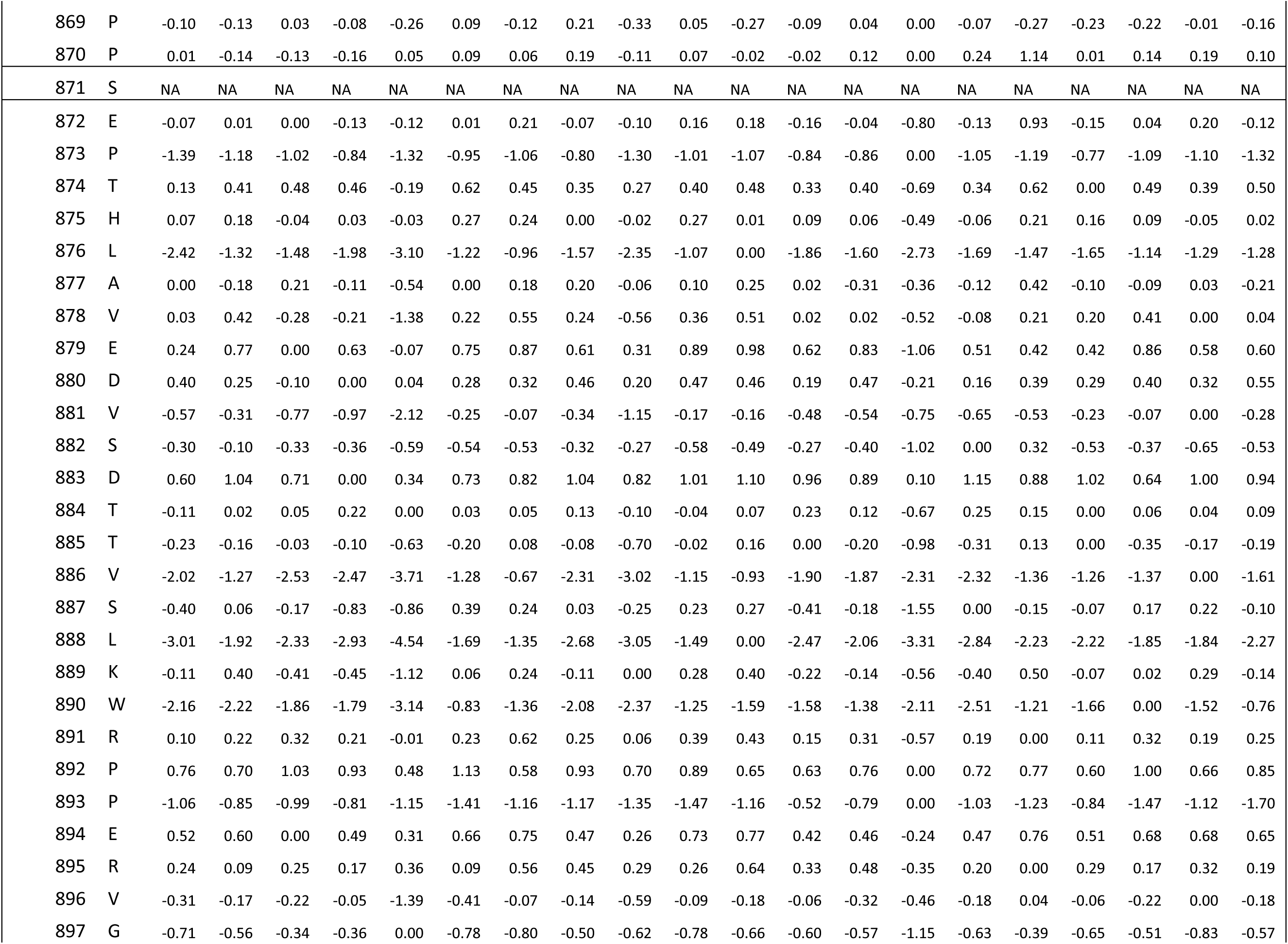
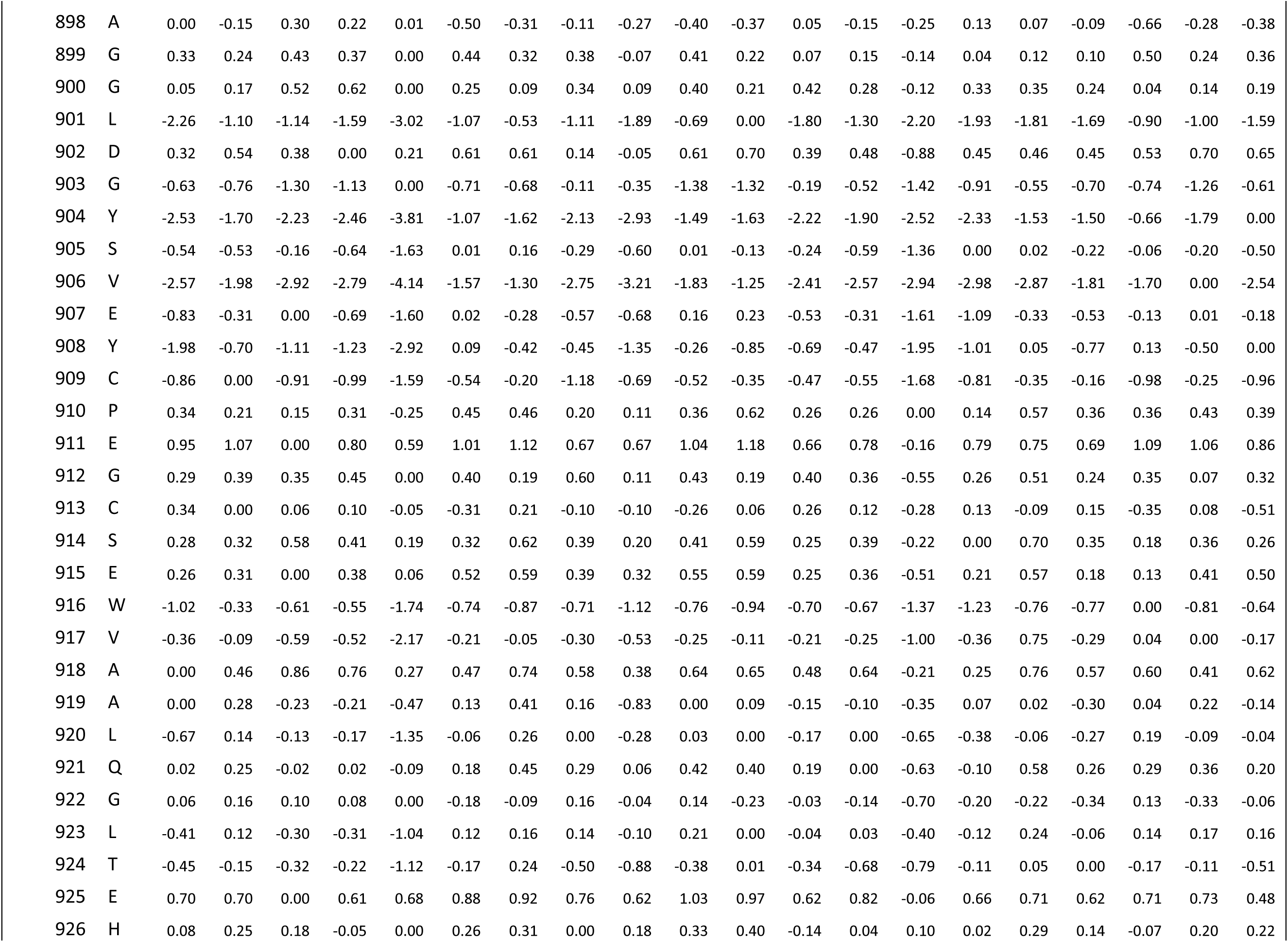
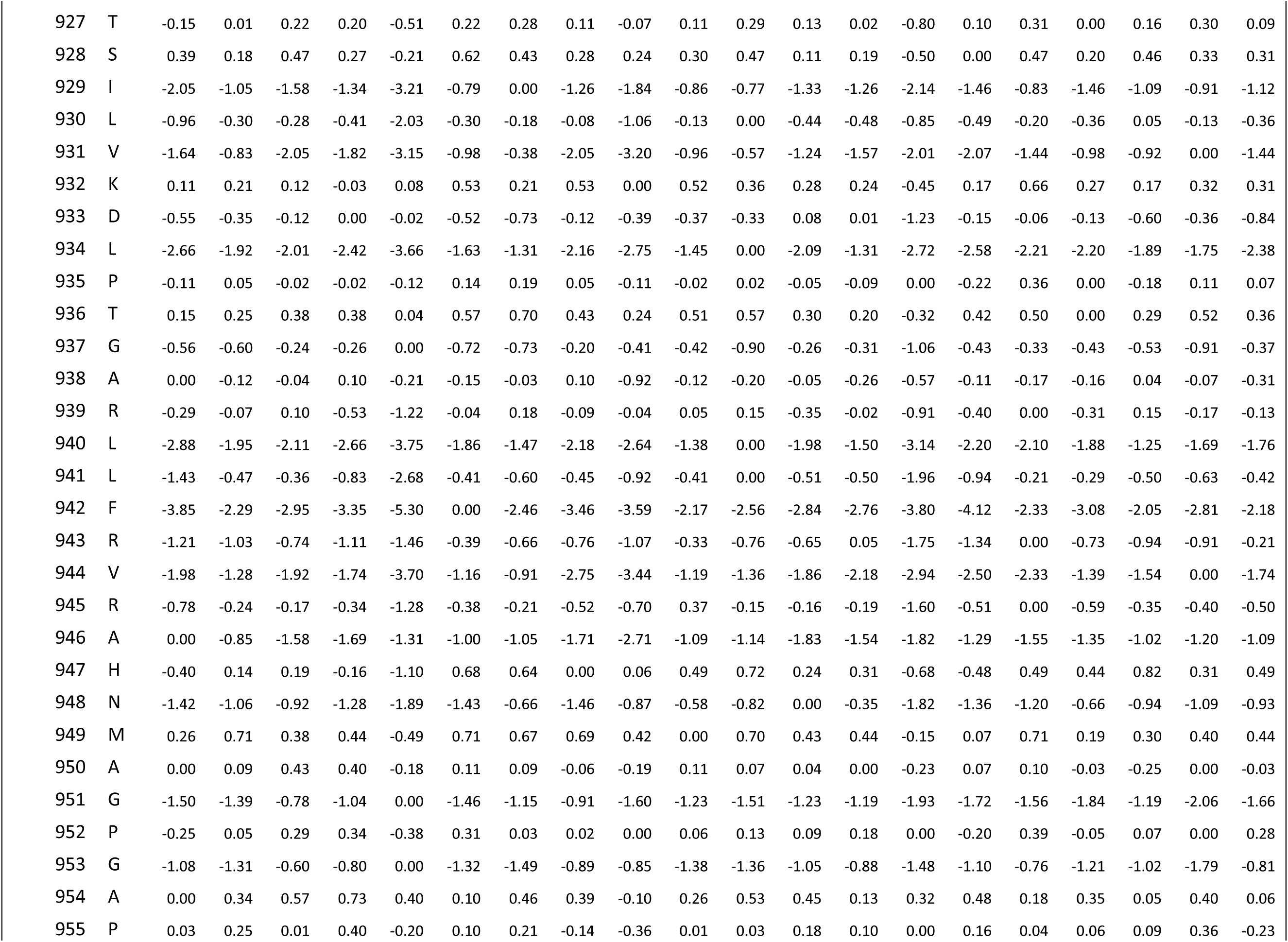
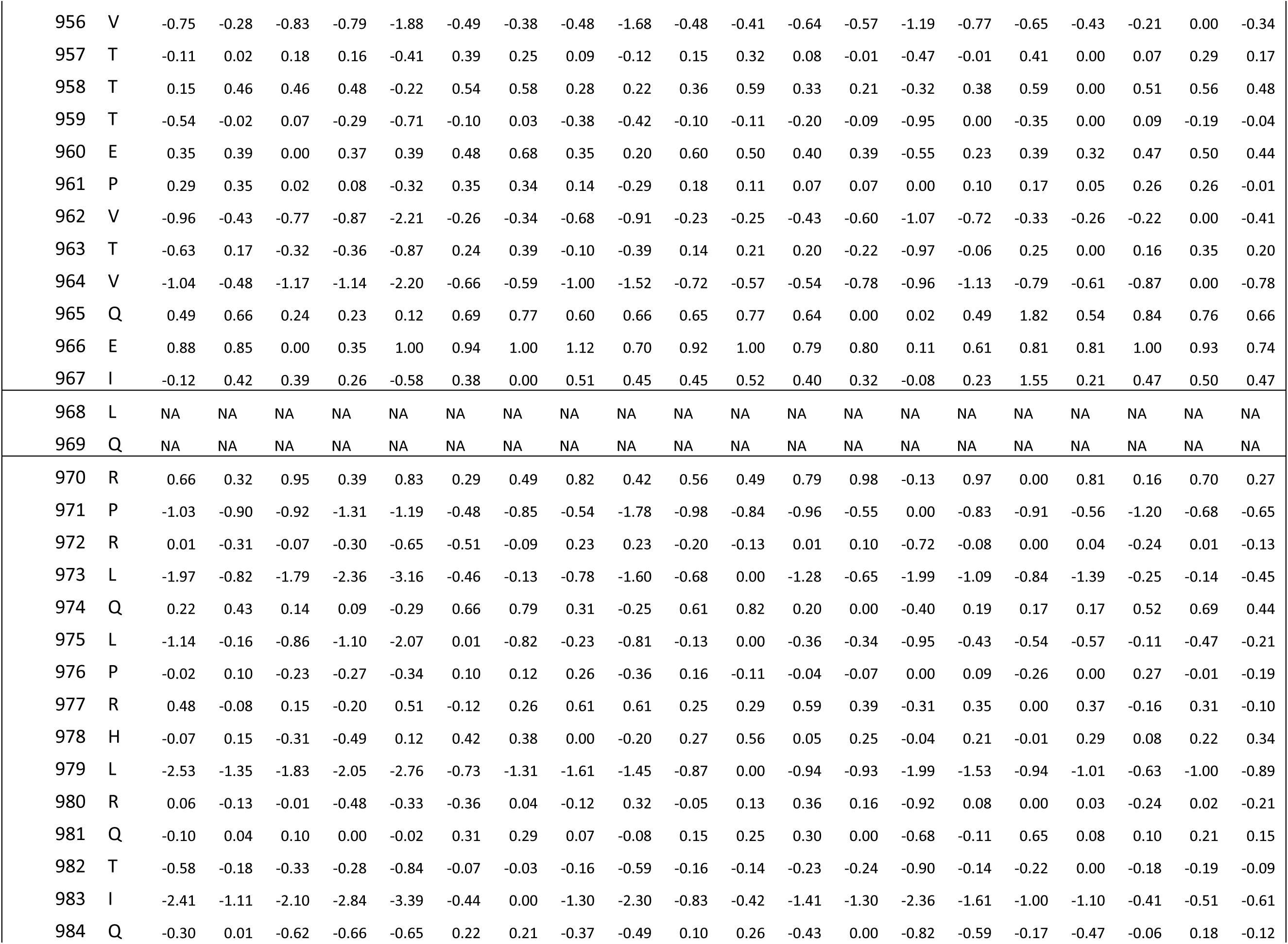
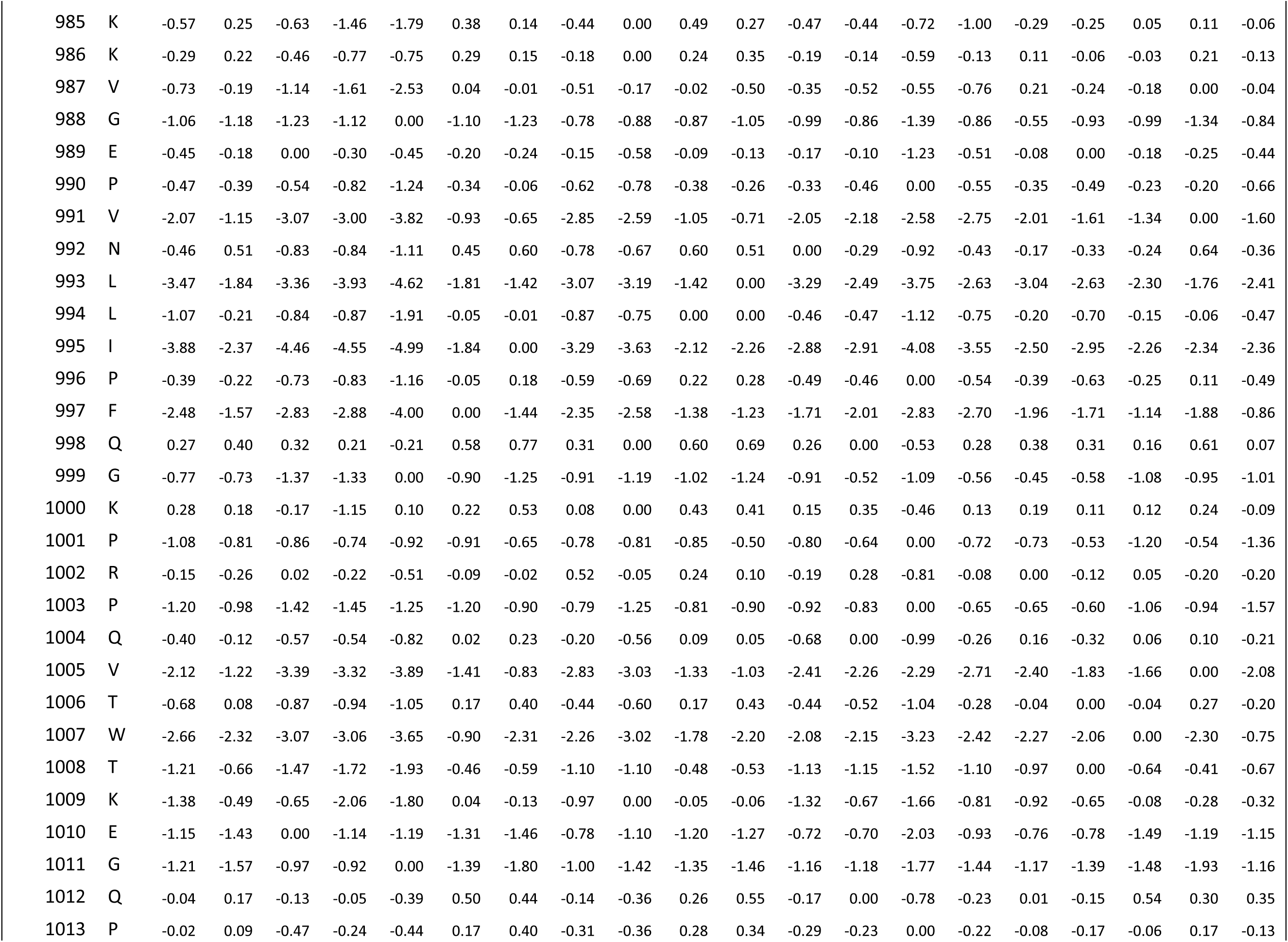
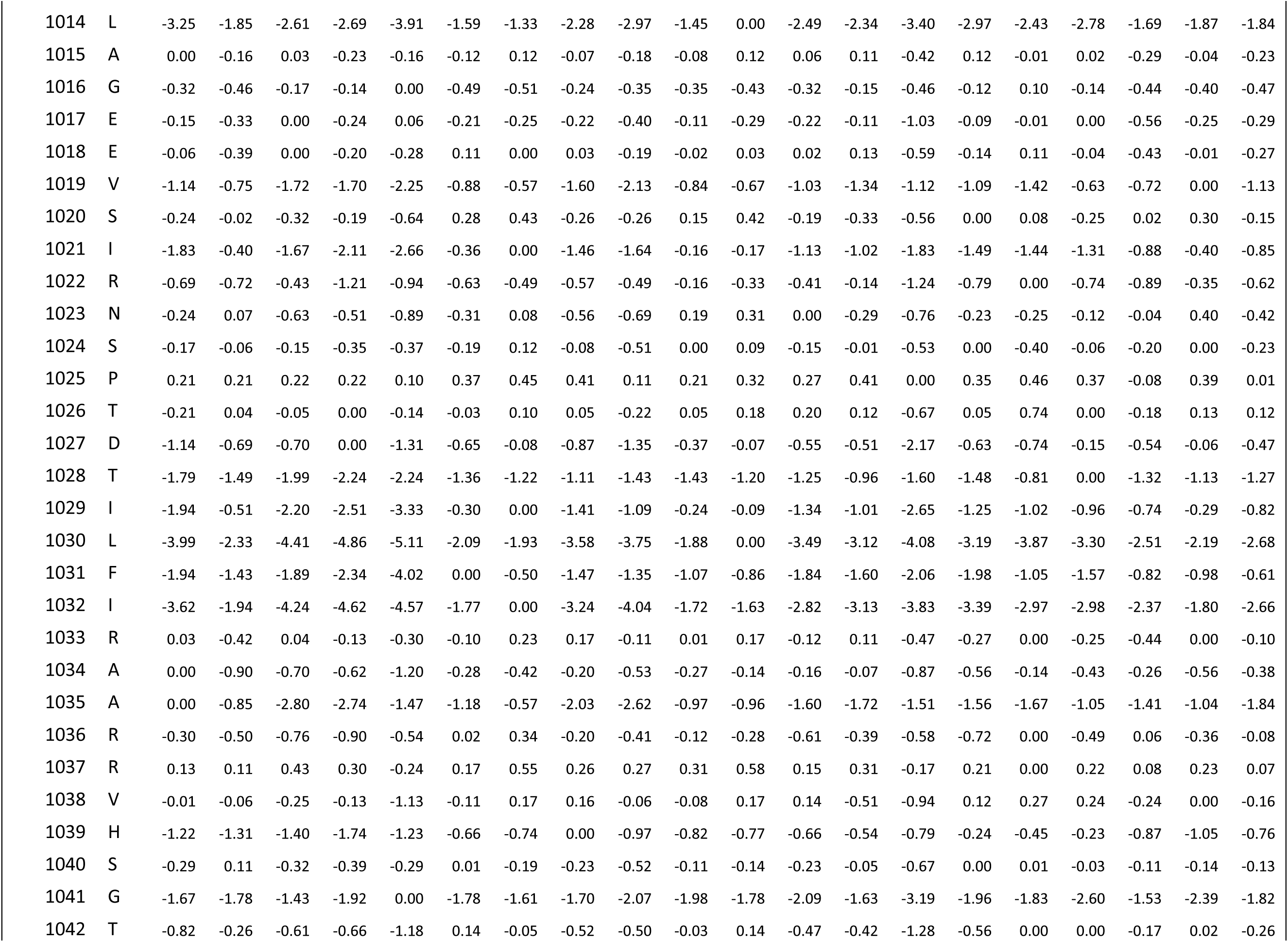
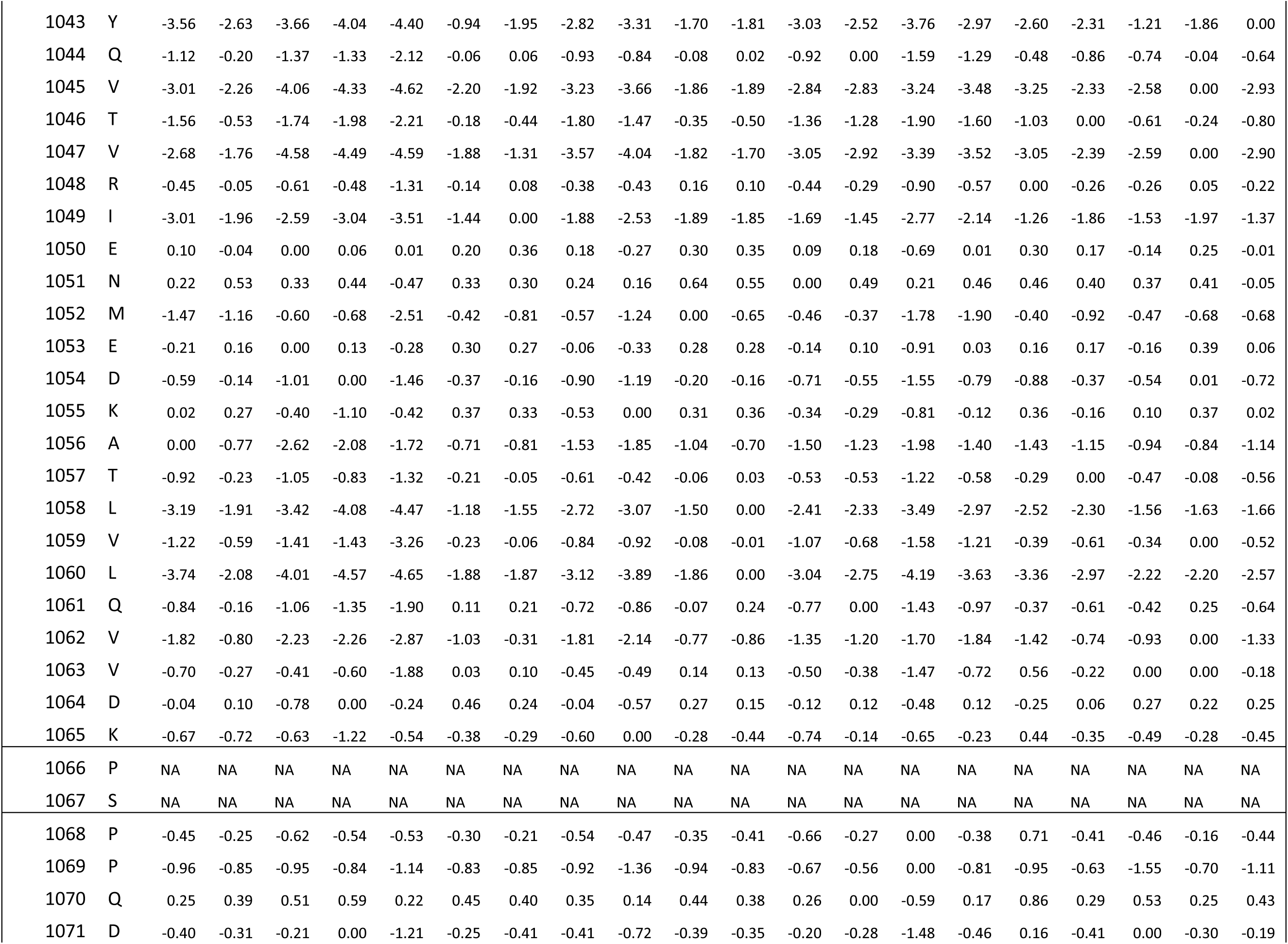
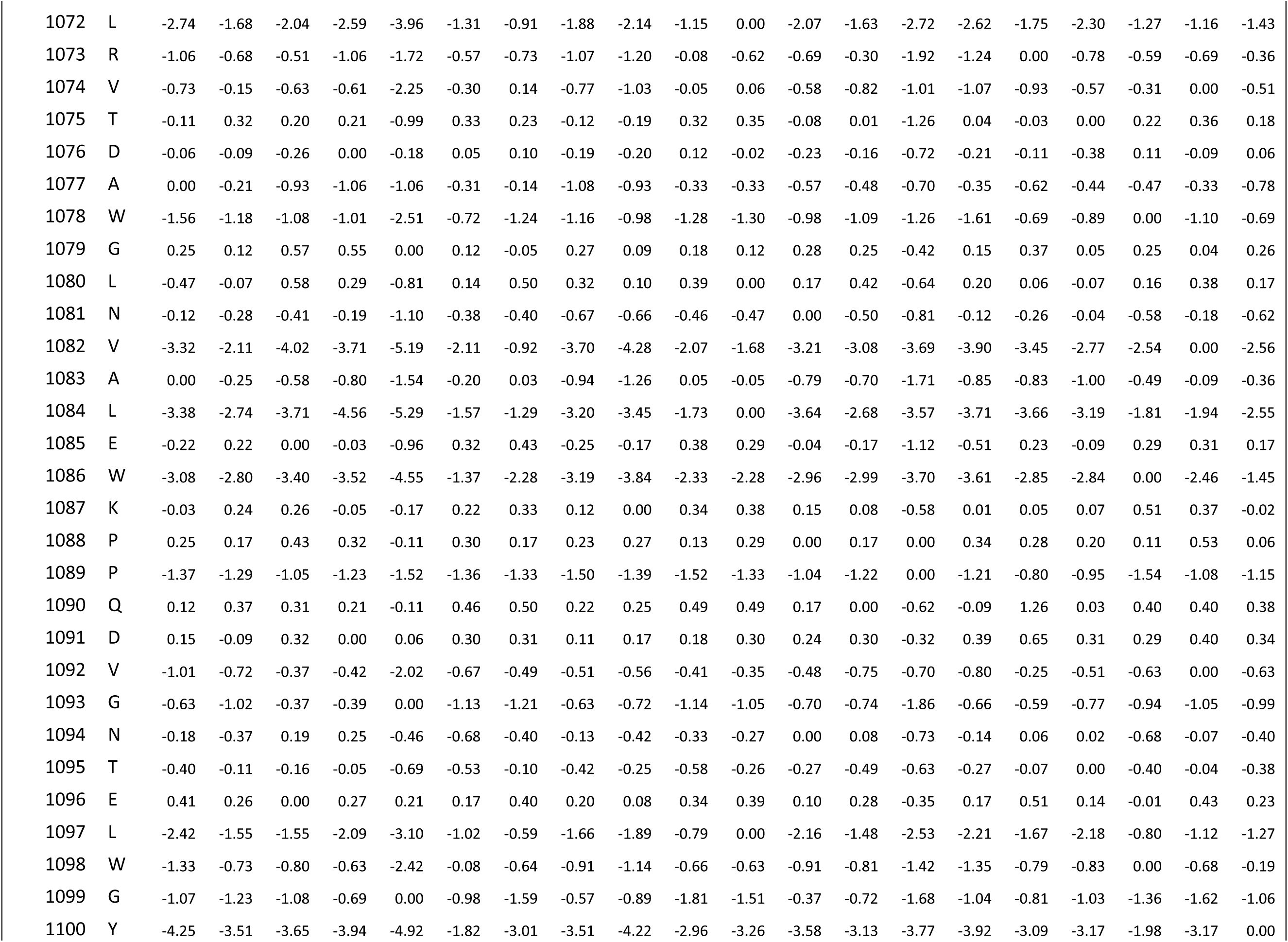
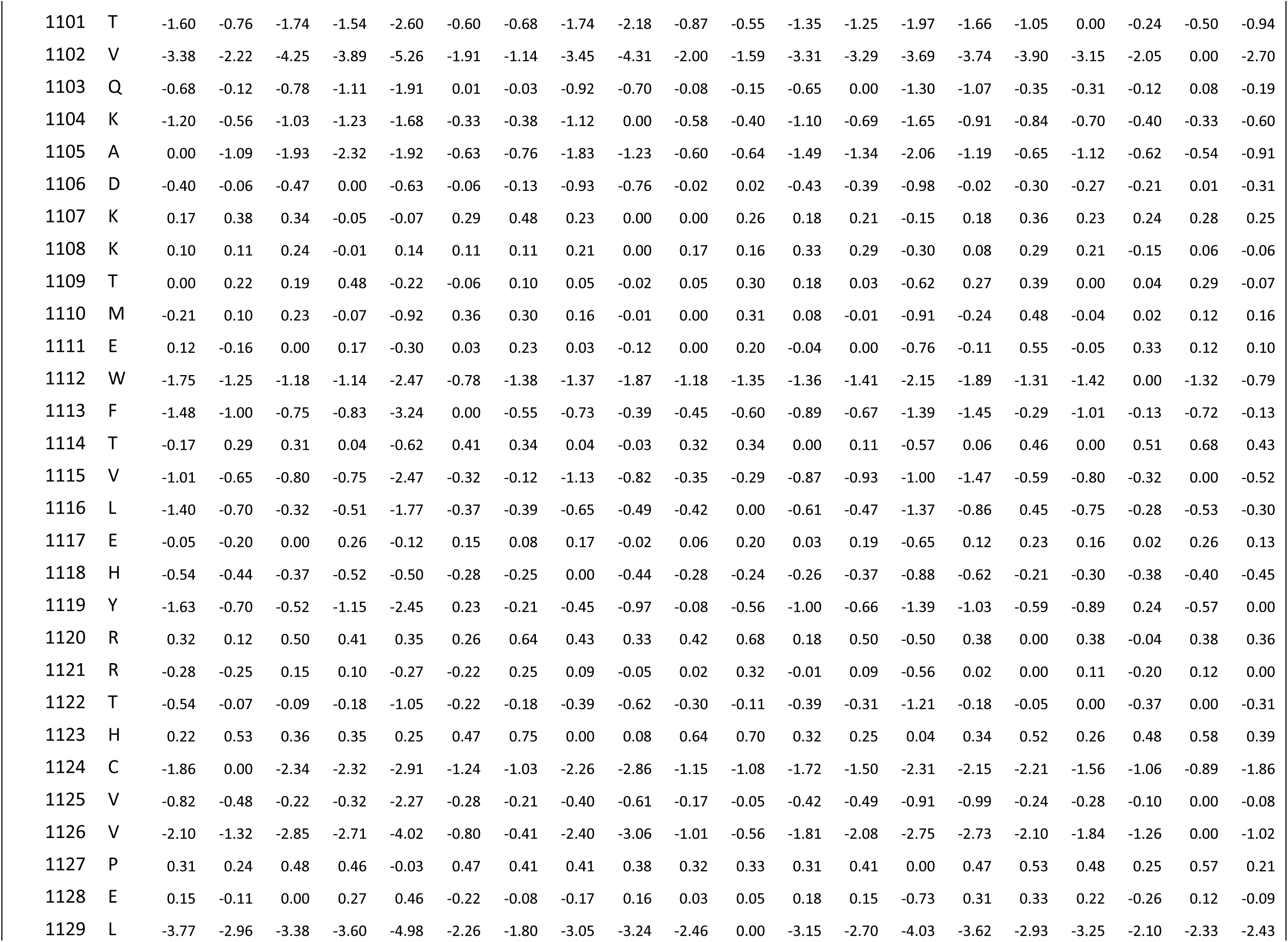
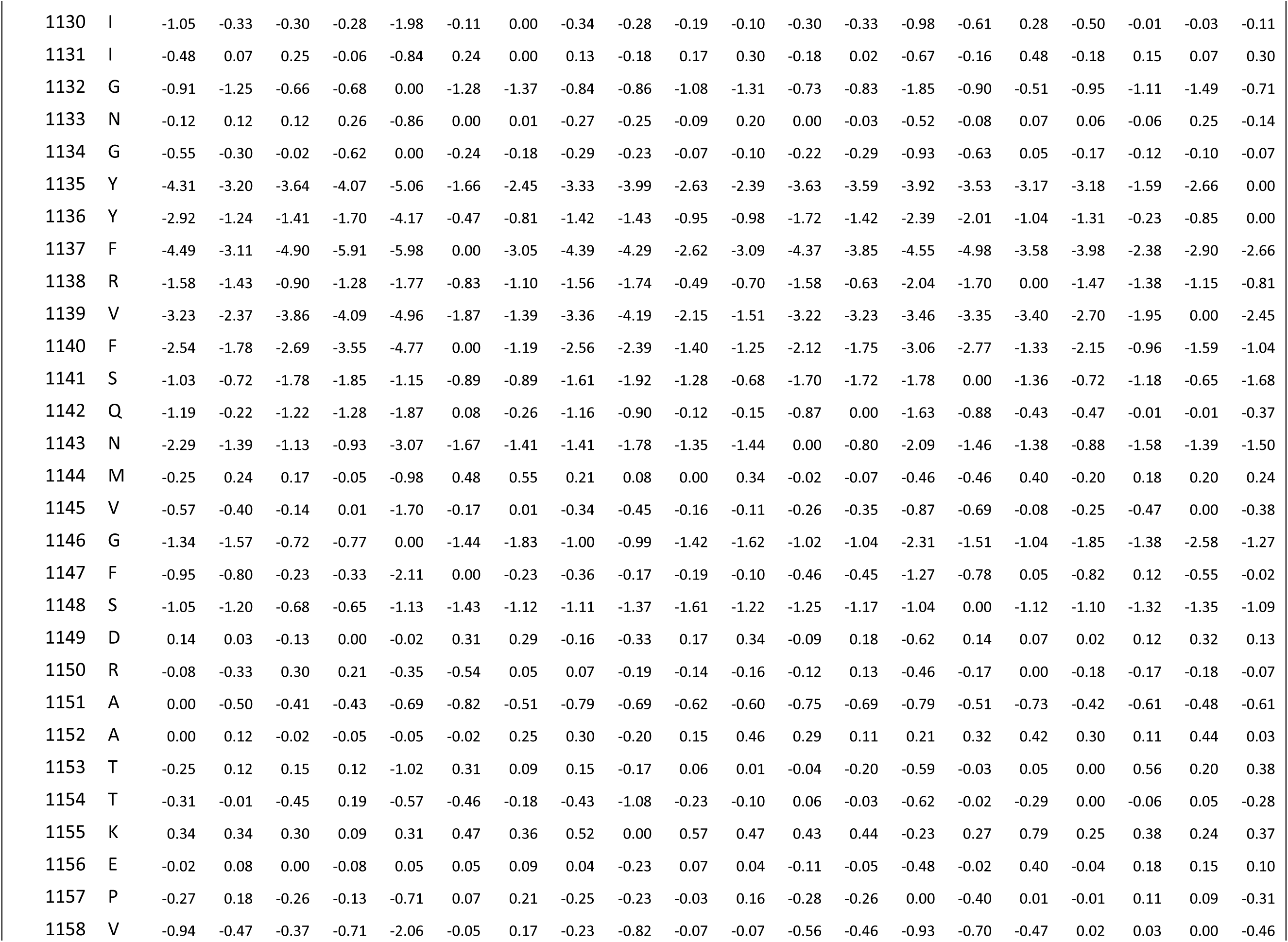
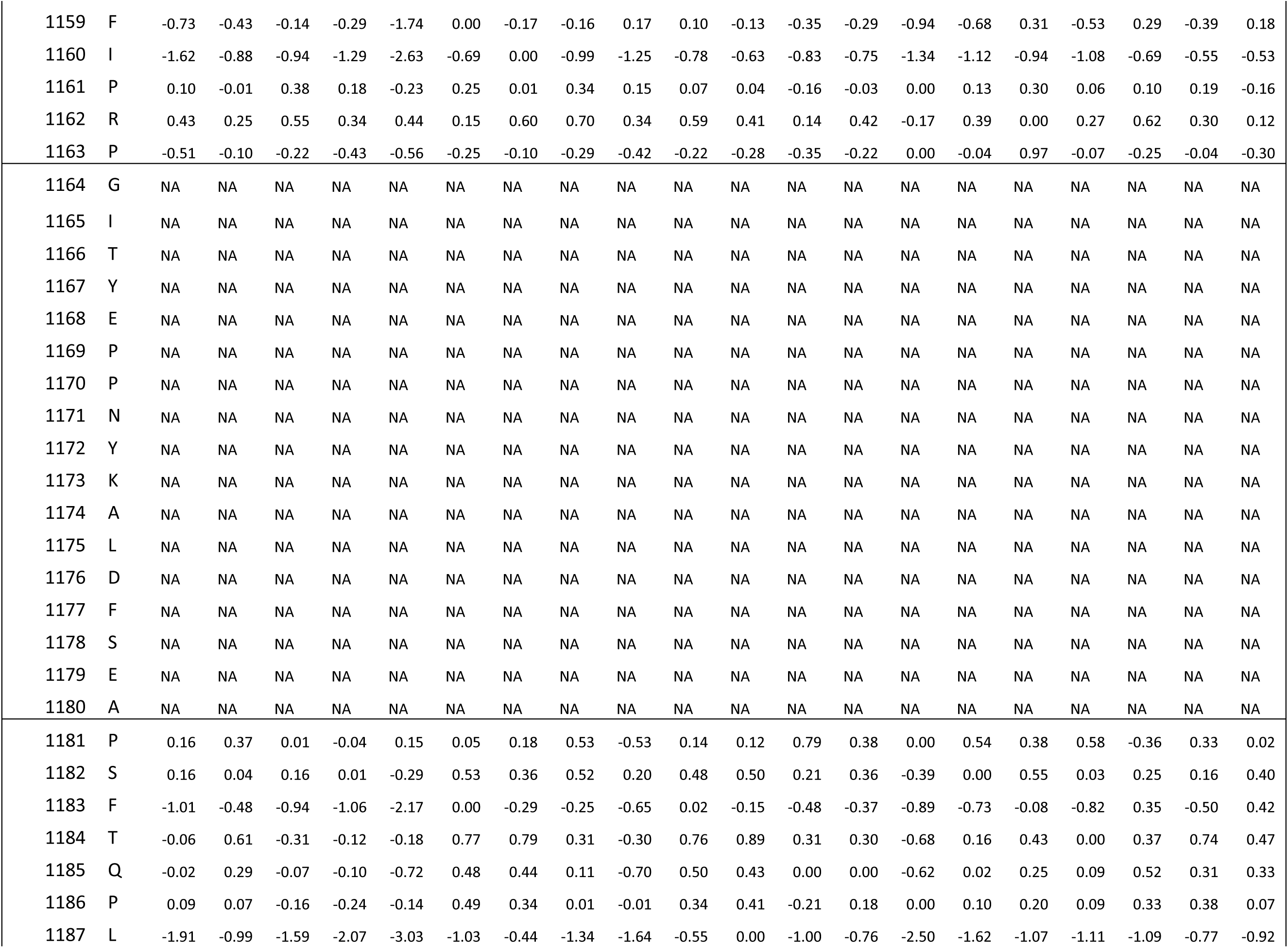
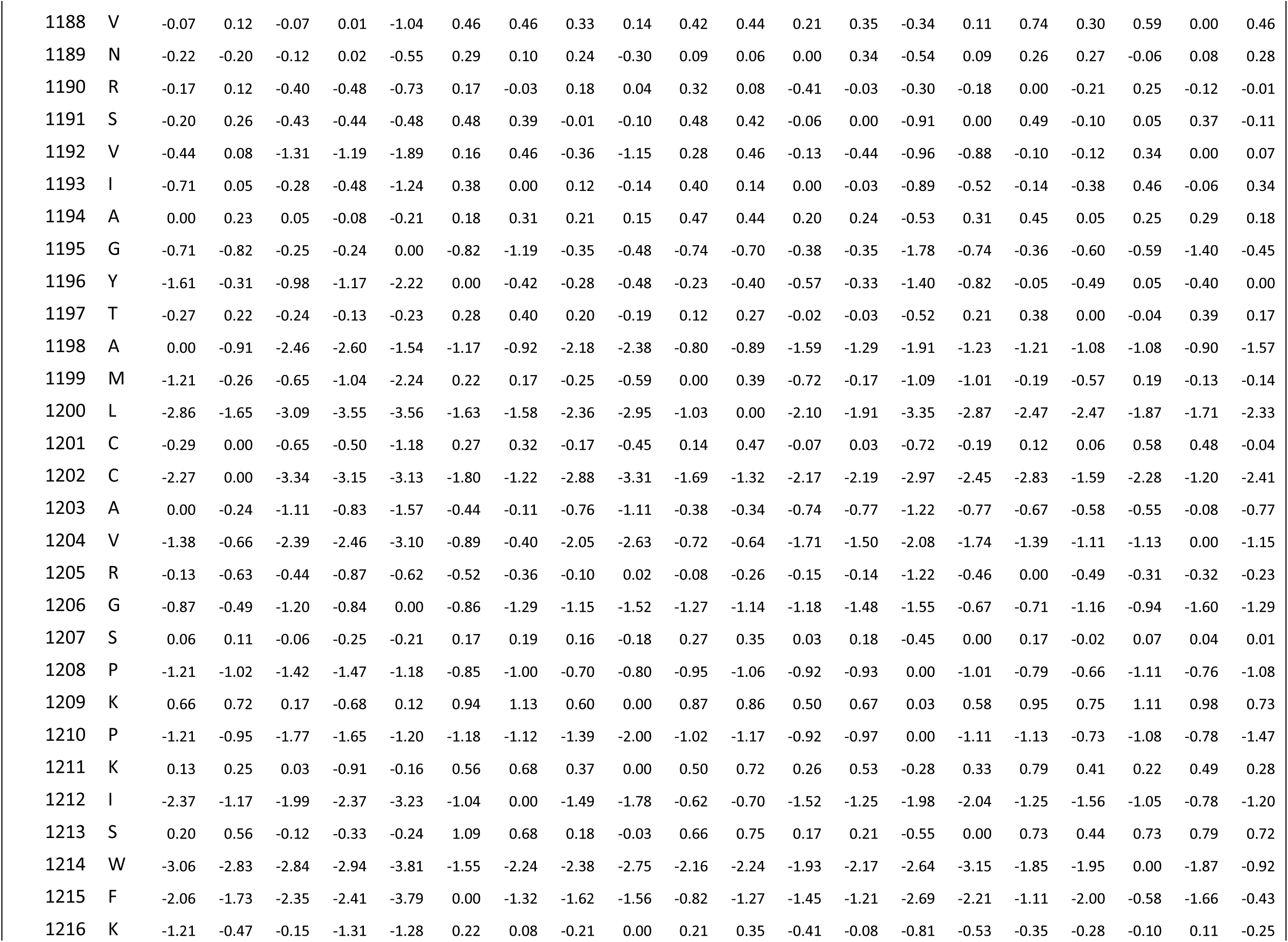
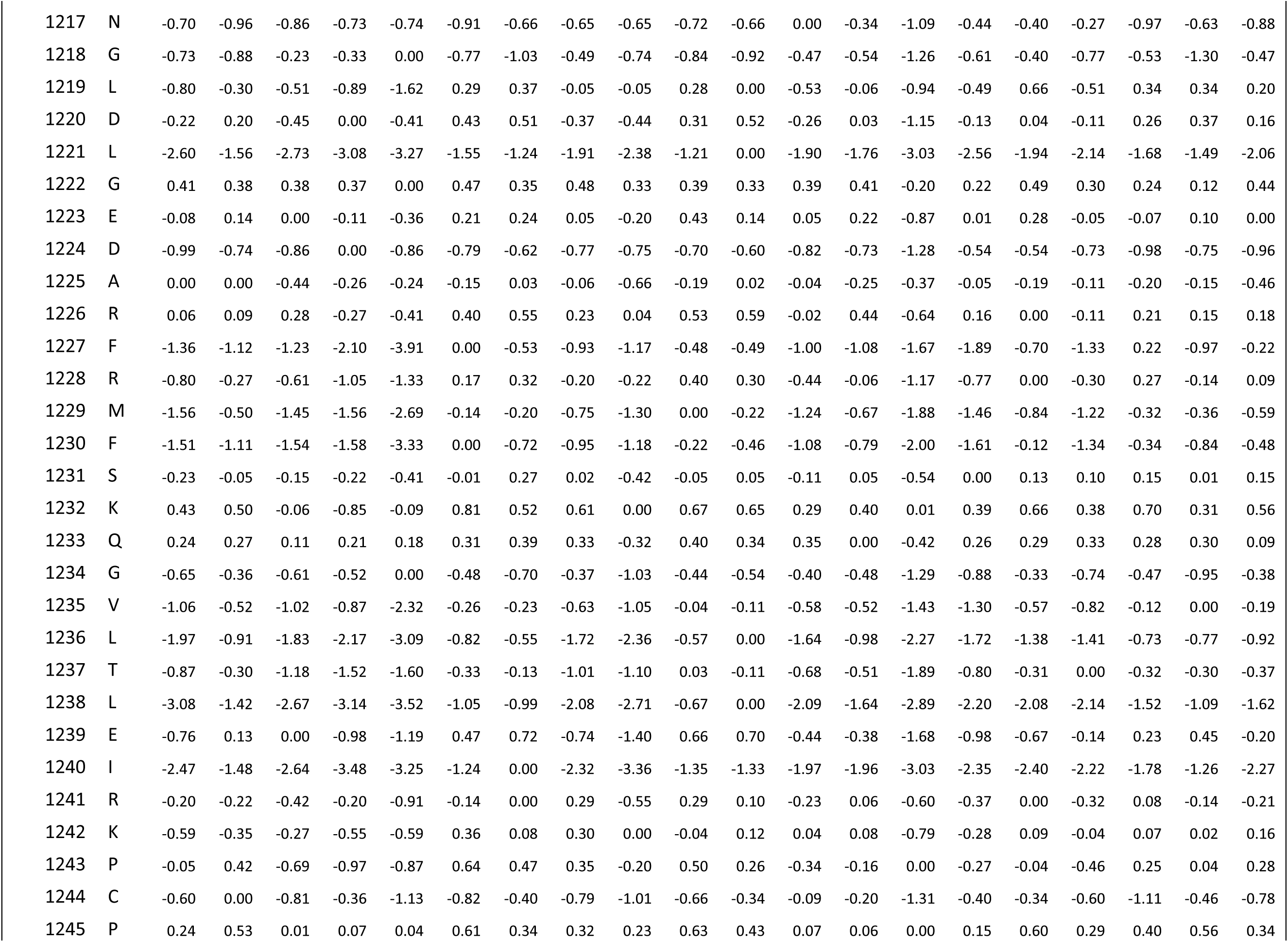
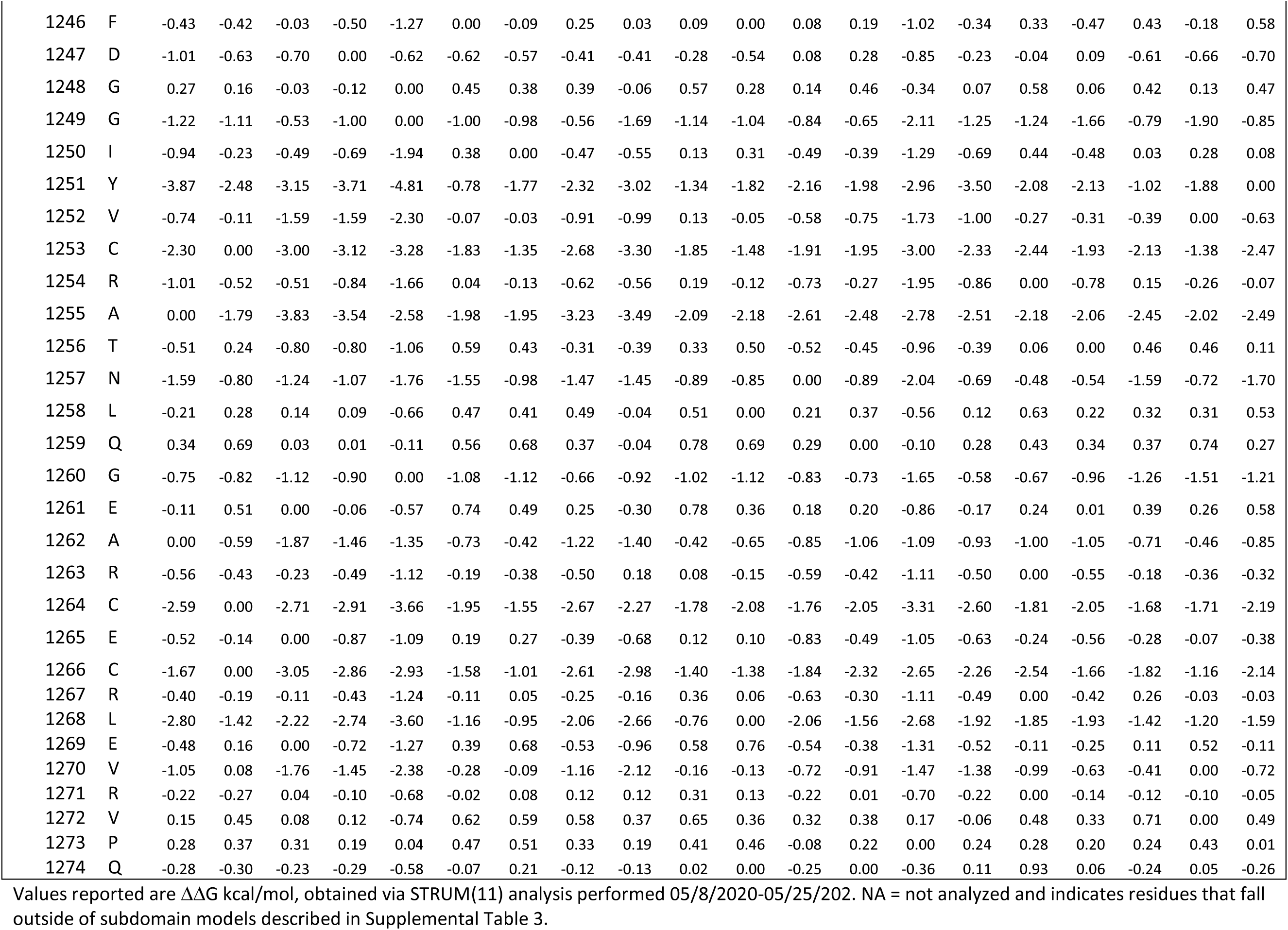
Complete STRUM analysis of *MYBPC*3 Variants

